# Simple synthesis and functionalisation of α-hydroxyglycine-containing peptide fragments

**DOI:** 10.64898/2026.05.29.728772

**Authors:** Pooja R. Solanke, Debolina Sarkar, Pranab C. Saha, Michael T. Taylor

## Abstract

We report here a method for the chemical synthesis of Fmocprotected α-hydroxyglyine (α-OH-Gly) dipeptides. Our method features operational simplicity and is compatible with protecting groups for peptide synthesis. Utility is then demonstrated through substitution at the α-OH-Gly position to yield myriad non-natural amino acid-containing dipeptide fragments.

---

α-hydroxy-glycine (α-OH-Gly) amino acid residues are naturally occurring intermediates in C-terminal amide synthesis^1^ and are useful intermediates in the chemical synthesis of peptides and amide-containing small molecules. Owing to the electronically sensitive hemiaminal functional group, α-OH-Gly residues typically requires N-terminal carbonyl substitution to maintain stability^2^, and thus synthetic strategies for incorporating α-OH-Gly into peptides reflect this requirement. Oxidative strategies have been used to synthesize α-OH-Gly-containing structures both in biosynthesis^1^ and in organic synthesis (**Figure 1A**). With respect to synthetic strategies, Steglich showed that α-acetoxyglycine and α-hydroxy-glycine peptides could be accessed by a transposition of serine (Ser) and threonine (Thr) sidechain *via* oxidative fragmentation using Pb(OAc)_4_^3^. A recent example, such as light-mediated alkyne difunctionalisation of aromatic alkynes with aromatic amides, show elegant new synthetic disconnects but are limited to aromatic-substitute reactants^4^.

**Figure 1.**
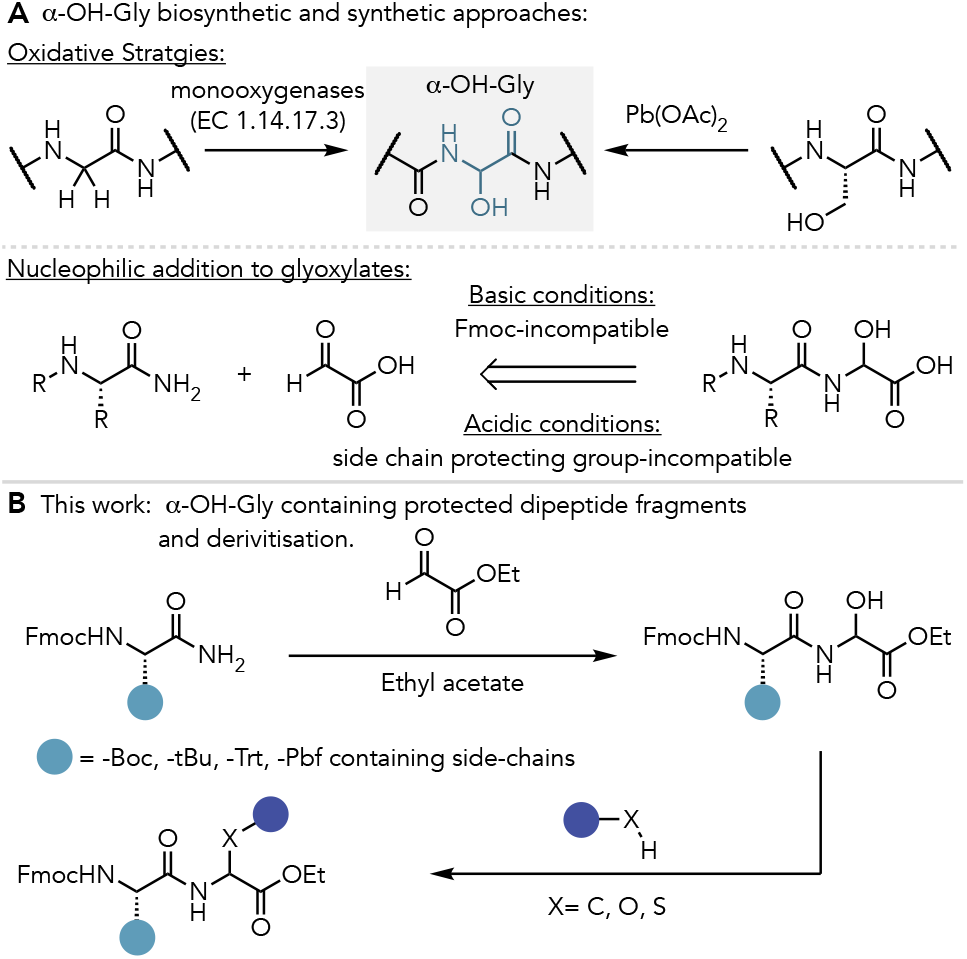
(**A**) Synthetic and biosynthetic routes to α-OH-Gly – containing peptides (**B**) This work: neutral conditions enable the synthesis of α-OH-Gly containing dipeptide fragments possessing relevant protecting groups for peptide synthesis as well as their subsequent derivatisation.

A simple and appealing approach to accessing α-OH-Gly-containing peptides consists of nucleophilic addition of an amide to a glyoxylate equivalent (**Figure 1A**). Multiple approaches to this synthesis have been reported, however, amides are modest nucleophiles and thus many of these approaches require acidic or basic additives. In 2000, Kraus showed that N-terminal Boc-protected amino amides formed unstable mixtures of α-OH-Gly adducts with methyl glyoxylate using amine bases that were then stabilized upon silyl ether the α-OH-Gly alcohol^5^. Acid-mediated conditions have also been reported for α-OH-Gly synthesis either through external acid sources or by use of stoichiometric glyoxylic acid^6^. Whilst these methods can generate α-OH-Gly containing molecules in good yields, the requirement of either acid or base can present challenges with respect to commonly used protecting groups for both the N-terminus (base-sensitive Fmoc groups) and amino acid side chains (acid-sensitive groups). Thus, there are fewer examples of α-OH-Gly synthesis using amino amide nucleophiles that feature fully protected N-termini and side chains that are needed for complex peptide synthesis^7^. Gilvarg and Schacht reported neutral synthesis conditions using CH_2_Cl_2_ or refluxing acetone and glyoxylate esters as coupling partner^2,8^. Here, we present a mild synthetic approach to α-OH-Gly containing dipeptides through nucleophilic addition of amides to glyoxylate esters that is fully compatible with sensitive protecting groups and then present opportunities for subsequent “on peptide” derivatisation of the α-OH-Gly functional group (**Figure 1B**).

To achieve this, we first sought to identify neutral conditions for the nucleophilic addition of Fmoc-Ala-NH_2_ (**1a**) to ethyl glyoxylate under neutral conditions to form (α-OH-Gly)-containing dipeptide analogue **2a** (**Table 1**). Screening across solvents with heating to the solvent’s reflux temperature revealed a strong dependency on solvent for reaction efficiency. For example, DMF, Toluene, THF and 1,2-dichloroethane failed to give yields beyond 10% (entries 1-4), whilst nitromethane, 1,4-dioxane, and dichloromethane gave yields ranging from 14-27% (entries 5-7). Marked increases in yields were observed using acetone (50%, entry 8), acetonitrile (88%, entry 9)) and ethyl acetate (>95%, 77% isolated yield, entry 10). We also observed a strong dependence on temperature (20% in ethyl acetate at RT, entry 11) and diminished yield under acidic conditions (54% yield, entry 12). We decided upon refluxing ethyl acetate as our optimal conditions (entry 10), and these conditions are notable for their simplicity and mildness with the added benefit of use of a green solvent^9^.

**Table 1.** Reaction Optimisation.

| Entry | Solvent | T (°C) | Additive | %Yield<sup>a</sup> (%IY) |
| --- | --- | --- | --- | --- |
| 1 | DMF | 110 | — | 0 |
| 2 | Toluene | 110 | — | 0 |
| 3 | THF | 54 | — | 8 |
| 4 | 1,2-dichloroethane | 72 | — | 10 |
| 5 | CH<sub>3</sub>NO<sub>2</sub> | 82 | — | 14 |
| 6 | 1,4-dioxane | 110 | — | 14 |
| 7 | CH<sub>2</sub>Cl<sub>2</sub> | 40 | — | 27 |
| 8 | Acetone | 58 | — | 50<sup>b</sup> |
| 9 | CH<sub>3</sub>CN | 82 | — | 88 |
| 10 | EtOAc | 82 | — | >95 (77) |
| 11 | EtOAc | 25 | — | 20 |
| 12 | EtOAc | 82 | AcOH (1 eq.) | 54 |

<sup>a</sup> <sup>1</sup>H-NMR yields using 1,1,2,2-tetrachloroethane as an internal standard. <sup>b</sup> dimethyl sulfone used as internal standard. IY= isolated yield.

With optimal conditions in hand, we then proceeded to explore α-OH-Gly dipeptide ester synthesis starting from commercially available amino acids that possess the standard Fmoc N-terminal and sidechain protecting groups used for solid-phase peptide synthesis (SPPS). Fmoc-protected amino acids were first converted to their corresponding amides (**1a-q, Figure 2**, see supplementary information Figure S1 for a full list of aminoamides) by adapting the method of Sureshbabu using ethyl chloroformate and ammonium hydroxide and N-methylmorpholine in THF^10^. The Fmoc-protected amino amides were then reacted with ethyl glyoxylate in ethyl acetate at 82°C for 18-30 hours and α-OH-Gly dipeptide esters (**2a-q**) were then isolated by precipitation with ethyl acetate/hexane and, when necessary, further purified through flash column chromatography with results summarized in **Figure 2**.

**Figure 2.**
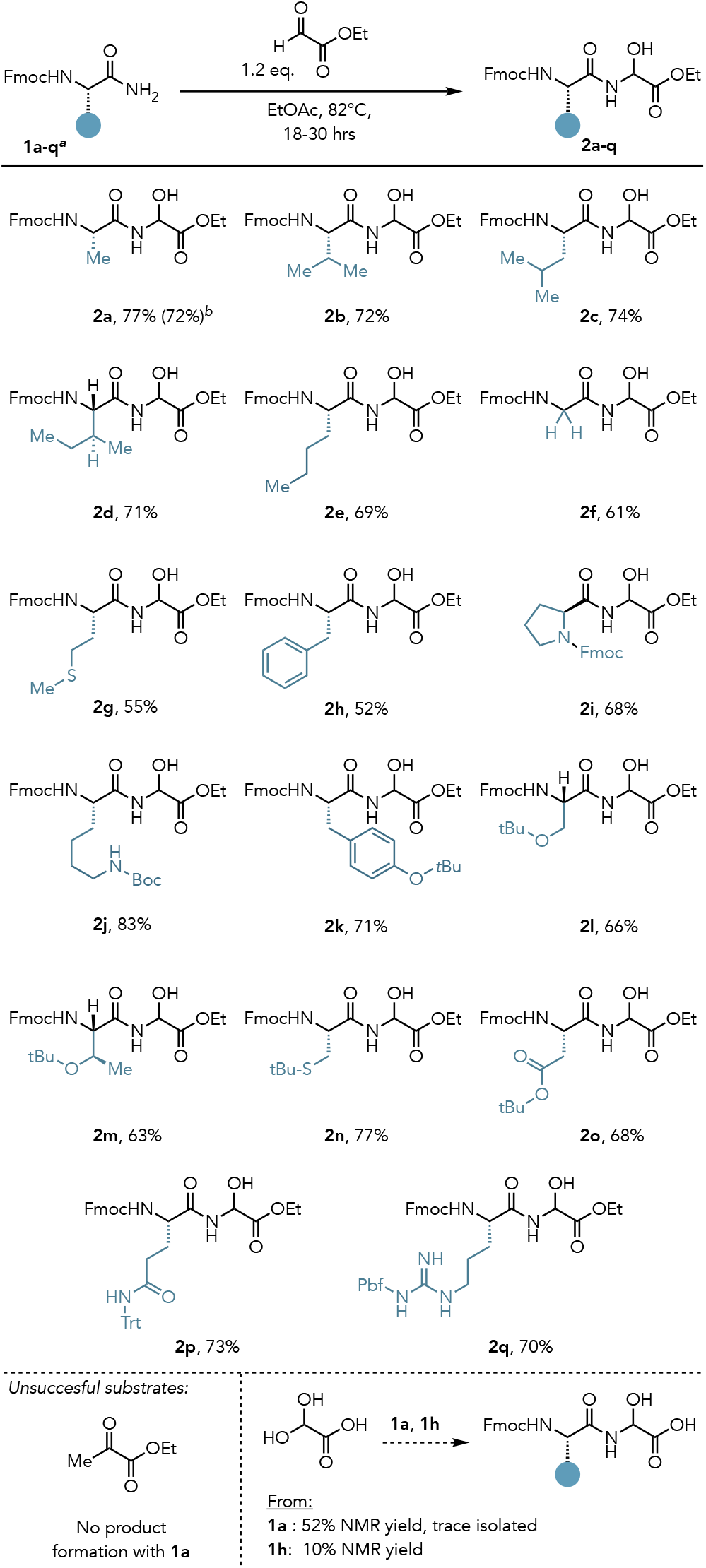
α-OH-Gly containing dipeptide fragment synthesis: protected amino acid substrate scope that shows compatibility with key side chain functional groups and both sidechain and N-terminal protecting groups. ^a^**1a-q** were synthesized by adapting conditions from ref. 10 (see SI for details). ^b^1 gram (3.22 mmol) scale

Alkyl side chain-containing amino acids were well tolerated with this method, forming α-OH-Gly containing dipeptide esters **2a**-**2e** in 69-77% yield. We additionally found that the addition step scales readily, with **2a** being isolated in 72% yield on a 1-gram (3.22 mmol) scale reaction. Other Fmoc amino amides not requiring side chain protection, such as Fmoc-Gly-NH_2_ (**1f**), Fmoc-Phe-NH_2_ (**1g**), and Fmoc-Met-NH_2_ (**1h**) are also well tolerated, forming **2f, 2g**, and **2h** in 61%, 55%, and 52% yields respectively. In **2g**, the oxidizable thioether side chain of Met was found to be well-tolerated. Finally, Fmoc-protected proline amide analogue **1i** was also tolerated, forming **2i** in 68% yield.

Outside of these substrates, the other amino acids possess reactive groups on their sidechains that are masked by protecting groups that are removed using Brønsted acids and are thus labile under the acidic synthesis conditions used in many α-OH-Gly syntheses^6^. However, using the conditions developed here, these groups appear to be well tolerated. For example, Fmoc-Lys(Boc)-NH_2_ **1j** was smoothly converted to **2j** in 83% yield. Substrates containing tert-butyl (*t*Bu) ether sidechain protecting groups such as tyrosine (**2k**, 71%), serine (**2l**, 66%), and threonine (**2m**, 63%) were well tolerated, as were *t*Bu-thioethers such as cysteine (**2n**, 77%) and *t*Bu-esters such as aspartate (**2o**, 68%). The trityl (Trt) protected side chain of glutamine (Gln) analogue Fmoc-Gln(Trt)-NH_2_ **1p** readily converted to **2p** in 73% yield. Finally, the benzofuran-sulfonamide (Pbf) protected arginine (Arg) analogue Fmoc-Arg(Pbf)-NH_2_ **1q** was converted to **2q** in 70% yield. Limitations of the procedure include substrate tolerance on the glyoxylate fragment (**Figure 2C**). For example, the reaction of **1a** with glyoxylic acid-hydrate gave a significant 52% yield by NMR but proved unstable towards myriad purification strategies in our hands whilst attempts to couple Fmoc-Phe-NH_2_ (**1g**) with glyoxylic acid-hydrate gave a low 10% ^1^H-NMR yield of coupling. Attempts to use the ketone of ethyl pyruvate as an electrophile resulted in no observable product formation. As with other approaches, no diastereoselectivity is observed in the reaction. Taken collectively, the conditions developed here represent a mild approach to α-OH-Gly incorporation that are fully compatible with the amino acid protecting groups most commonly used for solid-phased peptide synthesis.

α-OH-Gly is a versatile synthetic intermediate as it can dehydrate to form an electrophilic “glycyl-imine” that can subsequently be functionalized with a nucleophile to form a new, non-natural amino acid. Given the rapidly expanding relevance of peptides containing non-natural amino acids to multiple communities^11^, we sought to harness this capability to structurally diversify protected dipeptide fragments by show that productive bond forming reactions could be utilized to access non-natural amino acid side chains by directly manipulating the peptide backbone (**Figure 3**). First, we highlighted C-C bond formation capabilities through nucleophilic addition of arenes under mild acidic conditions (**Figure 3A**). Specifically, dipeptide ester **2a** was treated with indole in refluxing dichloromethane using catalytic (20 mol%) *p*-toluene sulfonic acid (p-TsOH) for 24 hours. This resulted in nucleophilic addition of the indole into the imine with C3 regioselectivity to form 3-indolyl glycine analogue **3a** in 65% yield. Ester hydrolysis under acidic conditions^12^ such as to not perturb the Fmoc protecting group, gave Fmoc-protected dipeptide acid **3a’** in 45% that can be subsequently used for peptide synthesis.

**Figure 3.**
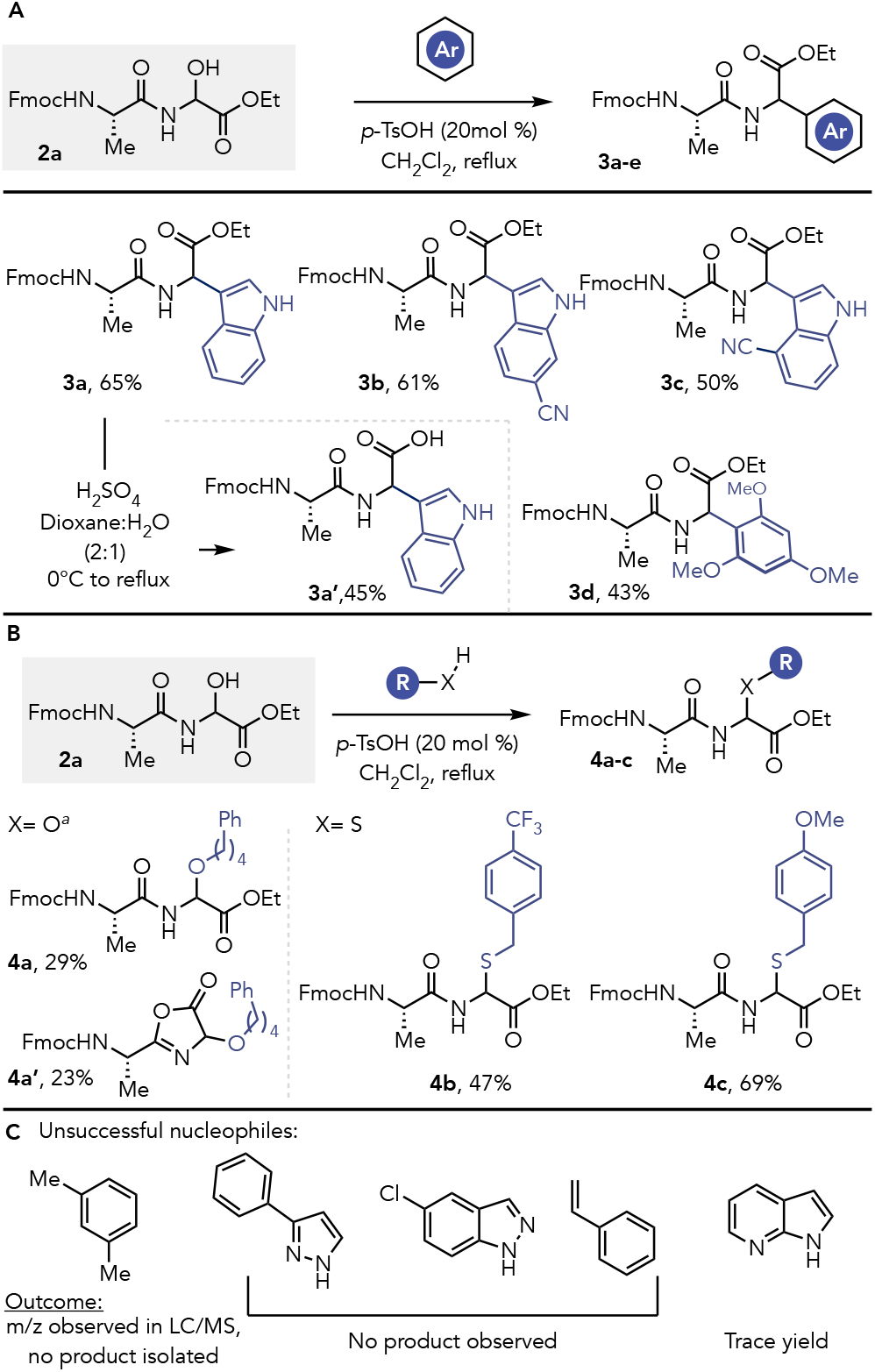
**(A)** Nucleophilic addition with π-based carbon nucleophiles and representative Fmoc-compatible ester hydrolysis to give dipeptide fragments suitable for use in peptide synthesis. **(B)** a-OH-Gly side chain metathesis with representative alcohols and thiols. **(C)** Nucleophiles that either showed low- or no reactivity towards side-chain derivatisation. ^a^1 eq. p-TsOH, performed in neat alcohol^17^.

Next, we further extended the substrate scope by demonstrating that electron-deficient 6-cyanoindole, as well as electron deficient and sterically congested 4-cyanoindole were also competent nucleophiles, yielding 3-indolyl glycine analogues **3b** and **3c** in 61% and 50% yields respectively. Electron-rich 1,3,5-trimethoxybenzene also proved to be a suitable nucleophile, yielding sterically congested aryl-glycine analogue **3d** in a synthetically useful 43% yield. The mild conditions used in this approach enable facile addition to the glycyl-imine intermediate directly on small dipeptide fragments containing protecting groups, which makes this method amenable to rapid structural diversification. Various elegant methods have been developed for substitution/functionalization at the α-position of glycine, including *via* the electrophilic imino-glycine as shown here and several others^5,13^, glycine containing leaving groups in the α-position^14^, enolates^15^, and dehydrogenative couplings^16^. This work complements alternative approaches, such as dehydrogenative couplings that require oxidants, metal catalysts, and sacrificial pre-activating groups requiring aggressive deprotection conditions that could show cross-reactivity with amino acid sidechains and sensitive protecting groups.

Further extending the scope, we show that α-OH-Gly-containing **2a** can be elaborated to a substitute hemiaminal ether using a suitable alcohol, which was exemplified using 1-phenyl-butane-4-ol under acidic conditions^17^ to gives a mix **4a** and oxazolinone in 52% combined yield (**Figure 3B**). Similar transformations using thiols proceeded smoothly to afford thioaminals **4b** and **4c** in 47% and 69% yields respectively. Our derivatisation attempts were not without limitations (**Figure 3C**). For example, nucleophilic addition of meta-xylene proceeded modestly, and whilst an m/z value matching the desired product was observed by LC/MS, we were unable to isolate the desired product. Use of 3-phenylpyrazole, 5-chloroindazole, as well as styrene, also did not yield any desired products whilst 7-azaindole gave trace yield. Taken together, these examples highlight the synthetic versatility of the α-OH-Gly and, when incorporated into peptide fragments, highlights the potential to rapidly generate molecular complexity on peptide fragments directly.

In conclusion, we have developed a simple and mild synthesis of α-OH-Gly containing dipeptide esters. Key features of this method include mild conditions and operational simplicity, which made our synthetic conditions compatible with the key protecting groups used in Fmoc-based peptide synthesis. We then demonstrated that α-OH-Gly dipeptides could undergo subsequent derivatisation reactions, allowing facile access to structurally diverse dipeptides from a common intermediate and yielding dipeptides with protecting group configurations suitable for polypeptide synthesis. Given the growing relevance of accessing structurally diverse peptides in therapeutic development and chemical biology, this method may aid in facilitating access to diverse new analogues.

## Supporting information

Experimental Procedures

## Conflicts of interest

There are no conflicts to declare.

## Data availability

Data for this article, including experimental procedures and spectroscopic characterization of all new compounds, is included in the supplementary information document associated with this manuscript.

## Acknowledgements

We thank R35 GM143120 for support for this work. We also acknowledge the following core facilities for instrumentation support: W.M. Keck Center for Nano-Scale Imaging (RRID:SCR_022884) and the Nuclear Magnetic Resonance Facility (RRID:SCR_012716).

## Conflicts of interest

There are no conflicts to declare.

## Notes

### Competing Interest Statement

The authors have declared no competing interest.

