## Supplementary material for "Simple synthesis and functionalisation of α-hydroxyglycine-containing peptide fragments": Experimental Procedures

Pooja R. Solanke, Debolina Sarkar, Pranab C. Saha, and Michael T. Taylor*
Department of Chemistry & Biochemistry, University of Arizona, Tucson, AZ 85721, United States

*Contact

**Table of Contents**

1. General Information……………………………………………………………………..............3

2. Reaction Optimisation…………………………………………………………………...…4

3. Experimental and Characterization of Compounds……………………….…………..…….5-23

2.1. General Procedure for synthesis of amino amides ………………………....…................5

2.2. General Procedure for synthesis of $\alpha$-OH-Gly derivatives ……..........................................6

2.3. Experimental and Characterization of $\alpha$-OH-Gly derivatives.......................................6-22

4. Product Derivatization…………………………………………………….………………..…23-31

5. NMR Spectra of Compounds………………………...………...……………………...……..32-84

6 References…….……………………………………………………….………......……....85

### 1. General Information

All chemicals have been purchased from commercial sources and were used without further purification unless otherwise noted. All solvents are reagent grade or HPLC grade. The synthetic transformations have been monitored by analytical thin layer chromatography (TLC). TLC was performed using Merck pre-coated glass backed silica gel plates (TLC Silica gel 60 F254). TLC plates were visualized using either UV-light (254 nm), phosphomolybdic acid (PMA) or ceric ammonium molybdate (CAM) staining solutions. Concentration under reduced pressure was performed by rotary evaporation below 48 °C. Flash column chromatography was performed under positive pressure of compressed air using SiliCycle SiliaFlash® P60 silica gel (230-400 mesh). Nuclear magnetic resonance (NMR) spectra were acquired at ambient temperature unless otherwise stated using either a Bruker Avance III 400 or a NEO 500 at the University of Arizona Nuclear Magnetic Resonance Facility. Chemical shifts (δ) are reported in ppm and coupling constants (J) are reported in Hz. Data are reported in the following format: Chemical shift (multiplicity, coupling constants, number of protons). The following convention is used to report multiplicity: s = singlet, d = doublet, t = triplet, q = quartet, qn = quintet, sext = sextet, sept = septet, m = multiplet, br = broad. High-resolution mass spectra were acquired at the analytical & biological mass spectrometry facility at the University of Arizona.

### 2. Optimization of α-OH-Gly containing dipeptide fragments.

To a stirring solution of **1a** (1 eq.) in the solvent listed was added ethyl glyoxylate (1.2 eq.) and any additives used were stirred at the conditions listed for the given time. Reaction solutions were then spiked with an internal standard (1,1,2,2-tetrachloroethane for entries 1-7, and 9-12, and dimethyl sulfone for entry 8) and analysed directly by $^1$H-NMR.

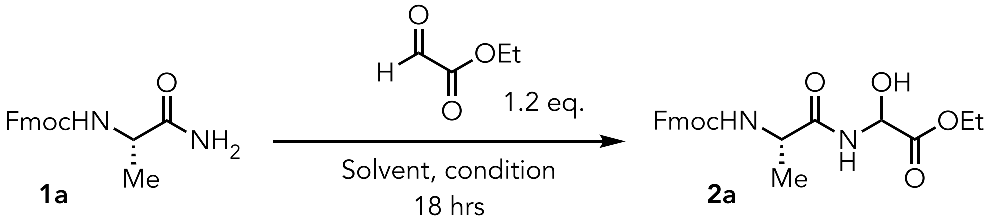


| Entry | Solvent | T (°C) | Additive | %Yield |
|---|---|---|---|---|
| 1 | DMF | 110 | — | 0 |
| 2 | Toluene | 110 | — | 0 |
| 3 | THF | 54 | — | 8 |
| 4 | 1,2-dichloroethane | 72 | — | 10 |
| 5 | $CH_3NO_2$ | 82 | — | 14 |
| 6 | 1,4-dioxane | 110 | — | 14 |
| 7 | $CH_2Cl_2$ | 40 | — | 27 |
| 8 | Acetone | 58 | — | 50 |
| 9 | $CH_3CN$ | 82 | — | 88 |
| 10 | EtOAc | 82 | — | >95 |
| 11 | EtOAc | 25 | — | 20 |
| 12 | EtOAc | 82 | AcOH (1 eq.) | 54 |

**Table S1.** Optimisation of synthetic conditions for **2a**.

### 3. Experimental and Characterization of Compounds

#### 3.1. General Procedure for Synthesis of Amino Amides 1a-1q

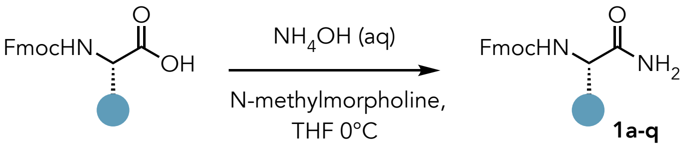


The synthesis of amino amides **1a-1q** was achieved by adapting a previously reported procedure as follows[1]: To a solution of N-fmoc-amino acid (1 mmol) in THF at 0 °C, N-methylmorpholine (NMM) (1.8 mmol), Ethyl Chloroformate (1.5 mmol) were added and stirred at the same temperature for 60 min. After the complete conversion of starting material into mixed anhydride (by TLC), Aq. ammonia (2 mL) was added, and the reaction continued for further 20 min. After the completion of the reaction THF was evaporated using rotary evaporator, and the residue was washed with $NaHCO_3$ (aq), water, and hexanes sequentially, then filtered and dried to afford corresponding amides.

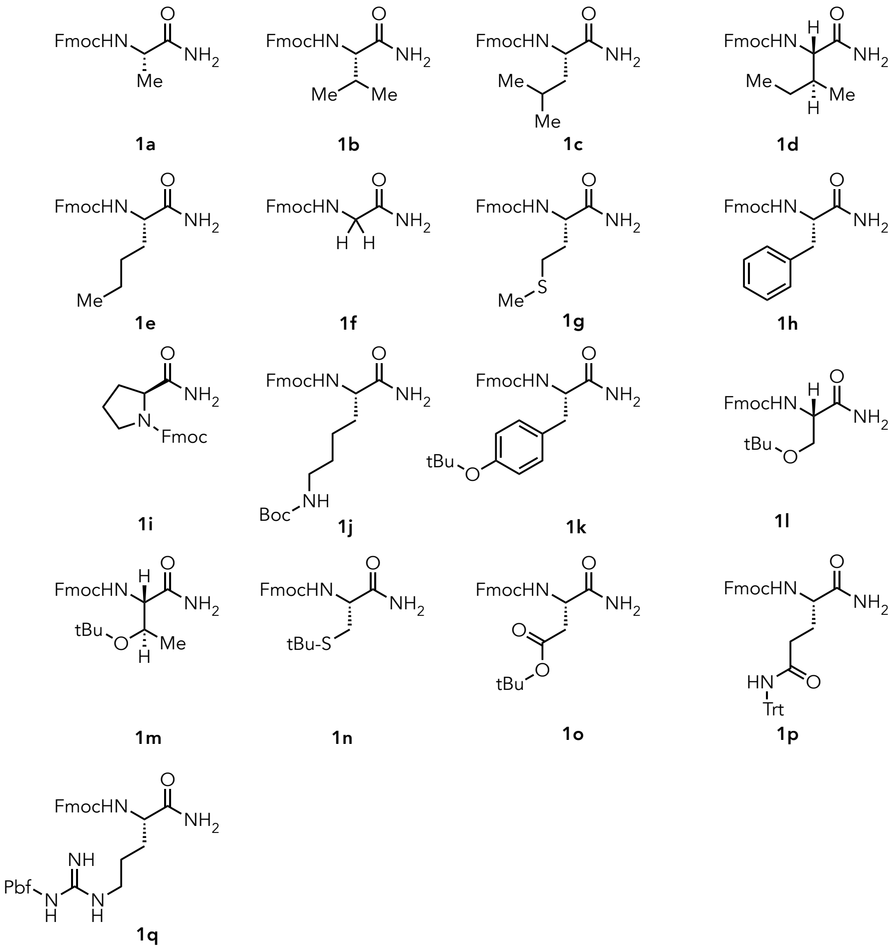


**Figure S1**. Amino amide precursors synthesized for this work.

#### 3.2. General Procedure A: Synthesis of α-OH-Gly dipeptide fragments

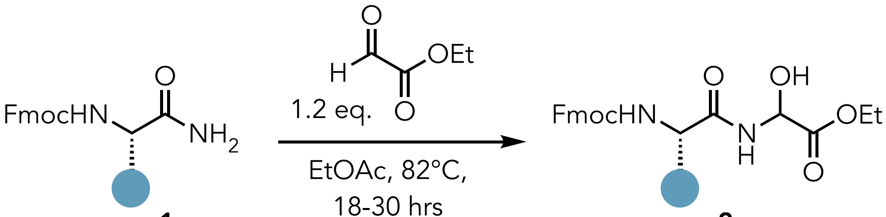


A heat-dried round-bottom flask equipped with a stir bar, a reflux condenser, and a sidearm inlet adapter was charged with the desired protected amino amide (1 mmol, 1.0 eq.) and the setup was evacuated and backfilled with $N_2$. To the flask was added sequentially dry ethyl acetate and ethyl 2-oxoacetate (50 % in toluene solution, 1.2 mmol, 1.2 eq.), and the resulting reaction mixture was stirred at reflux until consumption of the starting material was observed by TLC (18 to 30 hrs). After the reaction was complete and the resulting mixture was cooled to room temperature, and the volatiles were removed using rotary evaporator. The residue was dissolved in ethyl acetate and precipitated with hexanes. The resulting solid was washed with hexanes and then filtered and dried under vacuum.

#### 3.3. Experimental and Characterization of α-OH-Gly dipeptide fragments

**Ethyl 2-((S)-2-((((9H-fluoren-9-yl) methoxy)carbonyl)amino)propanamido)-2 hydroxyacetate (2a)**

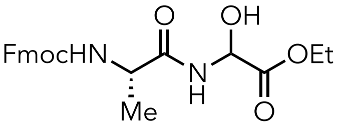


**2a**, 77% (gram scale: 72%)

Compound **2a** was synthesized using General Procedure A as follows: the reaction was performed with **2a** (100 mg, 0.323mmol) and ethyl 2-oxoacetate (80 μL, 0.39 mmol) in ethyl acetate (1mL) were added and allowed to stir at 82 °C until complete conversion of starting material (by TLC analysis). After the reaction was complete and the resulting mixture was cooled to room temperature and the volatiles were removed using rotary evaporator. The residue was dissolved in ethyl acetate and precipitated with hexanes. The resulting solid was washed with hexane and then filtered and dried under vacuum to afford **2a** (103 mg, 77%) as a white solid.

A larger scale reaction was performed using **2a** (1.00 g, 3.22mol), ethyl 2-oxoacetate (790 µL, 3.87 mmol) in 10 mL ethyl acetate and stirring for 24 hours followed by identical reaction workup afforded **2b** (957 mg, 72%).

**$^{1}$H NMR (400 MHz, DMSO)** δ 8.72 (d, *J* = 8.0 Hz, 1H), 7.89 (d, *J* = 7.5 Hz, 2H), 7.73 (t, *J* = 6.6 Hz, 2H), 7.54 (d, *J* = 5.6 Hz, 1H), 7.42 (t, *J* = 7.3 Hz, 2H), 7.33 (t, *J* = 7.3 Hz, 2H), 6.62 (dd, *J* = 32.0, 6.7 Hz, 1H), 5.43 (d, *J* = 7.1 Hz, 1H), 4.23 (d, *J* = 9.6 Hz, 3H), 4.16 – 4.02 (m, 3H), 1.26 – 1.12 (m, 6H).

**$^{13}$C NMR (151 MHz, DMSO)** δ 173.0, 170.2, 156.1, 144.4, 144.3, 141.2, 128.1, 127.6, 125.8, 125.7, 120.6, 71.8, 66.1, 61.3, 61.2, 50.2, 47.1, 18.5, 14.4.

**FT-IR** $\nu_{\max}$ (cm$^{-1}$) 3304, 2981, 1738, 1692, 1660, 1534, 1451, 1370, 1346, 1325, 1247, 1154, 1117, 1101, 1085, 1057, 974, 862, 737.

**HRMS (ESI)** m/z calculated for $[M+Na]^+$ for $C_{22}H_{24}N_2O_6Na$ 435.153208, found 435.15266.

**MP:** 124-125 °C

**Ethyl 2-((S)-2-((((9H-fluoren-9-yl)methoxy)carbonyl)amino)-3-methylbutanamido)-2-hydroxyacetate (2b)**

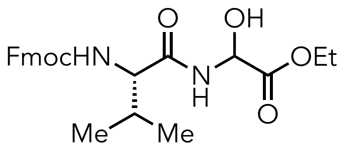


**2b**, 72%

Compound **2b** was synthesized using General Procedure A as follows: the reaction was performed with **1b** (101 mg, 0.298 mmol) and ethyl 2-oxoacetate (72.6 µL, 0.358 mmol) in ethyl acetate (1 mL) were added and allowed to stir at 82 °C until complete conversion of starting material (by TLC analysis). After 30h the reaction was complete and the resulting mixture was cooled to room temperature, and the volatiles were removed using rotary evaporator. The residue was dissolved in ethyl acetate and precipitated with hexanes. The resulting solid was washed with hexanes and then filtered and dried under vacuum to afford **2b** (95 mg, 72 %) as white solid.

**$^{1}$H NMR (500 MHz, Acetone)** δ 8.63 (d, *J* = 8.3 Hz, 1H), 8.31 (d, *J* = 7.6 Hz, 2H), 8.17 (t, *J* = 7.9 Hz, 2H), 7.86 (t, *J* = 7.5 Hz, 2H), 7.78 (t, *J* = 8.0 Hz, 2H), 7.03 (t, *J* = 10.2 Hz, 1H), 6.11 – 6.01 (m, 1H), 5.98 – 5.90 (m, 1H), 4.85 – 4.53 (m, 6H), 2.63 – 2.55 (m, 1H), 1.71 – 1.61 (m, 3H), 1.46 – 1.36 (m, 6H).

$^{13}$**C NMR (126 MHz, Acetone)** δ 170.3, 157.1, 144.9, 144.8, 141.9, 128.4, 127.8, 127.8, 126.0, 120.6, 72.4, 72.3, 67.1, 61.9, 61.8, 60.8, 47.8, 31.7, 31.6, 19.5, 18.0, 17.9, 14.2.

**HRMS (ESI)** m/z calculated for $[M+Na]^+$ for $C_{24}H_{28}N_2O_6Na$ 463.184508, found 463.18396

**FT-IR** $\nu_{max}$ ($cm^{-1}$) 3300, 2958, 1744, 1689, 1657, 1534, 1468, 1451, 1435, 1388, 1370, 1339, 1293, 1246, 1158, 1102, 1081, 1063, 1029, 935, 864, 755, 740.

**MP:** 129-132 °C

**Ethyl 2-((S)-2-((((9H-fluoren-9-yl)methoxy)carbonyl)amino)-4-methylpentanamido)-2-hydroxyacetate (2c)**

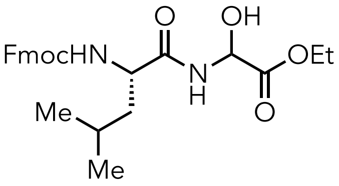


**2c**, 74%

Compound **2c** was synthesized using General Procedure A as follows: the reaction was performed with **1c** (100 mg, 0.284 mmol) and ethyl 2-oxoacetate (69.4 µL, 0.341 mmol) in ethyl acetate (1 mL) were added and allowed to stir at 82 °C until complete conversion of starting material (by TLC analysis). After 22h the reaction was complete and the resulting mixture was cooled to room temperature, and the volatiles were removed using rotary evaporator. The residue was dissolved in ethyl acetate and precipitated with hexanes. The resulting solid was washed with hexanes and then filtered and dried under vacuum to afford **2c** (95 mg, 74 %) as white solid.

$^{1}$**H NMR (500 MHz, $CDCl_3$)** δ 7.74 (d, *J* = 7.5 Hz, 2H), 7.55 (d, *J* = 6.9 Hz, 2H), 7.38 (t, *J* = 7.4 Hz, 2H), 7.29 (d, *J* = 7.4 Hz, 2H), 5.56 – 5.43 (m, 1H), 5.19 (dd, *J* = 16.4, 8.2 Hz, 1H), 4.45 -4.15 (m, 7H), 1.64 (s, 1H), 1.51 (s, 1H), 1.26 (d, *J* = 3.6 Hz, 3H), 0.91 (d, *J* = 6.4 Hz, 6H).

$^{13}$**C NMR (126 MHz, Acetone)** δ 173.5, 173.4, 170.5, 157.1, 145.1, 144.9, 142.1, 128.6, 128.0, 128.0, 126.2, 126.2, 120.8, 120.8, 72.7, 67.2, 67.22 62.1, 62.0, 54.3, 54.3, 48.0, 42.0, 41.9, 25.4, 23.6, 23.5, 21.9, 21.9, 14.4.

**HRMS (ESI)** m/z calculated for $[M+Na]^+$ for $C_{25}H_{30}N_2O_6Na$ 477.200158, found 477.19961.

**FT-IR** $\nu_{max}$ ($cm^{-1}$) 3299, 3062, 2961, 2870, 1755, 1698, 1670, 1536, 1467, 1455, 1372, 1321, 1263, 1241, 1103, 1085, 1038, 992, 858, 755, 742.

**MP:** 144-146 °C

**Ethyl 2-((2S,3S)-2-((((9H-fluoren-9-yl)methoxy)carbonyl)amino)-3-methylpentanamido)-2-hydroxyacetate (2d)**

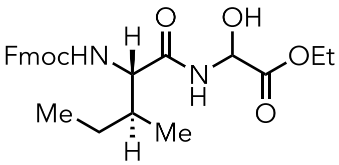

**2d**, 71%

Compound **2d** was synthesized using General Procedure A as follows: the reaction was performed with **1d** (101 mg, 0.287 mmol) and ethyl 2-oxoacetate (70.1 µL, 0.344 mmol) in ethyl acetate (1mL) were added and allowed to stir at 82 ˚C until complete conversion of starting material (by TLC analysis). After 20h the reaction was complete and the resulting mixture was cooled to room temperature and the volatiles were removed using rotary evaporator. The residue was dissolved in ethyl acetate and precipitated with hexanes. The resulting solid was washed with hexanes and then filtered and dried under vacuum to afford **2d** (92 mg, 71 %) as a white solid.

**$^1$H NMR (500 MHz, DMSO)** δ 8.81 (dd, *J* = 38.8, 8.2 Hz, 1H), 7.88 (d, *J* = 7.5 Hz, 2H), 7.78 – 7.68 (m, 2H), 7.46 – 7.38 (m, 3H), 7.35 – 7.29 (m, 2H), 6.70 – 6.58 (m, 1H), 5.49 – 5.38 (m, 1H), 4.32 – 4.18 (m, 3H), 4.15 – 4.04 (m, 3H), 3.94 (t, *J* = 8.5 Hz, 1H), 1.78 – 1.66 (m, 1H), 1.47 – 1.37 (m, 1H), 1.22 – 1.07 (m, 5H), 0.82 (dt, *J* = 15.3, 6.2 Hz, 6H).

**$^{13}$C NMR (126 MHz, DMSO)** δ 171.6, 171.5, 170.7, 169.9, 169.8, 156.1, 156.1, 144.0, 143.9, 140.8, 127.8, 127.2, 127.2, 125.5, 125.5, 120.2, 86.9, 71.3, 71.3, 65.8, 60.9, 60.9, 60.3, 59.1, 58.9, 46.8, 36.7, 36.6, 24.4, f24.4, 15.3, 15.3, 14.1, 14.0, 14.0, 11.0.

**HRMS (ESI)** m/z calculated for $[M+Na]^+$ for $C_{25}H_{31}N_2O_6$ 455.218213, found 455.21766.

**FT-IR** $\nu_{max}$ ($cm^{-1}$) 3450, 3322, 3062, 2965, 1729, 1707, 1671, 1525, 1382, 1366, 1258, 1040, 745.

**MP:** 142-145 °C

**Ethyl 2-((S)-2-((((9H-fluoren-9-yl)methoxy)carbonyl)amino)pentanamido)-2-hydroxyacetate (2e)**

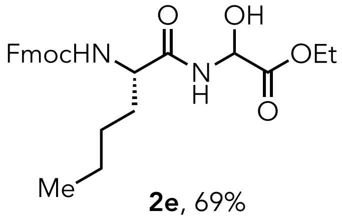

**2e**, 69%

Compound **2e** was synthesized using General Procedure A as follows: the reaction was performed with **1e** (100 mg, 0.284 mmol) and ethyl 2-oxoacetate (69.4 µL, 0.341 mmol) in ethyl acetate (1 mL) were added and allowed to stir at 82 °C until complete conversion of starting material (by TLC analysis). After 18 h the reaction was complete and the resulting mixture was cooled to room temperature and the volatiles were removed using rotary evaporator. The residue was dissolved in ethyl acetate and precipitated with hexanes. The resulting solid was washed with hexanes and then filtered and dried under vacuum to afford **2e** (89 mg, 69 %) as a white solid.

**1H NMR (400 MHz, CDCl3)** δ 7.76 (d, J = 7.4 Hz, 2H), 7.57 (d, J = 6.8 Hz, 2H), 7.40 (t, J = 7.3 Hz, 2H), 7.31 (t, J = 7.0 Hz, 2H), 5.52 (dd, J = 30.4, 7.0 Hz, 1H), 5.36 – 5.26 (m, 1H), 4.49 – 4.38 (m, 1H), 4.31 – 4.14 (m, 3H), 1.91 – 1.79 (m, 1H), 1.68 – 1.57 (m, 1H), 1.40 – 1.19 (m, 6H), 0.96 – 0.84 (m, 3H).

**$^{13}$C NMR (126 MHz, DMSO)** δ 172.3, 172.2, 169.9, 169.8, 156.0, 144.0, 143.8, 140.8, 127.8, 127.2, 125.4, 120.2, 71.4, 71.3, 65., 60.9, 60.9, 54.5, 54.5, 46.8, 31.8, 31.7, 27.6, 27.6, 22.0, 14.1, 14.0, 14.0.

**HRMS (ESI)** m/z calculated for $[M+Na]^+$ for $C_{25}H_{30}N_2O_6Na$ 477.200158, found 477.19961.

**FT-IR** $\nu_{max}$ (cm$^{-1}$) 3452, 3322, 3062, 2958, 2934, 1729, 1708, 1672, 1526, 1458, 1368, 1258, 1040, 745.

**MP:** 109-110 °C

**Ethyl 2-(2-((((9H-fluoren-9-yl)methoxy)carbonyl)amino)acetamido)-2-hydroxyacetate (2f)**

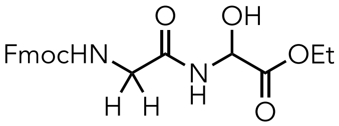


**2f**, 61%

Compound **2f** was synthesized using General Procedure A as follows: the reaction was performed with **1f** (296 mg, 1.00 mmol) and ethyl 2-oxoacetate (245 μL, 1.20 mmol) in ethyl acetate (2.0 mL) were added and allowed to stir at 82 °C until complete conversion of starting material (by TLC analysis). After the reaction was complete and the resulting mixture was cooled to room temperature and the volatiles were removed using rotary evaporator. The residue was purified by silica gel column chromatography (60% EtOAc/hexanes) to afford **2f** (243 mg, 61%) as a white solid.

**$^1$H NMR (400 MHz, $CDCl_3$)** δ 7.74 (d, $J$ = 7.5 Hz, 2H), 7.57 (d, $J$ = 7.3 Hz, 2H), 7.38 (t, $J$ = 7.5 Hz, 2H), 7.31 – 7.25 (m, 2H), 5.77 (s, 1H), 5.60 (d, $J$ = 7.8 Hz, 1H), 4.38 (d, $J$ = 7.1 Hz, 1H), 4.31 – 4.10 (m, 4H), 4.01 – 3.84 (m, 2H), 1.33 – 1.19 (m, 3H).

**$^{13}$C NMR (126 MHz, $CDCl_3$)** δ 170.2, 170.0, 169.2, 165.5, 156.7, 143.8, 141.5, 127.9, 127.3, 125.1, 120.2, 72.2, 67.5, 63.0, 47.2, 44.6, 14.2, 13.5.

**HRMS (ESI)** m/z calculated for $[M+Na]^+$ for $C_{21}H_{23}N_2O_6$ 399.155613, found 399.15506.

**FT-IR** $\nu_{max}$ ($cm^{-1}$) 3458, 3322, 3062, 2980, 1731, 1709, 1676, 1532, 1448, 1258, 1042, 745.

**MP:** 116-118 °C

**Ethyl 2-((S)-2-((((9H-fluoren-9-yl)methoxy)carbonyl)amino)-4-(methylthio)butanamido)-2-hydroxyacetate (2g)**

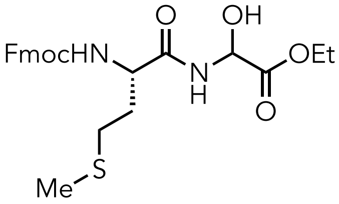

**2g**, 55%

The synthesis of the compound **2g** by using General Procedure A as follows: the reaction was performed with **1g** (370 mg, 1.00 mmol) and ethyl 2-oxoacetate (245 µL, 1.20 mmol) in ethyl acetate (2.0 mL) were added and allowed to stir at 82 °C until complete conversion of starting material (by TLC analysis). After the reaction was complete and the resulting mixture was cooled to room temperature and the volatiles were removed using rotary evaporator. The residue was dissolved in ethyl acetate and precipitated with hexanes. The resulting solid was washed with ether and hexane and then filtered and dried under vacuum to afford **2g** (260 mg, 55 %) as a white solid.

**$^1$H NMR (500 MHz, Acetone)** δ 8.06 (d, *J* = 8.1 Hz, 1H), 7.75 (d, *J* = 7.6 Hz, 2H), 7.63 – 7.59 (m, 2H), 7.30 (t, *J* = 7.4 Hz, 2H), 7.22 (t, *J* = 7.5 Hz, 2H), 6.70 (d, *J* = 8.1 Hz, 1H), 5.53 – 5.38 (m, 2H), 4.29 – 4.18 (m, 3H), 4.13 (t, *J* = 7.1 Hz, 1H), 4.09 – 4.02 (m, 2H), 2.51 – 2.42 (m, 2H), 2.03 – 1.96 (m, 4H), 1.89 – 1.81 (m, 1H), 1.12 (t, *J* = 7.1 Hz, 3H).

**$^{13}$C NMR (126 MHz, Acetone)** δ 172.3, 170.5, 157.0, 145.1, 145.0, 142.1, 128.6, 128.0, 126.2, 120.8, 120.8, 72.7, 72.6, 67.3, 62.1, 55.0, 48.0, 32.9, 32.9, 30.7, 30.7, 15.2, 14.4.

**HRMS (ESI)** m/z calculated for $[M+Na]^+$ for $C_{24}H_{28}N_2O_6SNa$ 495.156579,

**FT-IR** $\nu_{max}$ ($cm^{-1}$) 3450, 3322, 3062, 2980, 2934, 1729, 1707, 1672, 1526, 1458, 1368, 1258, 1040, 1022, 702, 745.

**MP:** 136-139 °C

**Ethyl 2-((S)-2-((((9H-fluoren-9-yl)methoxy)carbonyl)amino)-3-phenylpropanamido)-2-hydroxyacetate (2h)**

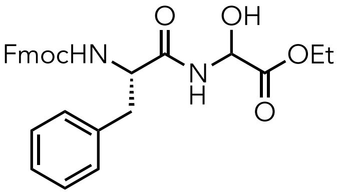

**2h**, 52%

The synthesis of the compound **2h** by using General Procedure A as follows: the reaction was performed with **1h** (386 mg, 1.00 mmol) and ethyl 2-oxoacetate (245 μL, 1.20mmol) in ethyl acetate (2.0 mL) were added and allowed to stir at 82 °C until complete conversion of starting material (by TLC analysis). After 24h the reaction was complete and the resulting mixture was cooled to room temperature and the volatiles were removed using rotary evaporator. The residue was dissolved in ethyl acetate and precipitated with hexanes. The resulting solid was washed with hexanes and then filtered and dried under vacuum to afford **2h** (255 mg, 52 %) as a white solid

**$^1$H NMR (500 MHz, Acetone)** δ 8.17 – 8.09 (m, 1H), 7.74 (d, *J* = 7.5 Hz, 2H), 7.55 – 7.49 (m, 2H), 7.29 (t, *J* = 7.5 Hz, 2H), 7.25 – 7.12 (m, 6H), 7.11 – 7.06 (m, 1H), 6.65 – 6.58 (m, 1H), 5.55 – 5.41 (m, 2H), 4.44 – 4.38 (m, 2H), 4.22 – 4.13 (m, 1H), 4.10 – 4.01 (m, 5H), 3.15 – 3.07 (m, 1H), 2.87 – 2.81 (m, 1H), 1.16 – 1.09 (m, 3H).

**$^{13}$C NMR (126 MHz, Acetone)** δ 184.5, 171.5, 171.4, 170.5, 169.5, 169.5, 155.9, 155.9, 144.1, 144.1, 144.1, 141.1, 141.1, 137.7, 137.6, 129.42, 129.4, 128.2, 127.6, 127.0, 126.4, 125.3, 125.2, 119.9, 87.4, 87.3, 71.8, 71.7, 66.4, 61.8, 61.2, 60.5, 56.2, 47.0, 37.8, 37.7, 13.5, 13.5, 13.3.

**HRMS (ESI)** m/z calculated for $[M+Na]^+$ for $C_{28}H_{28}N_2O_6Na$ 511.184508, found 511.18396.

**FT-IR** $\nu_{max}$ (cm$^{-1}$) 3301, 3020, 1747, 1693, 1662, 1535, 1477, 1466, 1445, 1370, 1301, 1259, 1214, 1102, 1036, 934, 865, 750, 702.

**MP:** 126-128 °C

**(9H-fluoren-9-yl)methyl (2S)-2-((2-ethoxy-1-hydroxy-2-oxoethyl)carbamoyl)pyrrolidine-1-carboxylate (2i)**

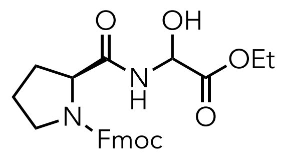


**2i**, 68%

Compound **2i** was synthesized using General Procedure A as follows: the reaction was performed with **1i** (101 mg, 0.299 mmol) and ethyl 2-oxoacetate (73.3 μL, 0.359 mmol) in ethyl acetate (1 mL) were added and allowed to stir at 82 °C until complete conversion of starting material (by TLC analysis). After the reaction was complete and the resulting mixture was cooled to room temperature and the volatiles were removed using rotary evaporator. The resulting residue was purified by silica gel column chromatography (60% EtOAc/hexanes) to afford **2i** (89 mg, 68 %) as a white solid.

**$^1$H NMR (400 MHz, CDCl$_3$)** δ 7.95 (dd, *J* = 18.4, 6.3 Hz, 1H), 7.76 (d, *J* = 7.3 Hz, 2H), 7.58 (d, *J* = 6.9 Hz, 2H), 7.39 (t, *J* = 7.0 Hz, 2H), 7.31 (t, *J* = 7.4 Hz, 2H), 5.60 – 5.34 (m, 1H), 4.39 (d, *J* = 20.3 Hz, 3H), 4.23 (d, *J* = 6.4 Hz, 3H), 3.48 (d, *J* = 44.9 Hz, 2H), 2.18 (dd, *J* = 63.1, 34.5 Hz, 1H), 1.91 (d, *J* = 25.8 Hz, 2H), 1.25 (dd, *J* = 8.9, 5.3 Hz, 3H).

**$^{13}$C NMR (126 MHz, CDCl$_3$)** δ 172.6, 171.4, 171.3, 169.6, 156.1, 156.0, 155.3, 143.9, 143.7, 141.3, 127.8, 127.1, 125.1, 125.1, 120.0, 72.1, 67.8, 62.3, 60.8, 60.5, 47.5, 47.1, 31.2, 28.9, 28.7, 24.5, 23.5, 21.1, 14.2.

**HRMS (ESI)** m/z calculated for $[M+Na]^+$ for $C_{24}H_{26}N_2O_6Na$ 461.168858, found 461.16831.

**FT-IR** $\nu_{max}$ ($cm^{-1}$) 3451, 3062, 2980, 1730, 1708, 1658, 1442, 1258, 1040, 745.

**MP:** 106-110 °C

**Ethyl2-((S)-2-((((9H-fluoren-9-yl)methoxy)carbonyl)amino)-6-((tertbutoxycarbonyl)amino) hexanamido)- 2-hydroxyacetate (2j)**

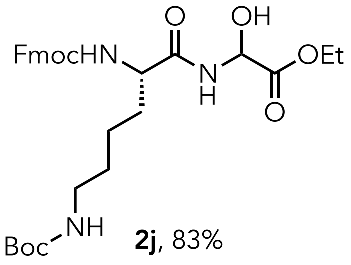


**2j**, 83%

Compound **2j** was synthesized using General Procedure A as follows: the reaction was performed with **1j** (468 mg, 1.00 mmol) and ethyl 2-oxoacetate (245 µL, 1.20 mmol) in ethyl acetate (2.0 mL) were added and allowed to stir at 82 °C until complete conversion of starting material (by TLC analysis). After 20h the reaction was complete, and the resulting mixture was cooled to room temperature and the volatiles were removed using rotary evaporator. The residue was dissolved in ethyl acetate and precipitated with hexanes. The resulting solid was washed with hexanes and then filtered and dried under vacuum to afford **2j** (475 mg, 83 %) as a white solid.

**$^1$H NMR (500 MHz, Acetone)** δ 8.04 (d, *J* = 7.0 Hz, 1H), 7.76 (d, *J* = 7.4 Hz, 2H), 7.61 (t, *J* = 6.6 Hz, 2H), 7.31 (t, *J* = 7.5 Hz, 2H), 7.22 (t, *J* = 7.4 Hz, 2H), 6.62 – 6.55 (m, 1H), 5.86 (s, 0.5H), 5.53 – 5.45 (m, 1H), 5.42 – 5.33 (m, 1H), 5.19 (s, 0.5H), 4.94 (d, *J* = 25.9 Hz, 0.5H), 4.26 – 4.18 (m, 2H), 4.16 – 4.02 (m, 4H), 3.15 (d, *J* = 47.5 Hz, 1H), 3.00 – 2.91 (m, 1H), 1.79 – 1.69 (m, 1H), 1.63 – 1.43 (m, 3H), 1.43 – 1.29 (m, 9H), 1.19 – 1.08 (m, 5H).

**$^{13}$C NMR (126 MHz, $CDCl_3$)** δ 172.7, 172.7, 172.4, 169.4, 169.3, 163.5, 156.3, 143.7, 141.3, 127.7, 127.1, 125.1, 120.0, 72.2, 67.2, 62.6, 47.1, 31.58, 29.5, 28.4, 28.3, 22.2, 22.0, 14.1, 14.0.

**HRMS (ESI)** m/z calculated for $[M+Na]^+$ for $C_{30}H_{40}N_3O_8$ 570.281542, found 570.28099

**FT-IR** $\nu_{max}$ ($cm^{-1}$) 3317, 2978, 2937, 1682, 1523, 1478, 1451, 1393, 1367, 1249, 1167, 1103, 862, 759, 741.

**MP:** 87-90°C

**Ethyl 2-((S)-2-((((9H-fluoren-9-yl)methoxy)carbonyl)amino)-3-(4-(tert-butoxy)phenyl)propanamido)-2-hydroxyacetate (2k)**

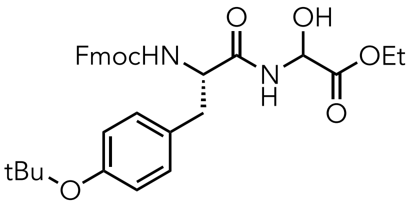


**2k**, 71%

The synthesis of the compound **2k** by using General Procedure A as follows: the reaction was performed with **1k** (100 mg, 0.219 mmol) and ethyl 2-oxoacetate (53.5 μL, 0.263 mmol) in ethyl acetate (0.730 mL) were added and allowed to stir at 82 ˚C until complete conversion of starting material (by TLC analysis). After the reaction was complete and the resulting mixture was cooled to room temperature and the volatiles were removed using rotary evaporator. The residue was dissolved in ethyl acetate and precipitated with hexanes. The resulting solid was washed with hexanes and then filtered and dried under vacuum to afford **2k** (87 mg, 71 %) as a white solid.

**$^1$H NMR (400 MHz, $CDCl_3$)** δ 7.78 (d, *J* = 7.5 Hz, 2H), 7.56 (d, *J* = 7.4 Hz, 2H), 7.42 (t, *J* = 7.4 Hz, 2H), 7.33 (t, *J* = 7.4 Hz, 2H), 7.14 – 7.05 (m, 2H), 6.97 – 6.90 (m, 2H), 5.52 – 5.32 (m, 2H), 4.49 – 4.33 (m, 3H), 4.31 – 4.17 (m, 3H), 3.06 (s, 2H), 1.46 – 1.19 (m, 12H).

$^{13}$C NMR (126 MHz, $CDCl_3$) δ 172.9, 171.5, 169.2, 143.6, 141.3, 127.8, 127.1, 125.0, 120.1, 82.3, 72.3, 67.4, 62.8, 5.22, 47.1, 28.1, 28.0, 14.0, 14.0.

**HRMS (ESI)** m/z calculated for $[M+Na]^+$ for $C_{32}H_{36}N_2O_7Na$ 583.242023, found 583.24147.

**FT-IR** $\nu_{max}$ ($cm^{-1}$) 3311, 3065, 2977, 1670, 1609, 1507, 1478, 1451, 1390, 1365, 1237, 1162, 1105, 1047, 924, 898, 855, 759, 741.

**MP:** 80-82 °C

**Ethyl 2-((S)-2-((((9H-fluoren-9-yl)methoxy)carbonyl)amino)-3-(tert-butoxy)propanamido)-2-hydroxyacetate (2l)**

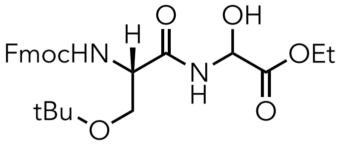


**2l**, 66%

Compound **2l** was synthesized using General Procedure A as follows: the reaction was performed with **1l** (382 mg, 1.00 mmol) and ethyl 2-oxoacetate (245 μL, 1.20 mmol) in ethyl acetate (2.0 ml) were added and allowed to stir at 82 ˚C until complete conversion of starting material (by TLC analysis). After 18h the reaction was complete, and the resulting mixture was cooled to room temperature and the volatiles were removed using rotary evaporator. The residue was dissolved in ethyl acetate and precipitated with hexanes. The resulting solid was washed with hexane then filtered and dried under vacuum to afford **2l** (322 mg, 66 %) as a white solid.

**$^1$H NMR (400 MHz, CDCl$_3$)** δ 7.95 (s, 1H), 7.77 (d, *J* = 7.5 Hz, 2H), 7.60 (d, *J* = 7.4 Hz, 2H), 7.41 (t, *J* = 7.4 Hz, 2H), 7.32 (td, *J* = 7.4, 1.0 Hz, 2H), 5.71 (d, *J* = 4.2 Hz, 1H), 5.58 (d, *J* = 7.4 Hz, 1H), 4.42 (d, *J* = 7.0 Hz, 2H), 4.36 – 4.19 (m, 4H), 3.88 – 3.76 (m, 1H), 3.39 (t, *J* = 8.5 Hz, 1H), 1.33 (t, *J* = 7.1 Hz, 3H), 1.23 (s, 9H).

**$^{13}$C NMR (126 MHz, CDCl$_3$)** δ 171.6, 169.3, 156.2, 143.9, 143.8, 141.4, 130.8, 129.3, 127.9, 127.2, 125.2, 120.2, 118.6, 74.8, 72.4, 72.4, 67.3, 62.8, 61.6, 54.3, 47.2, 27.5, 14.2.

**HRMS (ESI)** m/z calculated for $[M+Na]^+$ for $C_{26}H_{32}N_2O_7Na$ 507.210723, found 507.21017.

**FT-IR** *ν*max (cm$^{-1}$) 3452, 3323, 3062, 2978, 1729, 1708, 1673, 1526, 1366, 1252, 1110, 852, 745.

**MP:** 156-159°C

**Ethyl 2-((2S,3R)-2-((((9H-fluoren-9-yl)methoxy)carbonyl)amino)-3-(tert-butoxy)butanamido)-2-hydroxyacetate (2m)**

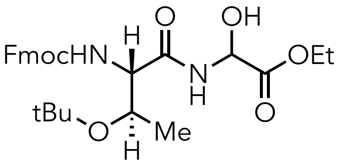


**2m**, 63%

Compound **2m** was synthesized using General Procedure A as follows: the reaction was performed with **1m** (398 mg, 1.00 mmol) and ethyl 2-oxoacetate (245 µL, 1.20 mmol) in ethyl acetate (2.0 mL) added and allowed to stir at 82 °C until complete conversion of starting material (by TLC analysis). After the reaction was complete and the resulting mixture was cooled to room temperature and the volatiles were removed using rotary evaporator. The residue was dissolved in ethyl acetate and precipitated with hexanes. The resulting solid was washed with hexanes and then filtered and dried under vacuum to afford **2m** (316 mg, 63 %) as a white solid.

**$^{1}$H NMR (500 MHz, $CDCl_3$)** δ 8.41 – 8.22 (m, 1H), 7.76 (d, $J$ = 1 Hz, 2H), 7.60 (d, $J$ = 7.0 Hz, 2H), 7.40 (t, $J$ = 6.9 Hz, 2H), 7.31 (t, $J$ = 7.1 Hz, 2H), 5.97 (d, $J$ = 4.4 Hz, 1H), 5.54 (dd, $J$ = 30.6, 17.0 Hz, 1H), 4.39 (d, $J$ = 6.9 Hz, 2H), 4.26 (dd, $J$ = 45.3, 18.3 Hz, 6H), 1.41 – 1.21 (m, 12H), 1.13 – 1.03 (m, 3H).

**$^{13}$C NMR** (126 MHz, $CDCl_3$) δ 170.8, 170.6, 169.5, 169.4, 156.1, 144.0, 143.7, 141.4, 141.4, 127.8, 127.2, 125.2, 120.1, 120.1, 76.0, 75.9, 72.3, 72.1, 67.1, 66.7, 62.8, 62.7, 58.7, 58.6, 47.2, 28.2, 16.9, 16.7, 14.2.

**HRMS (ESI)** m/z calculated for $[M+Na]^+$ for $C_{27}H_{35}N_2O_7$ 499.244427, found 499.24388.

**FT-IR** $\nu_{max}$ ($cm^{-1}$) 3316, 3068, 2979, 1726, 1679, 1526, 1478, 1451, 1413, 1393, 1368, 1249, 1154, 1105, 1083, 1051, 959, 845, 759, 741.

**MP:** 158-161°Cs

**Ethyl 2-((R)-2-((((9H-fluoren-9-yl)methoxy)carbonyl)amino)-3-(tert-butylthio)propanamido)-2-hydroxyacetate (2n)**

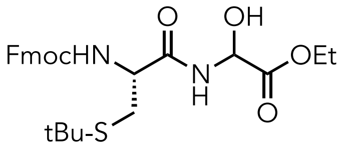


**2n**, 77%

Compound **2n** was synthesized using General Procedure A as follows: the reaction was performed with **1n** (100 mg, 0.251 mmol) and ethyl 2-oxoacetate (61 µL, 0.30 mmol) in ethyl acetate (1 mL) were added and allowed to stir at 82 °C until complete conversion of starting material (by TLC analysis). After 18 h the reaction was complete and the resulting mixture was cooled to room temperature, and the volatiles were removed using rotary evaporator. The residue was dissolved in ethyl acetate and precipitated with hexanes. The resulting solid was washed with hexanes and then filtered and dried under vacuum.to afford **2n** (103 mg, 77%) as a white solid.

**$^1$H NMR (500 MHz, Acetone)** δ 8.26 (s, 1H), 7.88 (d, *J* = 7.6 Hz, 2H), 7.75 (d, *J* = 7.5 Hz, 2H), 7.49 – 7.33 (m, 4H), 6.82 (t, *J* = 9.0 Hz, 1H), 5.68 – 5.54 (m, 2H), 4.45 – 4.36 (m, 2H), 4.34 - 4.25 (m, 2H), 4.24 – 4.15 (m, 2H), 3.09 – 3.03 (m, 1H), 2.91 – 2.86 (m, 1H), 1.34 (d, *J* = 3.6 Hz, 9H), 1.28 - 1.24 (m, 3H).

**$^{13}$C NMR (126 MHz, Acetone)** δ 171.3, 170.3, 170.3, 156.9, 156.9, 145.0, 142.1, 128.6, 128.0, 126.3, 126.2, 120.8, 72.8, 72.7, 67.5, 62.1, 56.2, 47.9, 42.8, 31.4, 31.2, 14.4, 14.4.

**HRMS (ESI)** m/z calculated for $[M+Na]^+$ for $C_{26}H_{32}N_2O_6SNa$ 523.187880, found 523.18733.

**FT-IR** $\nu_{max}$ (cm$^{-1}$) 3317, 3064, 2961, 1738, 1669, 1527, 1451, 1366, 1322, 1246, 1161, 1103, 1049, 862, 759, 741, 702.

**MP:** 135-139°C

**Tert-butyl (3S)-3-((((9H-fluoren-9-yl)methoxy)carbonyl)amino)-4-((2-ethoxy-1-hydroxy-2-oxoethyl)amino)-4-oxobutanoate (2o)**

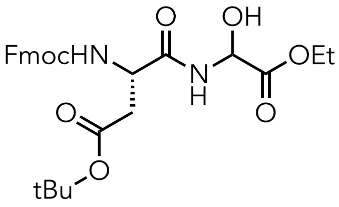


**2o**, 68%

Compound **2o** was synthesized using General Procedure A as follows: the reaction was performed with **1o** (100 mg, 0.247 mmol) and ethyl 2-oxoacetate (60 µL, 0.36 mmol) in ethyl acetate 1 mL) were added and allowed to stir at 82 ˚C until complete conversion of starting material (by TLC analysis). After 20h the reaction was complete and the resulting mixture was cooled to room temperature, and the volatiles were removed using rotary evaporator. The residue was dissolved in ethyl acetate and precipitated with hexanes. The resulting solid was washed with hexanes and then filtered and dried under vacuum to afford **2o** (82 mg, 68 %) as a white solid.

**$^{1}$H NMR (500 MHz, CDCl$_3$)** δ 7.76 (d, *J* = 7.5 Hz, 2H), 7.72 – 7.68 (m, 1H), 7.58 (d, *J* = 7.3 Hz, 2H), 7.40 (t, *J* = 7.4 Hz, 2H), 7.31 (td, *J* = 7.4, 1.7 Hz, 2H), 6.04 – 5.95 (m, 1H), 5.55 – 5.47 (m, 1H), 4.61 – 4.39 (m, 3H), 4.34 – 4.21 (m, 4H), 2.96 – 2.87 (m, 1H), 2.67 – 2.56 (m, 1H), 1.45 (s, 9H), 1.30 (t, *J* = 7.1 Hz, 3H).

**$^{13}$C NMR (126 MHz, CDCl$_3$)** δ 173.0, 171.5, 171.0, 169.4, 156.3, 156.2, 143.8, 143.7, 141.5, 141.4, 127.9, 127.2, 125.2, 125.1, 120.2, 120.2, 82.3, 82.3, 82.2, 72.4, 67.5, 67.2, 62.8, 62.8, 51.1, 50.9, 47.3, 47.2, 37.3, 37.3, 28.2, 28.1, 14.2.

**HRMS (ESI)** m/z calculated for $[M+Na]^+$ for $C_{27}H_{32}N_2O_8Na$ 535.205638, found 535.20509.

**FT-IR** $\nu_{max}$ (cm$^{-1}$) 3316, 3068, 2979, 1726, 1679, 1526, 1478, 1451, 1413, 1393, 1368, 1249, 1154, 1105, 1083, 1051, 959, 845, 759, 741.

**MP:** 133-135 °C

**Ethyl 2-((S)-2-((((9H-fluoren-9-yl)methoxy)carbonyl)amino)-5-oxo-5-(tritylamino)pentanamido)-2-hydroxyacetate (2p)**

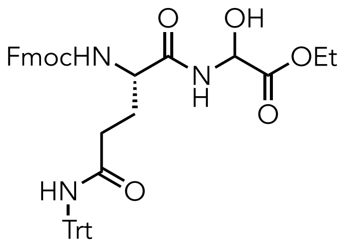


**2p**, 73%

Compound **2p** was synthesized using General Procedure A as follows: the reaction was performed with **1p** (100 mg, 0.164 mmol) and ethyl 2-oxoacetate (50 μL, 0.25 mmol) in ethyl acetate (1 mL) were added and allowed to stir at 82 °C until complete conversion of starting material (by TLC analysis). After the reaction was complete and the resulting mixture was cooled to room temperature, and the volatiles were removed using rotary evaporator. The residue was then dissolved in ethyl acetate and precipitated with hexanes. The resulting solid was washed with hexane then filtered and dried under vacuum to afford **2p** (89 mg, 73%) as a white solid.

**$^1$H NMR (400 MHz, CDCl$_3$)** δ 7.75 (d, *J* = 7.5 Hz, 2H), 7.58 (d, *J* = 7.4 Hz, 2H), 7.38 (d, *J* = 7.4 Hz, 2H), 7.32 – 7.26 (m, 7H), 7.23 (dd, *J* = 9.5, 3.5 Hz, 8H), 7.07 – 6.89 (m, 1H), 6.48 (s, 1H), 5.94 (d, *J* = 25.7 Hz, 1H), 5.31 (d, *J* = 7.6 Hz, 1H), 4.46 – 4.33 (m, 2H), 4.25 – 4.00 (m, 3H), 2.69 – 2.37 (m,2 H), 2.18 – 1.90 (m, 3H), 1.32 – 1.21 (m, 3H).

**$^{13}$C NMR (126 MHz, CDCl$_3$)** δ 153.3, 144.5, 14397, 141.4, 128.8, 128.2, 127.9, 127.3, 125.3, 120.1, 73.0, 71.0, 67.1, 62.8, 53.5, 53.5, 47.3, 47.3, 30.2, 14.2, 14.2.

**HRMS (ESI)** m/z calculated for [M+Na]$^+$ for $C_{43}H_{42}N_3O_7$ 712.302277, found 712.30173.

**FT-IR** $\nu_{max}$ (cm$^{-1}$) 3310, 3014, 1669, 1490, 1447, 1215, 1049, 902, 749, 699.

**MP:** 128-130 °C

**Ethyl 2-((S)-2-((((9H-fluoren-9-yl)methoxy)carbonyl)amino)-5-(3-((2,2,4,6,7-pentamethyl-2,3-dihydrobenzofuran-5-yl)sulfonyl)guanidino)pentanamido)-2-hydroxyacetate (2q)**

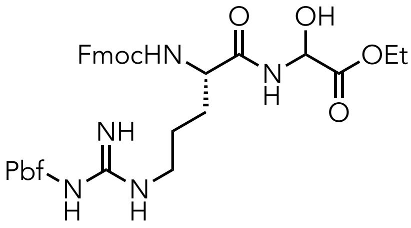


**2q**, 70%

Compound **2q** was synthesized using General Procedure A as follows: the reaction was performed with **1q** (200 mg, 0.309 mmol) and ethyl 2-oxoacetate (76 µL, 0.37 mmol) in ethyl acetate (1 mL) added and allowed to stir at 82 °C until complete conversion of starting material (by TLC analysis). After the reaction was complete and the resulting mixture was cooled to room temperature, and the volatiles were removed using rotary evaporator. The residue was purified by silica gel column chromatography (60% EtOAc/hexanes) to afford **2q** (157 mg, 70%) as a white solid.

**$^1$H NMR (400 MHz, $CDCl_3$)** δ 8.40 - 8.32 (s, 1H), 7.71 (d, J = 7.5 Hz, 2H), 7.57 (bs, 2H), 7.37 – 7.30 (bs, 2H), 7.25 – 7.16 (bs, 2H), 6.47 – 6.26 (m, 2H), 5.55 (bs, 1H), 4.39 – 4.11 (m, 5H), 3.33 – 3.17 (m, 1H), 2.89 (s, 2H), 2.56 (s, 3H), 2.48 (s, 3H), 2.06 (s, 3H), 1.96 – 1.57 (m, 4H), 1.42 (s, 6H), 1.24 – 1.13 (m, 3H).

**$^{13}$C NMR (151 MHz, $CDCl_3$)** δ 170.0, 156.7, 156.6, 144.0, 143.8, 141.3, 127.8, 127.2, 125.4, 125.0, 120.0, 117.9, 86.7, 72.2, 67.3, 62.6, 62.5, 47.2, 43.3, 29.8, 28.7, 19.5, 18.1, 14.1, 12.6.

**HRMS (ESI)** m/z calculated for $[M+H]^+$ for $C_{38}H_{49}N_5O_9H$ 750.317277, found 750.31673.

**FT-IR** $\nu_{max}$ (cm$^{-1}$) 3450, 3330, 3062, 2980, 1729, 1708, 1673, 1630, 1527, 1500, 1451, 1340, 1258, 1158, 745.

**MP:** 152-153°C

### 4. Product derivatization

#### 3.1. General Procedure B: α-OH-Gly dipeptide fragment derivatization

A heat-dried screw capped vial equipped with a stir bar was charged with Ethyl 2-((S)-2-((((9H-fluoren-9-yl)methoxy)carbonyl)amino)propanamido)-2 hydroxyacetate (**2a**) (1.0 eq.). To the vial was sequentially added dry $CH_2Cl_2$, a respective nucleophile (1.5 eq.) and pTSA (0.2 eq). The resulting reaction mixture was stirred at reflux until consumption of the starting material was observed by TLC. After the reaction was complete and the resulting mixture was cooled to room temperature, and the volatiles were removed using rotary evaporator. The residue was purified by column chromatography to obtain the product.

**Ethyl 2-((S)-2-((((9H-fluoren-9-yl)methoxy)carbonyl)amino)propanamido)-2-(1H-indol-3-yl)acetate (3a)**

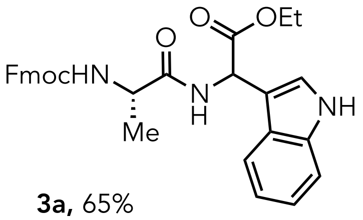


**3a,** 65%

Compound **3a** was synthesized using General Procedure B as follows: a heat-dried screw capped vial equipped with a stir bar was charged with Ethyl 2-((S)-2-((((9H-fluoren-9-yl)methoxy)carbonyl)amino)propanamido)-2 hydroxyacetate **2a** (300 mg, 0.728 mmol) and sequentially added dry $CH_2Cl_2$ and 1H-indole (127 mg, 1.09 mmol) and pTSA (27 mg, 0.14 mmol). The reaction mixture was stirred at reflux until consumption of the starting material was observed by TLC. After 24 h the reaction was complete and the resulting mixture was cooled to room temperature and the volatiles were removed using rotary evaporation. The residue was purified by silica gel column chromatography (50% EtOAc/hexanes) to afford **3a** (242 mg, 65 %).

**$^1$H NMR (400 MHz, $CDCl_3$)** δ 8.21 (bs, 1H), 7.75 (d, *J* = 7.4 Hz, 2H), 7.67 (d, *J* = 7.8 Hz, 1H), 7.59 - 7.50 (m, 2H), 7.39 (t, *J* = 7.4 Hz, 2H), 7.32 – 7.27 (m, 2H), 7.21 – 7.07 (m, 3H), 5.81 (d, *J* = 6.9 Hz, 1H), 5.47 (bs, 1H), 4.40 – 4.04 (m, 6H), 1.71 (bs, 1H), 1.32 - 1.47 (m, 3H), 1.23 - 1.15 (m, 3H).

**$^{13}$C NMR (126 MHz, $CDCl_3$)** δ 172.4, 172.4, 171.4, 171.3, 171.0, 156.0, 143.9, 143.7, 141.2, 136.3, 127.7, 127.1, 125.4, 125.4, 125.1, 123.8, 123.6, 122.5, 120.1, 120.0, 119.0, 118.9, 111.6, 110.2, 67.0, 61.8, 61.7, 60.5, 50.4, 50.2 47.0, 21.1, 19.0 18.9, 14.2, 14.0, 14.0.

**HRMS (ESI)** m/z calculated for $[M+Na]^+$ for $C_{30}H_{29}N_3O_5Na$ 534.200491, found 534.19994.

**FT-IR** $\nu_{max}$ (cm$^{-1}$) 3310, 3063, 2981, 1706, 1663, 1516, 1478, 1451, 1369, 1339, 1248, 1189, 1106, 1078, 1032, 759, 741.
**MP:** 107-109 °C

**2-((*S*)-2-((((9*H*-fluoren-9-yl)methoxy)carbonyl)amino)propanamido)-2-(1*H*-indol-3-yl)acetic acid (3a')**

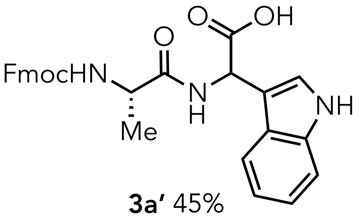


**3a'** 45%

The synthesis of the compound **3a'** by adapting a reported procedure[2] as follows: a heat-dried round-bottom flask equipped with a magnetic stir bar, reflux condenser, and sidearm inlet adapter was charged with compound **3a** (65 mg, 0.13 mmol) and dissolved in a 2:1 mixture of 1,4-dioxane and water (1.5 mL). The solution was cooled to 0 °C, and conc. $H_2SO_4$ (135 uL, 2.6 mmol) was added dropwise with stirring. After stirring at 0 °C for 10 minutes, the reaction mixture was heated to reflux and maintained at that temperature until complete consumption of the starting material was confirmed by TLC (18 h). Upon completion, the mixture was cooled to 0 °C and carefully quenched with $NaHCO_3$(aq). The volatiles were removed under reduced pressure, and the residue was purified by reverse-phased chromatography to afford **3a'** (28 mg, 45%).
Autoflash method: solvents: A: 0.1% TFA in $H_2O$, B: 0.1% TFA in $CH_3CN$; method: hold 10% B for 3 min, 10-45% B over 6 min, hold 45% B for 12 min, 45-90% B over 6 min, hold 90% B for 5 min, 90-10% B over 1 min, hold 10% B for 2 min; UV-Vis: 254, 214 nm, column: RediSep Gold C18Aq 24-30g 100 Å, flow rate: 30 mL/min.
**$^1$H NMR (800 MHz, DMSO)** δ 11.13 (d, *J* = 17.9 Hz, 1H), 8.37 (dd, *J* = 16.9, 7.1 Hz, 1H), 7.88 (d, *J* = 7.4 Hz, 3H), 7.71 (d, *J* = 7.8 Hz, 1H), 7.63 – 7.56 (m, 1H), 7.52 (d, *J* = 8.6 Hz, 1H), 7.42 – 7.30 (m, 4H), 7.10 (d, *J* = 4.2 Hz, 1H), 7.02 – 6.96 (m, 1H), 5.57 – 5.49 (m, 1H), 4.31 – 4.03 (m, 3H), 1.29 – 1.15 (m, 3H).
**$^{13}$C NMR (201 MHz, DMSO)** δ 172.5, 172.2, 172.2, 155.6, 143.9, 143.8, 140.7, 136.2, 136.2, 127.6, 127.1, 125.7, 125.3, 124.4, 124.1, 121.4, 120.1, 119.1, 118.9, 111.7, 111.6, 110.2, 109.7 65.7, 65.60, 49.9, 49.5, 49.5, 49.4, 46.6, 18.6, 18.3.
**FT-IR** $\nu_{max}$ (cm$^{-1}$) 3410, 3060, 2980, 1710,1685,1650, 1450, 1250, 740.
**HRMS (ESI)** m/z calculated for $[M+Na]^+$ for $C_{28}H_{26}N_3O_5$ 484.187247, found 484.18670.

**MP:** 107-110 °C

**Ethyl 2-((S)-2-((((9H-fluoren-9-yl)methoxy)carbonyl)amino)propanamido)-2-(6-cyano-1H-indol-3-yl)acetate (3b)**

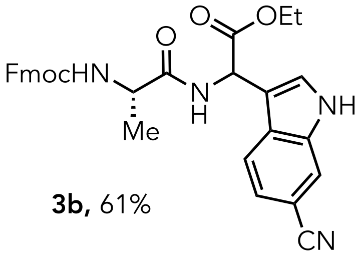


**3b,** 61%

Compound **3b** was synthesized using General Procedure B as follows: a heat-dried screw capped vial equipped with a stir bar was charged with Ethyl 2-((S)-2-((((9H-fluoren-9-yl)methoxy)carbonyl)amino)propanamido)-2 hydroxyacetate **2a** (50 mg, 0.12 mmol) and sequentially added dry $CH_2Cl_2$ and 1H-indole-6-carbonitrile (26 mg, 0.18 mmol) and pTSA (5 mg, 0.02 mmol). The reaction mixture was stirred at reflux until consumption of the starting material was observed by TLC. After 24 h the reaction was complete and the resulting mixture was cooled to room temperature and the volatiles were removed using rotary evaporation. The residue was purified by silica gel column chromatography (50% EtOAc/hexanes) to afford **3b** (40 mg, 61 %).

**$^1$H NMR (500 MHz, DMSO)** δ 11.77 (d, $J$ = 14.5 Hz, 1H), 8.62 (dd, $J$ = 25.7, 6.5 Hz, 1H), 7.92 – 7.84 (m, 3H), 7.76 – 7.66 (m, 4H), 7.56 – 7.51 (m, 1H), 7.44 – 7.26 (m, 6H), 5.61 (dd, $J$ = 22.8, 6.5 Hz, 2H), 4.27 – 4.15 (m, 4H), 4.12 – 4.03 (m, 2H), 1.29 – 1.15 (m, 3H), 1.14 – 1.07 (m, 3H).

**$^{13}$C NMR (126 MHz, $CDCl_3$)** δ 171.6, 171.6, 168.5, 168.5, 156.1, 143.8, 143.8, 141.5, 141.4, 129.5, 129.5, 127.9, 127.2, 125.6, 125.6, 125.6, 125.6, 125.1, 125.1, 123.1, 120.1, 67.4, 62.6, 62.6, 53.7, 50.6, 47.2, 35.1, 34.9, 18.3, 14.0.

**HRMS (ESI)** m/z calculated for $[M+Na]^+$ for $C_{31}H_{28}N_2O_5Na$ 559.195741, found 559.19519.

**FT-IR** $\nu_{max}$ (cm$^{-1}$) 3310, 3063, 2981, 1706, 1663, 1516, 1478, 1451, 1369, 1339, 1248, 1189, 1106, 1078, 1032, 759, 741.

**MP:** 188-189 °C.

**Ethyl 2-((S)-2-((((9H-fluoren-9-yl)methoxy)carbonyl)amino)propanamido)-2-(4-cyano-1H-indol-3-yl)acetate (3c)**

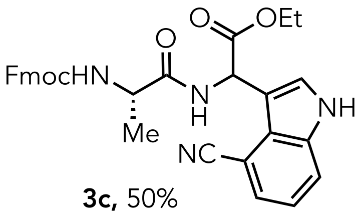

**3c,** 50%

Compound **3c** was synthesized using General Procedure B as follows: a heat-dried screw capped vial equipped with a stir bar was charged with Ethyl 2-((S)-2-((((9H-fluoren-9-yl)methoxy)carbonyl)amino)propanamido)-2 hydroxyacetate **2a** (50 mg, 0.12 mmol) and sequentially added dry $CH_2Cl_2$ and 1H-indole-4-carbonitrile (26 mg, 0.18 mmol) and pTSA (5 mg, 0.02 mmol). The reaction mixture was stirred at reflux until consumption of the starting material was observed by TLC. After 24 h the reaction was complete and the resulting mixture was cooled to room temperature and the volatiles were removed using rotary evaporation. The residue was purified by silica gel column chromatography (50% EtOAc/hexanes) to afford **3c** (33 mg, 50 %).

**$^1$H NMR (500 MHz, $CDCl_3$) δ** 7.77 (d, *J* = 4.9 Hz, 2H), 7.70 (dd, *J* = 10.3, 5.9 Hz, 1H), 7.63 (dd, *J* = 12.1, 6.0 Hz, 2H), 7.47 (ddd, *J* = 26.0, 14.8, 7.6 Hz, 2H), 7.35 (s, 2H), 7.29 – 7.18 (m, 3H), 6.12 (d, *J* = 20.7 Hz, 1H), 4.36 – 4.10 (m, 6H), 1.41 (t, *J* = 6.6 Hz, 3H), 1.18 (dd, *J* = 12.7, 6.7 Hz, 3H)

**$^{13}$C NMR (126 MHz, Acetone)** δ 173.0, 169.3, 160.0, 159.9, 156.9 145.1, 145.0, 142.1, 131.3, 131.23 130.0, 128.6, 128.1, 128.0, 126.3, 126.2, 126.2, 126.2, 120.9, 114.8, 114.7, 67.4, 62.5, 55.6, 55.5, 54.4, 51.6, 51.4, 48.0, 34.9, 34.7, 18.6, 18.3, 14.4.

**FT-IR** $\nu_{max}$ ($cm^{-1}$) 3306, 3016, 1660, 1509, 1451, 1427, 1345, 1215, 1077, 1021, 739.

**HRMS (ESI)** m/z calculated for $[M+H]^+$ for $C_{31}H_{29}N_4O_5$ 537.213796, found 537.213796.

**MP:** 118-121 °C

**Ethyl 2-((S)-2-((((9H-fluoren-9-yl)methoxy)carbonyl)amino)propanamido)-2-(2,4,6-trimethoxyphenyl)acetate (3d)**

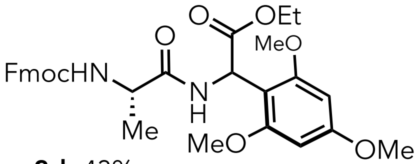


**3d,** 43%

Compound **3d** was synthesized using General Procedure B as follows: a heat-dried screw capped vial equipped with a stir bar was charged with Ethyl 2-((S)-2-((((9H-fluoren-9-yl)methoxy)carbonyl)amino)propanamido)-2 hydroxyacetate **2a** (100 mg, 0.243 mmol) and sequentially added dry $CH_2Cl_2$ and 1,3,5-trimethoxybenzene (61 mg, 0.36 mmol) and pTSA (9 mg, 0.05 mmol). The reaction mixture was stirred at reflux until consumption of the starting material was observed by TLC. After 24 h the reaction was complete and the resulting mixture was cooled to room temperature and the volatiles were removed using rotary evaporation. The residue was purified by silica gel column chromatography (30% EtOAc/hexanes) to afford **3c** (58 mg, 43%).

**$^1$H NMR (400 MHz, $CDCl_3$)** δ 7.78 (t, *J* = 6.2 Hz, 2H), 7.67 – 7.54 (m, 2H), 7.46 – 7.39 (m, 2H), 7.38 – 7.31 (m, 2H), 7.04 – 6.78 (m, 1H), 6.33 – 6.22 (m, 1H), 6.11 (d, *J* = 16.0 Hz, 2H), 5.64 – 5.49 (m, 1H), 4.35 (ddd, *J* = 31.2, 15.1, 8.8 Hz, 3H), 4.25 – 4.08 (m, 4H), 3.87 – 3.74 (m, 9H), 1.54 – 1.36 (m, 3H), 1.17 (dt, *J* = 14.1, 7.1 Hz, 3H).

**$^{13}$C NMR (126 MHz, $CDCl_3$)** δ 171.5 161.5, 158.9, 158.8, 156.0, 144.1, 141.3, 127.8, 127.2, 125.3, 120.0, 106.6, 91.0, 90.9, 67.2, 61.4, 60.5, 56.1, 56.0, 55.4, 50.5, 47.2, 47.2, 46.6, 46.4, 21.2, 19.5, 18.9, 14.3, 14.3, 14.3.

**FT-IR** $\nu_{max}$ ($cm^{-1}$) 3311, 2939, 2840, 1729, 1668, 1609, 1595, 1500, 1451, 1419, 1368, 1322, 1222, 1204, 1151, 1122, 1057, 1033, 951, 867, 815, 757, 741.

**HRMS (ESI)** m/z calculated for $[M+Na]^+$ for $C_{31}H_{35}N_2O_8$ 563.239343, found 563.23879.

**MP:** 88-90 °C

**Ethyl 2-((S)-2-((((9H-fluoren-9-yl)methoxy)carbonyl)amino)propanamido)-2-(4-phenylbutoxy)acetate (4a)**

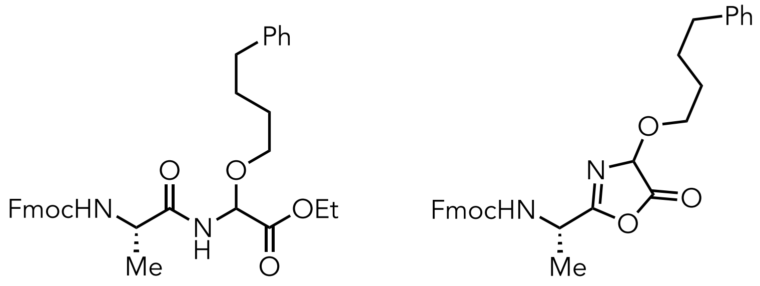

**4a,** 29% **4a',** 23%

The synthesis of the compound **4a** by adapting a reported procedure[3] as follows: a heat-dried screw capped vial equipped with a stir bar was charged with Ethyl 2-((S)-2-((((9H-fluoren-9-yl)methoxy)carbonyl)amino)propanamido)-2 hydroxyacetate **2a** (80 mg, 0.19 mmol) of in the 4-phenylbutan-1-ol (800 µL, 5.2mmol) was added p-TsOH ( 37 mg, 0.19 mmol). The resulting mixture was heated to 85°C with stirring until a clear solution (approximately 10 minutes stirring) was observed and persisted. The resulting homogenous solution was then allowed to cool to room temperature and was quenched by addition of a $NaHCO_3$ (aq) (10 mL). The mixture was extracted with $CH_2Cl_2$ (3 × 20 mL), and the combined organic layers were washed with brine (20 mL), dried over $Na_2SO_4$, filtered, and the solvent was removed using a rotary evaporator. The residual crude product was purified by silica gel column chromatography (30% EtOAc/hexanes) to afford **4a** (31 mg, 29 %) and cyclic oxazolone **4a'** (24 mg, 23%)

**4a** characterisation:

**$^1$H NMR (400 MHz, $CDCl_3$)** δ 7.77 (d, *J* = 7.6 Hz, 2H), 7.58 (d, *J* = 7.1 Hz, 2H), 7.40 (t, *J* = 7.5 Hz, 2H), 7.32 (ddd, *J* = 7.5, 6.6, 2 Hz, 2H), 7.24 (t, *J* = 6.4 Hz, 2H), 7.19 – 7.13 (m, 3H), 6.92 (s, 1H), 5.55 (t, *J* = 8.2 Hz, 1H), 5.27 (s, 1H), 4.40 (d, *J* = 7.0 Hz, 2H), 4.31 – 4.18 (m, 4H), 3.68 – 3.59 (m, 2H), 2.64 – 2.56 (m, 2H), 1.68 – 1.60 (m, 4H), 1.44 – 1.39 (m, 2H), 1.30 – 1.24 (m, 3H).

**$^{13}$C NMR (126 MHz, $CDCl_3$)** δ 168.0, 141.4, 128.5, 128.4, 127.9, 127.2, 125.8, 125.2, 120.2, 69.7, 69.7, 67.3, 62.4, 47.2, 35.7, 31.1, 29.1, 27.9, 14.2.

**HRMS (ESI)** m/z calculated for $[M+Na]^+$ for $C_{30}H_{29}N_3O_5Na$ 435.153200, found 534.19994.

**FT-IR** $\nu_{max}$ ($cm^{-1}$) 3306, 3025, 2937, 1746, 1672, 1604, 1522, 1478, 1451, 1374, 1333, 1248, 1210, 1106, 1077, 859, 759, 741, 700.

**4a'** characterisation

**$^1$H NMR (400 MHz, $CDCl_3$)** δ 7.79 (d, *J* = 7.5 Hz, 2H), 7.60 (d, *J* = 7.3 Hz, 2H), 7.43 (t, *J* = 7.4 Hz, 2H), 7.36 (s, 2H), 7.26 (dd, *J* = 7.5, 5.2 Hz, 3H), 7.22 – 7.14 (m, 6H), 5.58 (d, *J* = 8.9 Hz, 1H),

4.43 (d, *J* = 6.8 Hz, 2H), 4.30 (d, *J* = 6.4 Hz, 1H), 4.26 – 4.18 (m, 3H), 3.70 – 3.59 (m, 2H), 2.62 (d, *J* = 10.6 Hz, 4H), 1.74 – 1.57 (m, 10H), 1.43 (d, *J* = 6.2 Hz, 3H).

**$^{13}$C NMR (126 MHz, $CDCl_3$)** δ 172.7, 168.0, 143.8, 142.4, 141.9, 141.4, 128.5, 128.5, 128.4, 127.9, 127.2, 126.0, 125.9 125.2, 120.2, 69.7, 67.3, 66.2, 50.8, 47.23, 35.7, 35.4, 29.2, 28.1, 27.9, 27.6.

**FT-IR** $\nu_{max}$ (cm$^{-1}$) 3301, 3062, 3025, 2926, 2857, 1745, 1670, 1604, 1517, 1496, 1478, 1451, 1395, 1333, 1246, 1208, 1106, 1076, 939, 758, 740, 699.

**HRMS (ESI)** m/z calculated for $[M+H]^+$ for $C_{30}H_{31}N_2O_5$ 499.223298, found 499.22275.

**Ethyl 2-((S)-2-((((9H-fluoren-9-yl)methoxy)carbonyl)amino)propanamido)-2-((4-(trifluoromethyl)benzyl)thio)acetate (4b)**

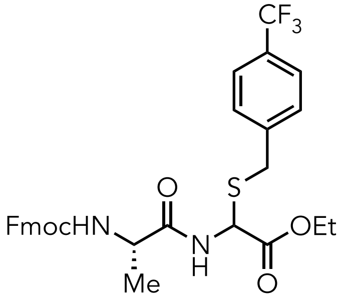


**4b,** 47%

Compound **4b** was synthesized using General Procedure B as follows: a heat-dried screw capped vial equipped with a stir bar was charged with Ethyl 2-((S)-2-((((9H-fluoren-9-yl)methoxy)carbonyl)amino)propanamido)-2 hydroxyacetate **2a** (50 mg, 0.12 mmol) and sequentially added dry $CH_2Cl_2$, (4-(trifluoromethyl)phenyl)methanethiol (35 mg, 0.18 mmol) and pTSA (5 mg, 0.02 mmol). The reaction mixture was stirred at reflux until consumption of the starting material was observed by TLC. After 25 h the reaction was complete and the resulting mixture was cooled to room temperature and the volatiles were removed using a rotary evaporator. The residue was purified by silica gel column chromatography (50% EtOAc/hexanes) to afford **4b** (34 mg, 47%).

**$^1$H NMR (500 MHz, Acetone)** δ 8.13 (d, *J* = 7.6 Hz, 2H), 7.86 (d, *J* = 7.6 Hz, 6H), 7.72 (dd, *J* = 11.1, 4.9 Hz, 6H), 7.41 (t, *J* = 7.5 Hz, 6H), 7.36 – 7.28 (m, 6H), 6.71 (d, *J* = 9.0 Hz, 2H), 5.58 (t, *J* = 8.9 Hz, 3H), 4.40 – 4.34 (m, 3H), 4.34 – 4.21 (m, 10H), 4.20 – 4.11 (m, 6H), 1.69 – 1.56 (m, 9H), 1.22 (t, *J* = 7.1 Hz, 9H).

**$^{13}$C NMR (126 MHz, $CDCl_3$)** δ 171.61, 171.56, 168.54, 168.48, 156.15, 143.81, 143.77, 141.52, 141.41, 129.57, 129.55, 127.90, 127.22, 125.65, 125.62, 125.61, 125.58, 125.15, 125.11, 123.07, 120.15, 67.37, 62.63, 62.61, 53.71, 50.64, 47.17, 35.08, 34.89, 18.34, 14.06.

**$^{19}$F NMR (471 MHz, $CDCl_3$)** δ -62.48, -62.51.

**FT-IR** $\nu_{max}$ ($cm^{-1}$) 3304, 3064, 2981, 1665, 1518, 1449, 1417, 1371, 1324, 1251, 1166, 1123, 1067, 1035, 1019, 856, 758, 741, 701

**HRMS (ESI)** m/z calculated for $[M+Na]^+$ for $C_{30}H_{30}F_3N_2O_5S$ 587.182753, found 587.18220.

**MP:** 105-106°C

**Ethyl 2-((S)-2-((((9H-fluoren-9-yl)methoxy)carbonyl)amino)propanamido)-2-((4-methoxybenzyl)thio)acetate (4c)**

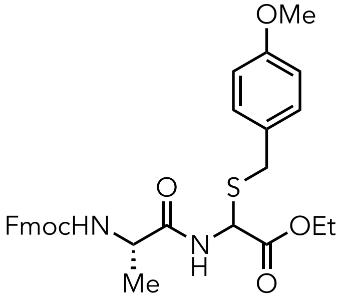


**4c,** 69%

Compound **4c** was synthesized using General Procedure B as follows: a heat-dried screw capped vial equipped with a stir bar was charged with Ethyl 2-((S)-2-((((9H-fluoren-9-yl)methoxy)carbonyl)amino)propanamido)-2 hydroxyacetate **2a** (100 mg, 0.243 mmol) and sequentially added dry $CH_2Cl_2$, (4-methoxyphenyl)methanethiol (52 uL, 0.36 mmol) and pTSA (9mg, 0.05 mmol). The reaction mixture was stirred at reflux until consumption of the starting material was observed by TLC. After 18 h the reaction was complete and the resulting mixture was cooled to room temperature and the volatiles were removed using rotary evaporation. The residue was purified by silica gel column chromatography (50% EtOAc/hexanes) to afford **4c** (93 mg, 69 %).

**$^1$H NMR (400 MHz, Acetone)** δ 7.94 – 7.88 (m, 1H), 7.86 (dd, $J$ = 7.5, 0.8 Hz, 2H), 7.76 – 7.67 (m, 2H), 7.41 (t, $J$ = 7.5 Hz, 2H), 7.36 – 7.29 (m, 2H), 7.26 (dd, $J$ = 8.2, 5.8 Hz, 2H), 6.87 – 6.83 (m, 1H), 6.82 – 6.78 (m, 1H), 5.46 (d, $J$ = 8.7 Hz, 1H), 4.35 (ddd, $J$ = 17.0, 9.8, 5.1 Hz, 3H), 4.29 – 4.23 (m, 1H), 4.14 (dt, $J$ = 11.3, 5.6 Hz, 2H), 3.90 (dd, $J$ = 12.8, 4.4 Hz, 1H), 3.83 (dd, $J$ = 12.8, 3.6 Hz, 1H), 3.74 (d, $J$ = 16.5 Hz, 3H), 1.39 (dd, $J$ = 7.1, 5.6 Hz, 3H), 1.22 (td, $J$ = 7.1, 2.5 Hz, 3H).

**$^{13}$C NMR (126 MHz, Acetone)** δ 173.4, 173.3, 170.6, 157.1, 145.2, 145.01, 142.2, 128.6, 128.0, 128.0, 126.2, 120.9, 120.9, 72.7, 67.2, 62.1, 54.3, 54.3, 48.1, 42.7, 42.1, 42.0, 41.9, 25.4, 23.6, 23.5, 22.0, 21.9, 14.4.

**FT-IR** $\nu_{max}$ (cm$^{-1}$) 3299, 2932, 1730, 1664, 1610, 1584, 1511, 1477, 1450, 1370, 1320, 1302, 1247, 1175, 1108, 1078, 1033, 832, 759, 741

**HRMS (ESI)** m/z calculated for $[M+H]^+$ for $C_{30}H_{33}N_2O_6S$ 549.205935, found 549.20538.

**MP:** 94-96 °C

### 5. NMR

#### Ethyl 2-((S)-2-((((9H-fluoren-9-yl)methoxy)carbonyl)amino)propanamido)-2 hydroxyacetate (2a)

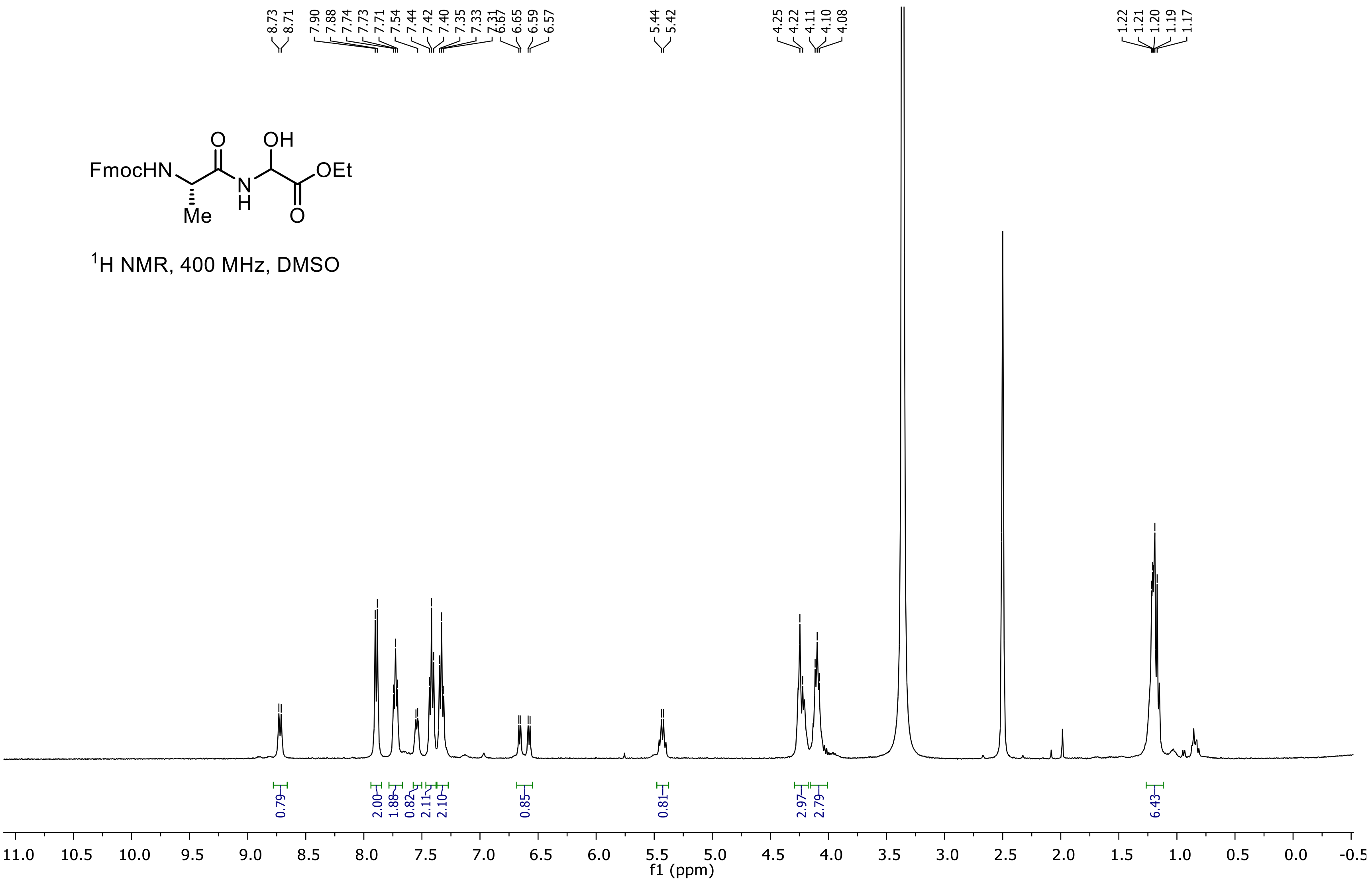

**Ethyl 2-((S)-2-((((9H-fluoren-9-yl)methoxy)carbonyl)amino)propanamido)-2 hydroxyacetate (2a)**

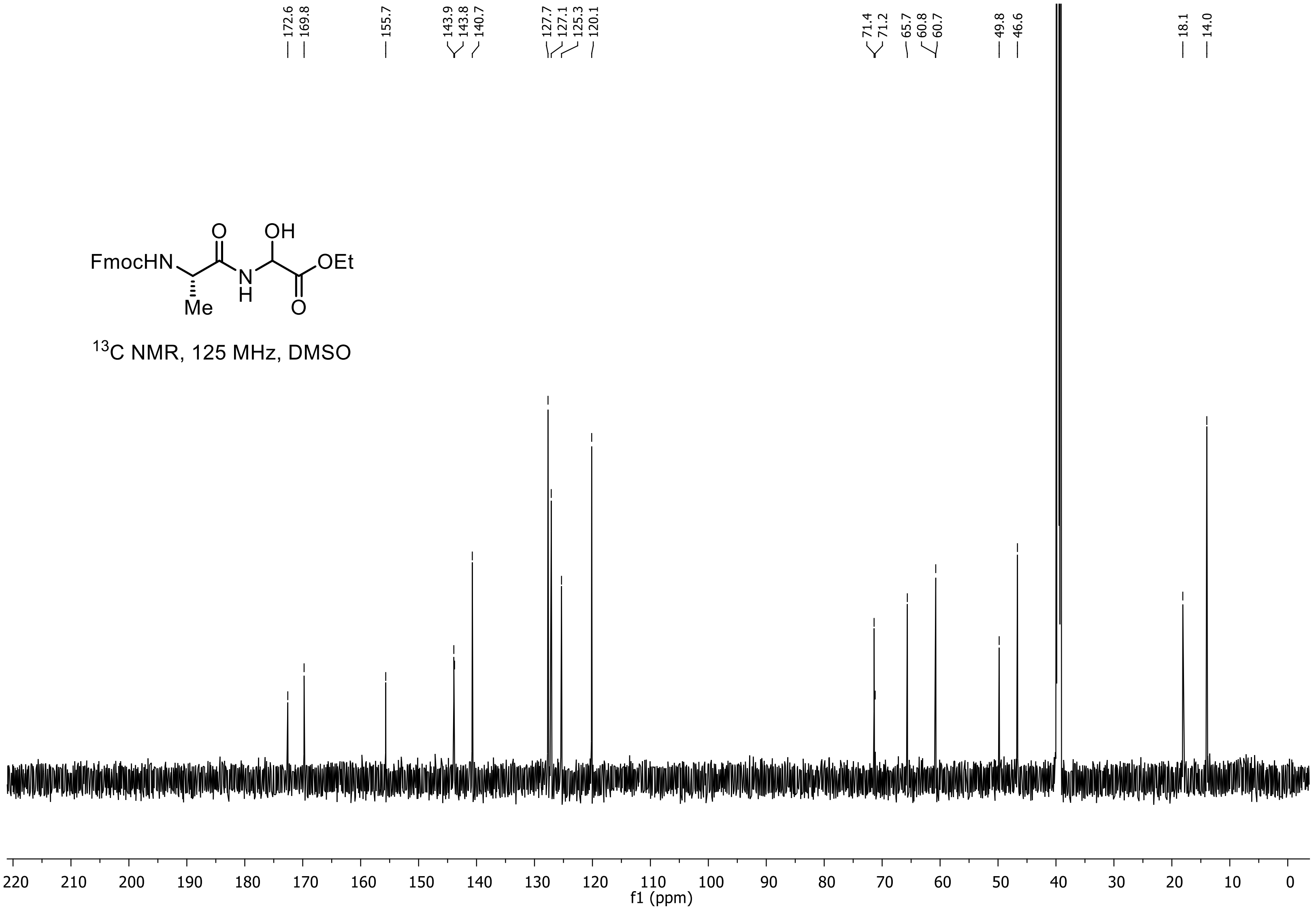

**Ethyl 2-((S)-2-((((9H-fluoren-9-yl)methoxy)carbonyl)amino)-3-methylbutanamido)-2-hydroxyacetate (2b)**

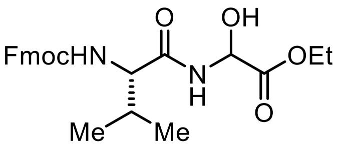


**2b**

$^{1}$H NMR, 500 MHz, CDCl3

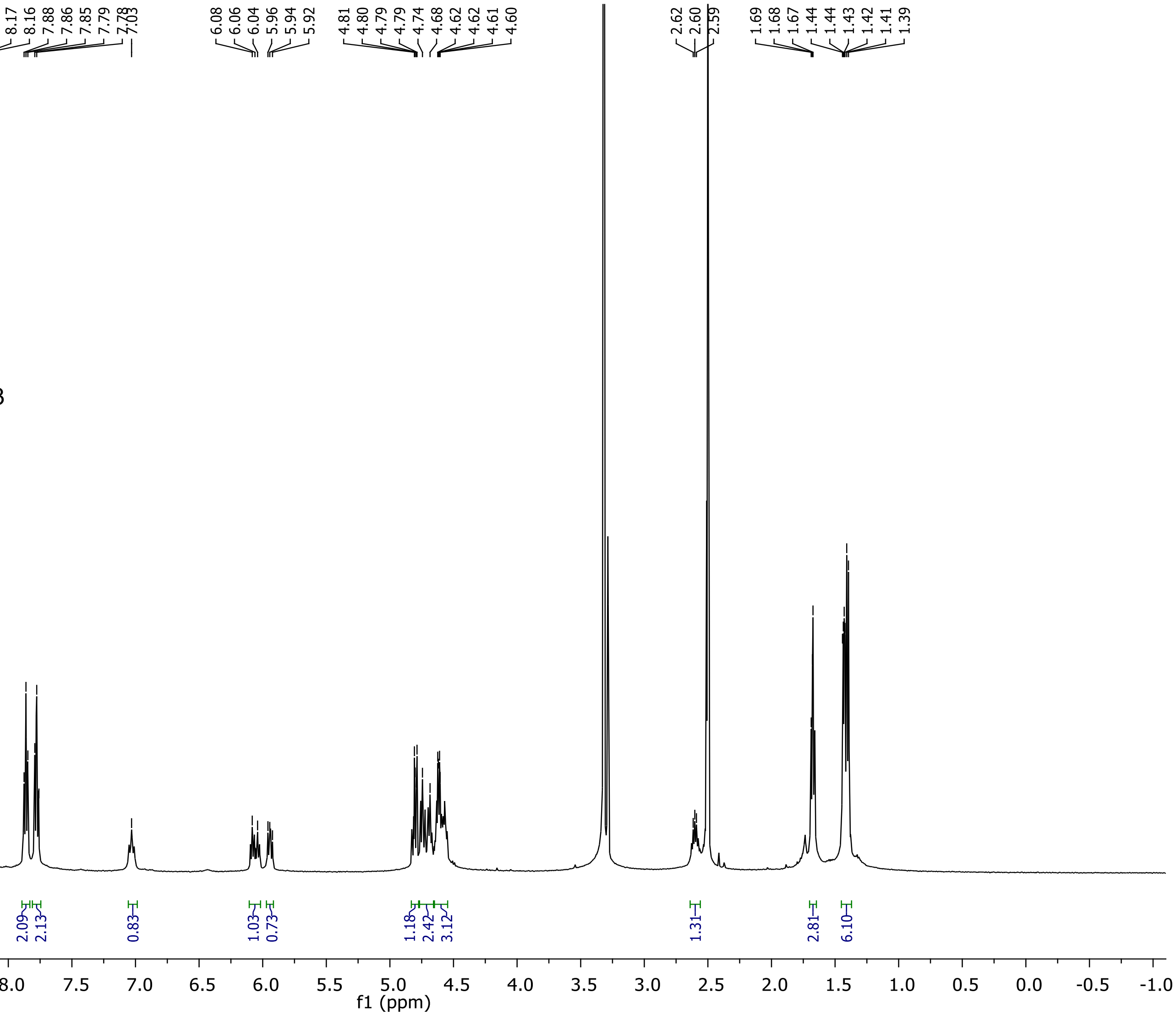

**Ethyl 2-((S)-2-((((9H-fluoren-9-yl)methoxy)carbonyl)amino)-3-methylbutanamido)-2-hydroxyacetate (2b)**

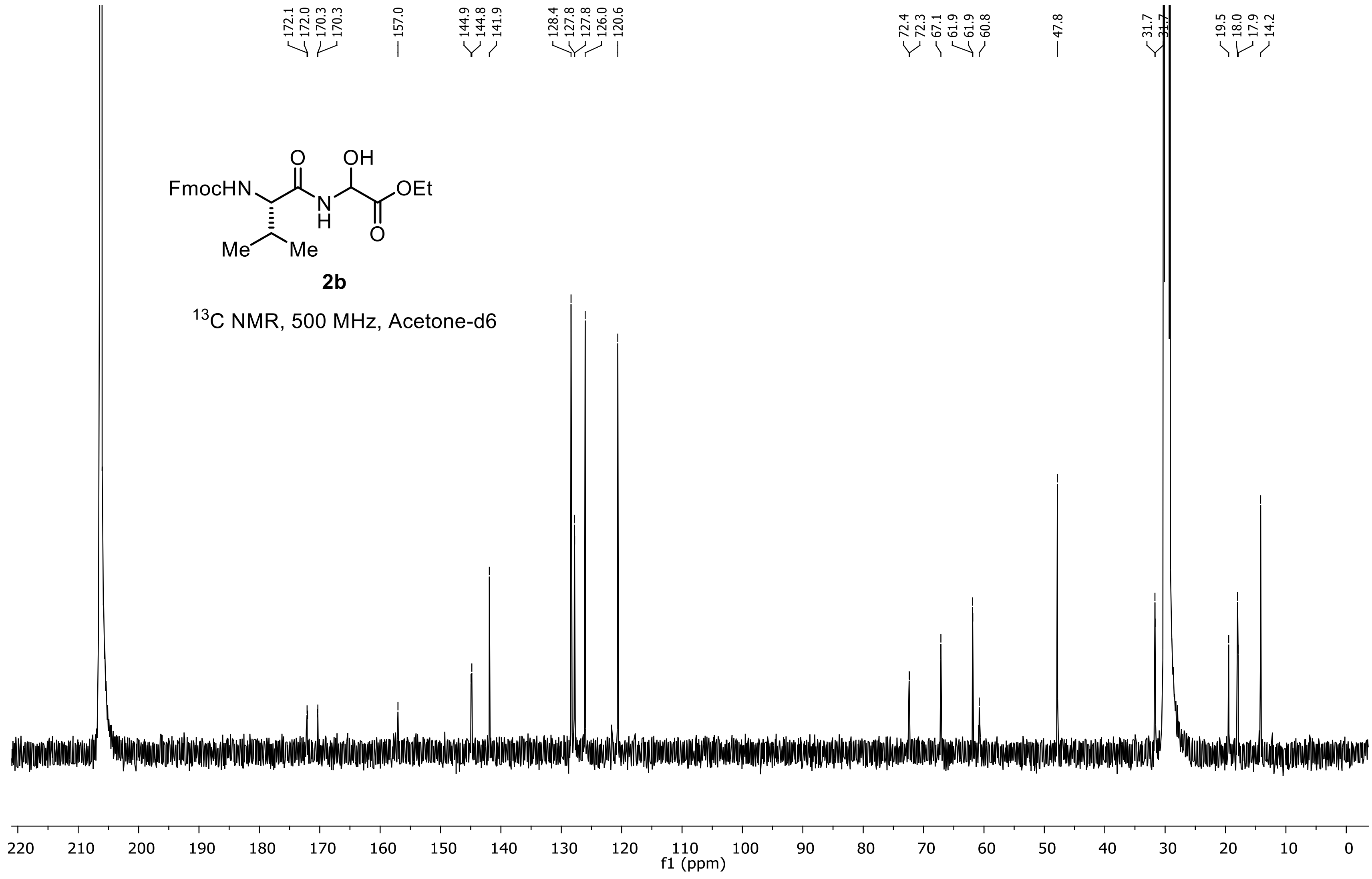

**Ethyl 2-((S)-2-((((9H-fluoren-9-yl)methoxy)carbonyl)amino)-4-methylpentanamido)-2-hydroxyacetate (2c)**

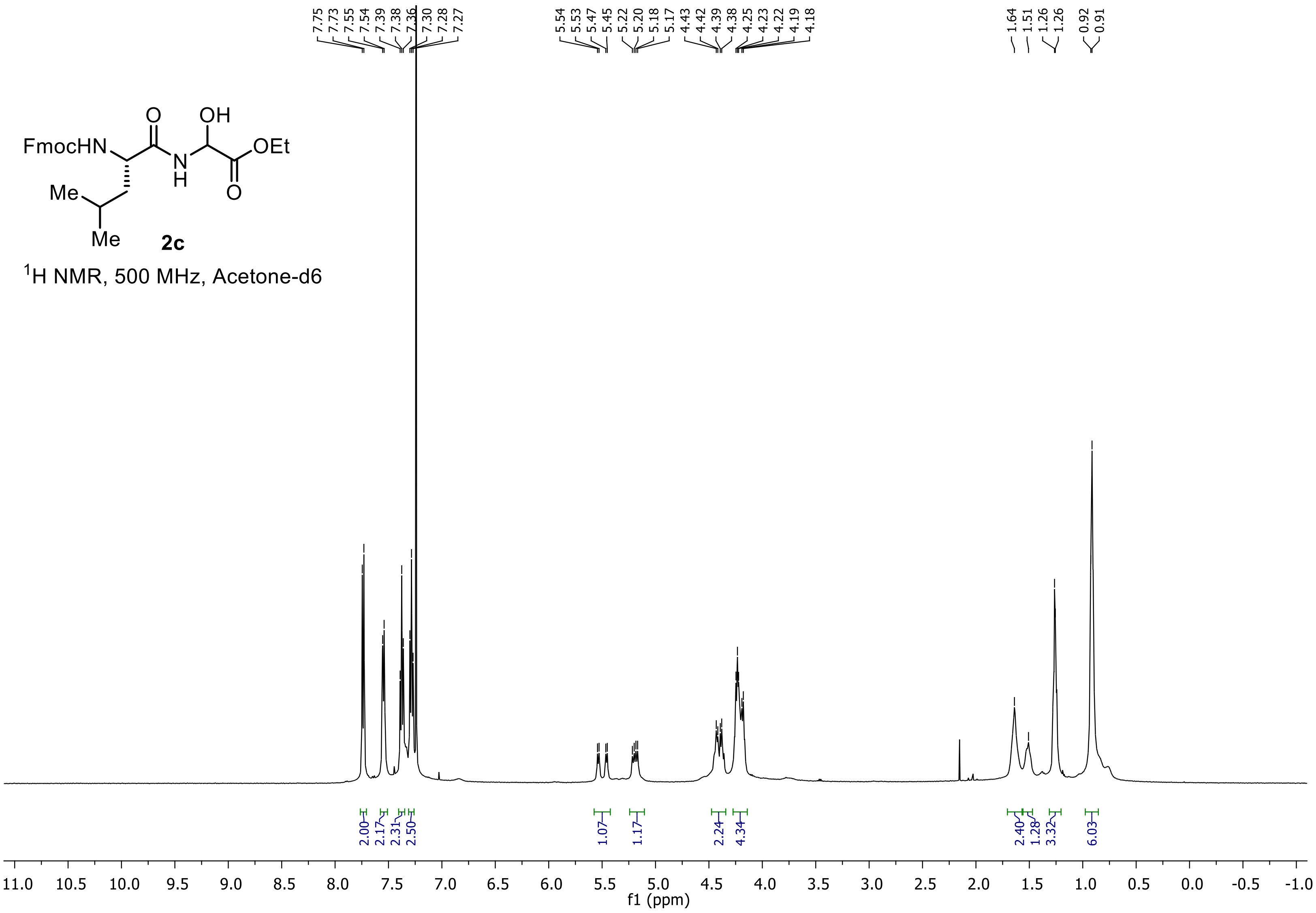

**Ethyl 2-((S)-2-((((9H-fluoren-9-yl)methoxy)carbonyl)amino)-4-methylpentanamido)-2-hydroxyacetate (2c)**

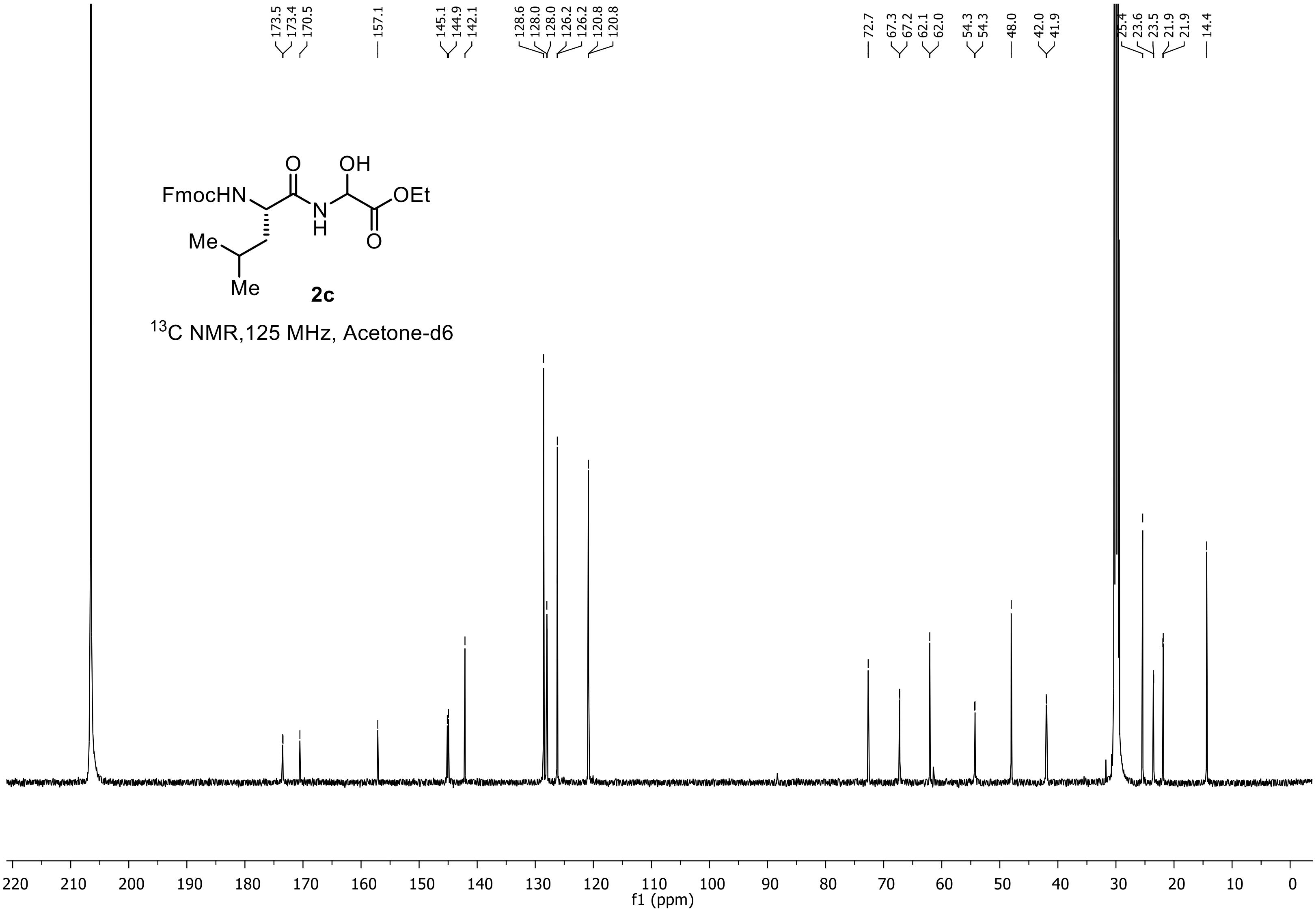

**Ethyl 2-((2S,3S)-2-((((9H-fluoren-9-yl)methoxy)carbonyl)amino)-3-methylpentanamido)-2-hydroxyacetate (2d)**

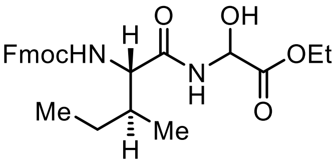


**2d**

$^1$H NMR, 500 MHz, DMSO-d6

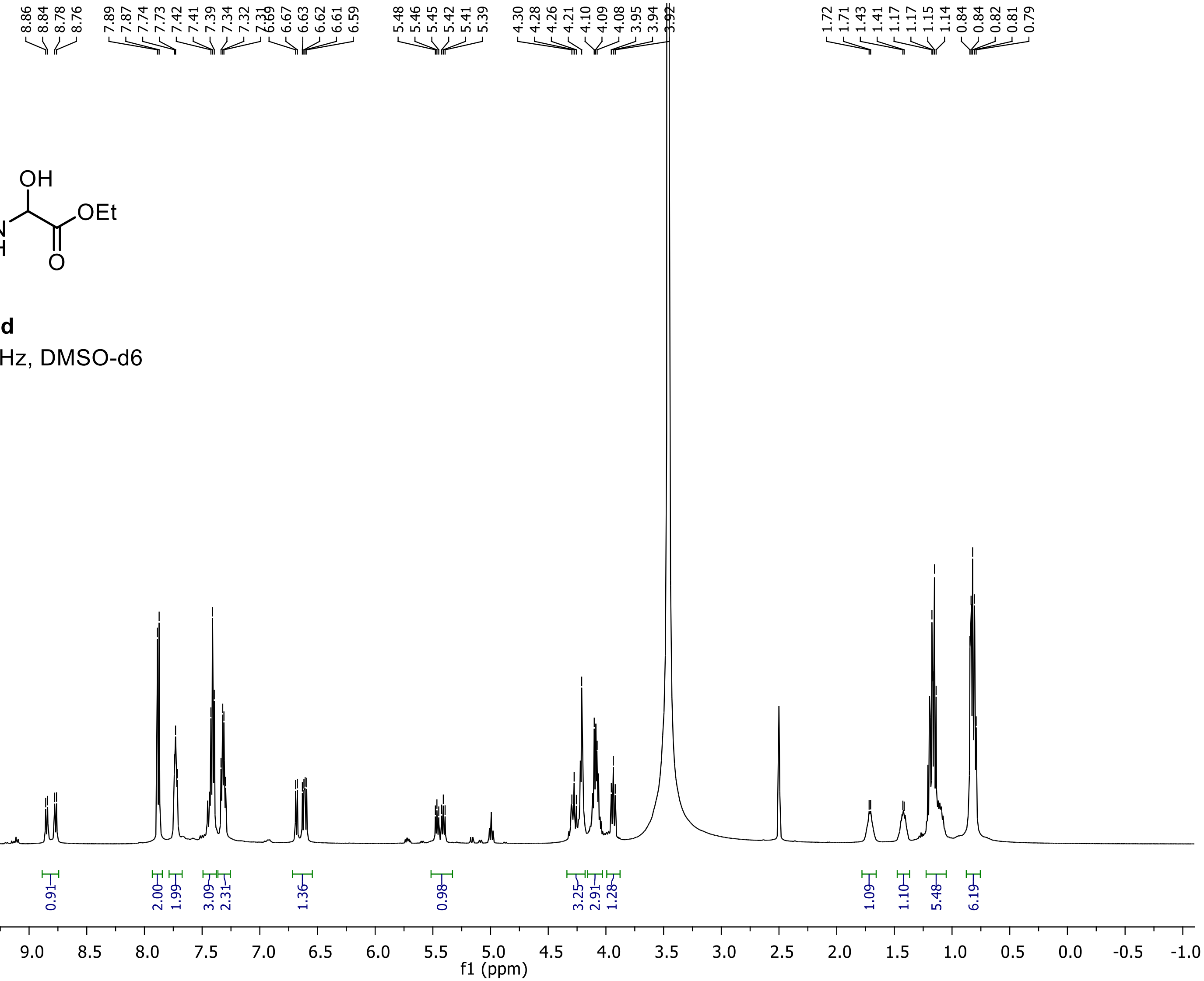

**Ethyl 2-((2S,3S)-2-((((9H-fluoren-9-yl)methoxy)carbonyl)amino)-3-methylpentanamido)-2-hydroxyacetate (2d)**

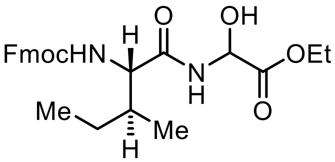

**2d**

$^{13}$C NMR,125 MHz, DMSO-d6

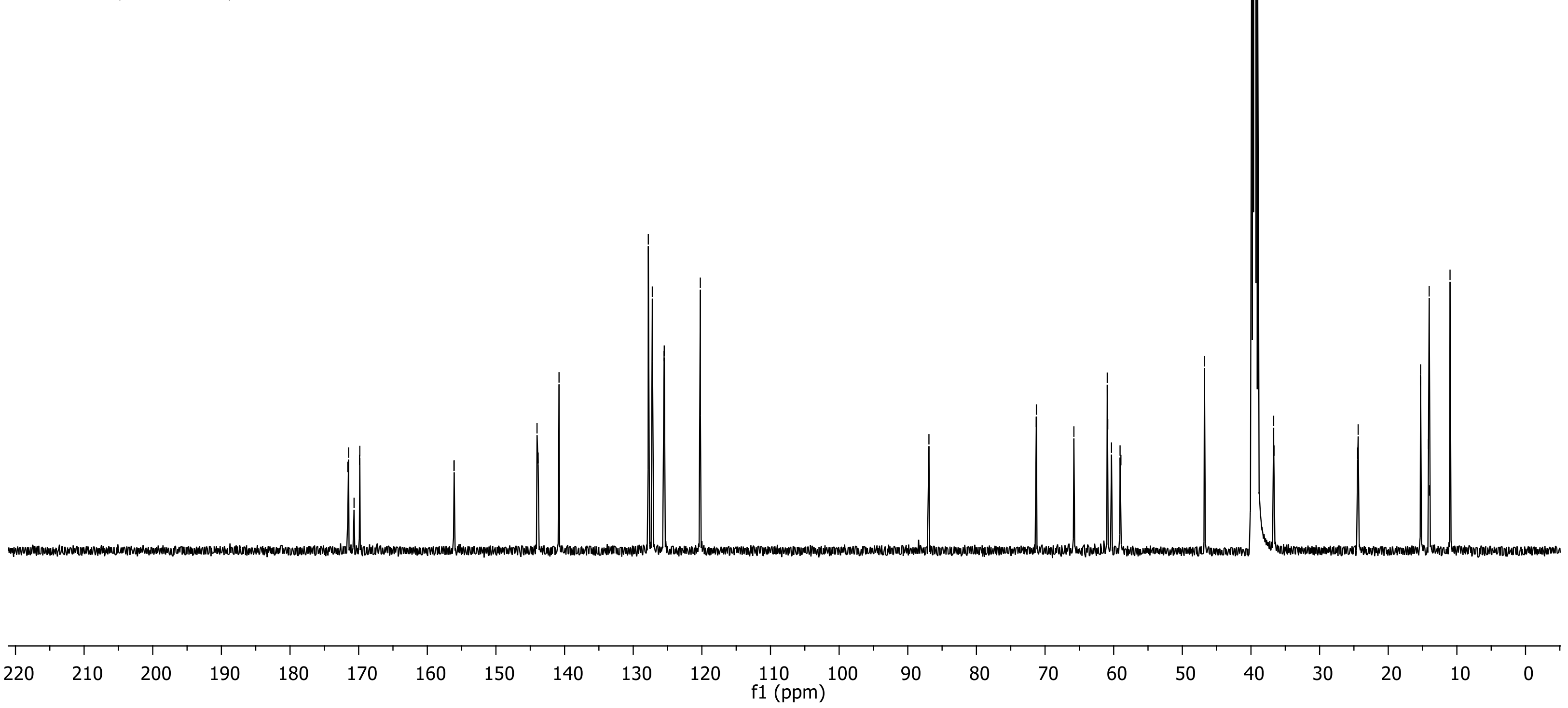

### Ethyl 2-((S)-2-((((9H-fluoren-9-yl)methoxy)carbonyl)amino)pentanamido)-2-hydroxyacetate (2e)

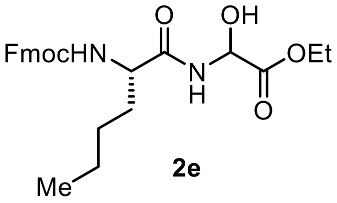


**2e**

$^1$H NMR, 500 MHz, CDCl3

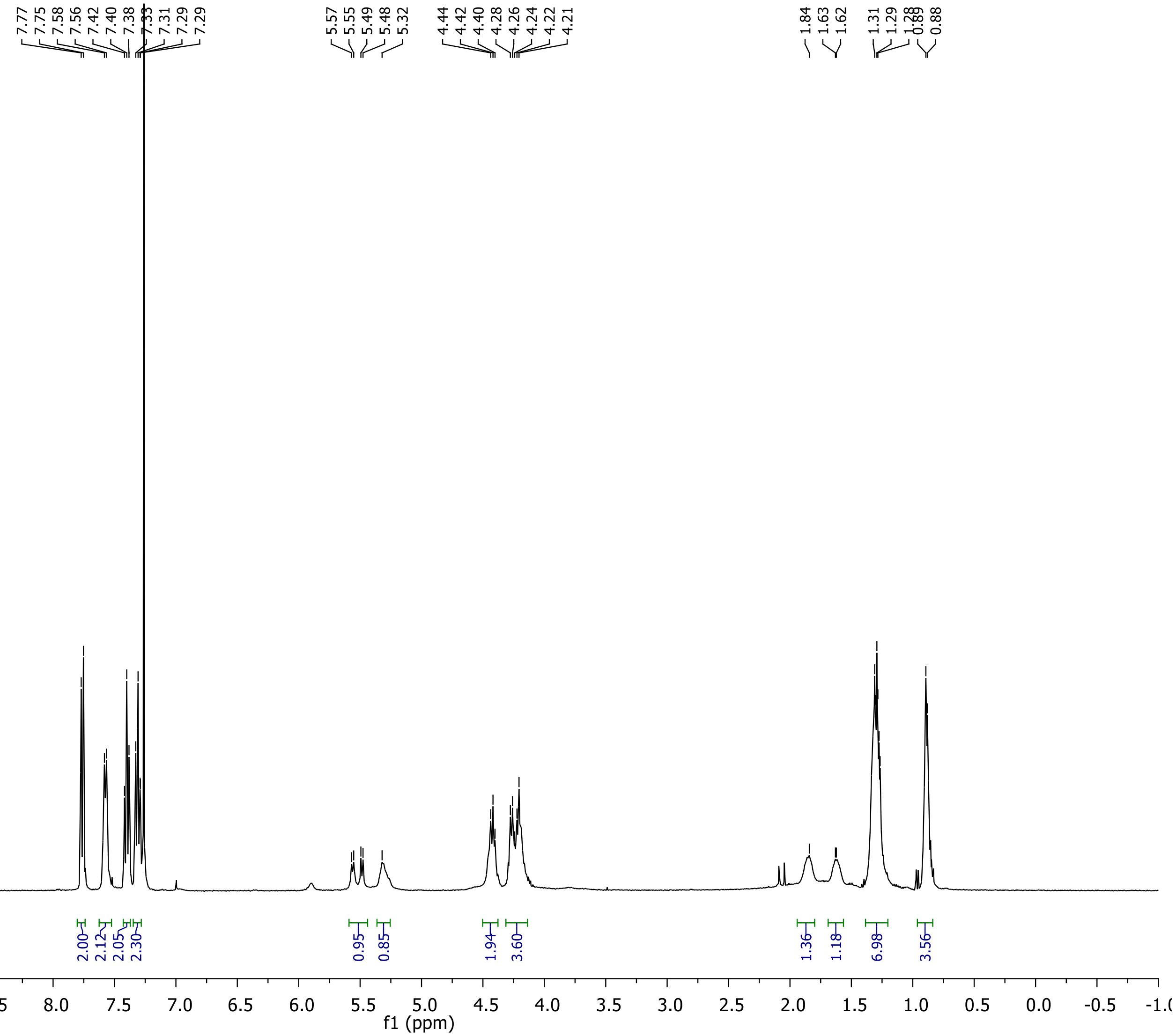

**Ethyl 2-((S)-2-((((9H-fluoren-9-yl)methoxy)carbonyl)amino)pentanamido)-2-hydroxyacetate (2e)**

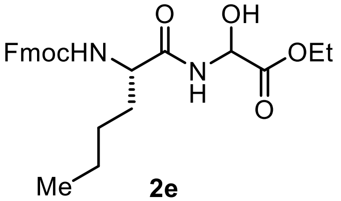


**2e**

$^{13}C$ NMR,125 MHz, DMSO-d6

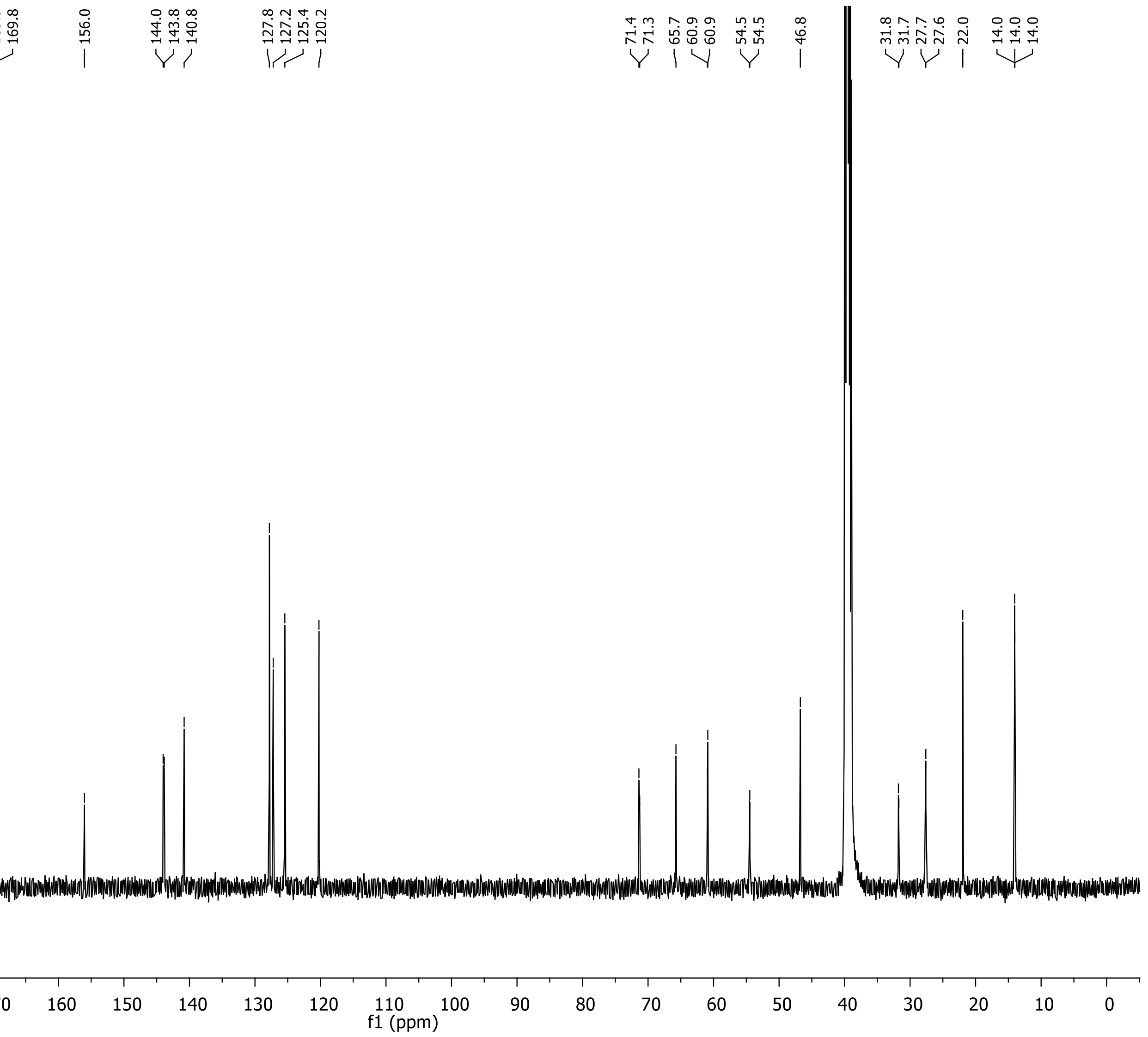

**Ethyl 2-(2-((((9H-fluoren-9-yl)methoxy)carbonyl)amino)acetamido)-2-hydroxyacetate (2f)**

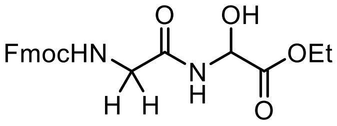


**2f**

$^{1}$H NMR, 500 MHz, CDCl3

O = Unknown impurities

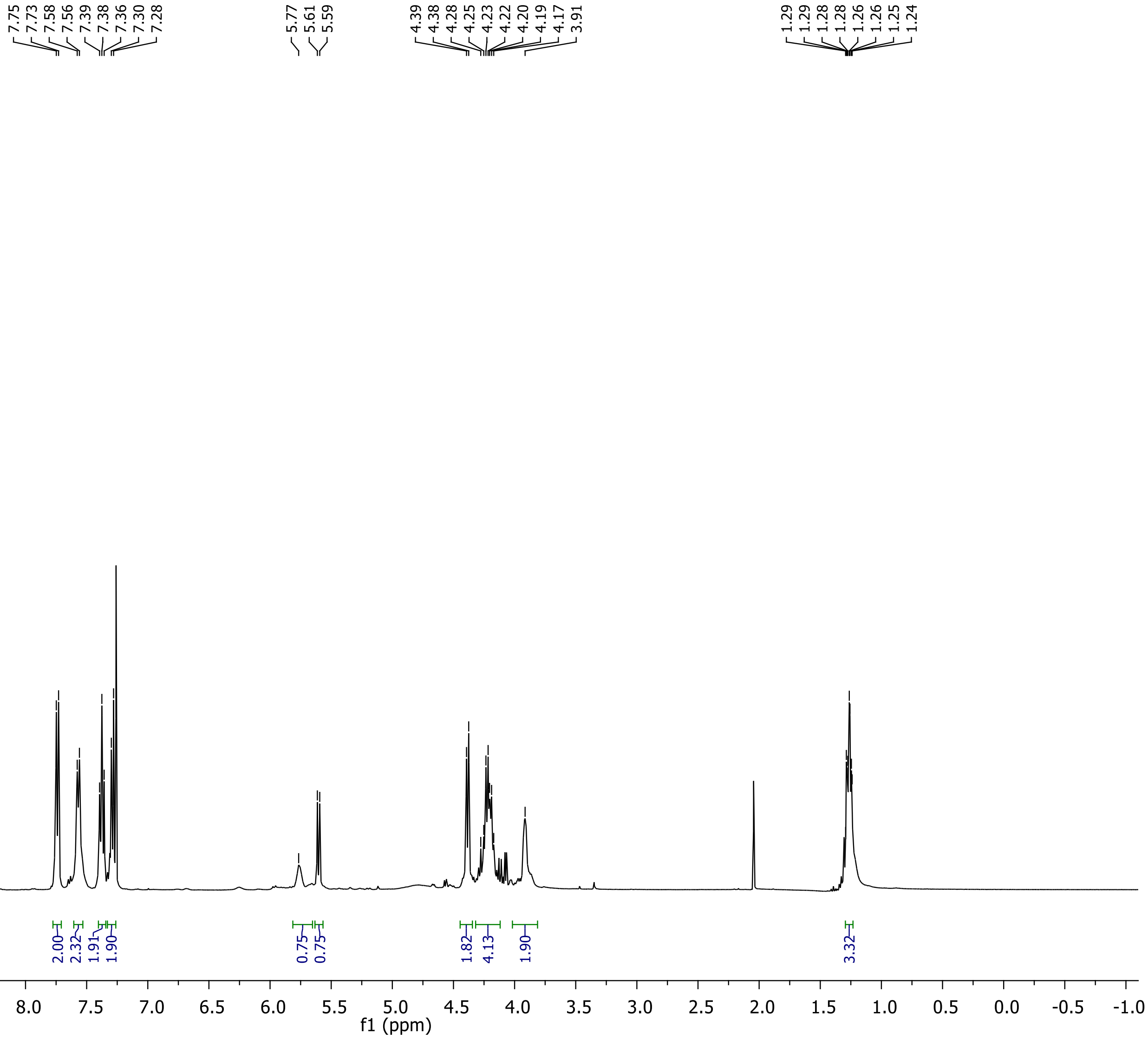

**Ethyl 2-(2-((((9H-fluoren-9-yl)methoxy)carbonyl)amino)acetamido)-2-hydroxyacetate (2f)**

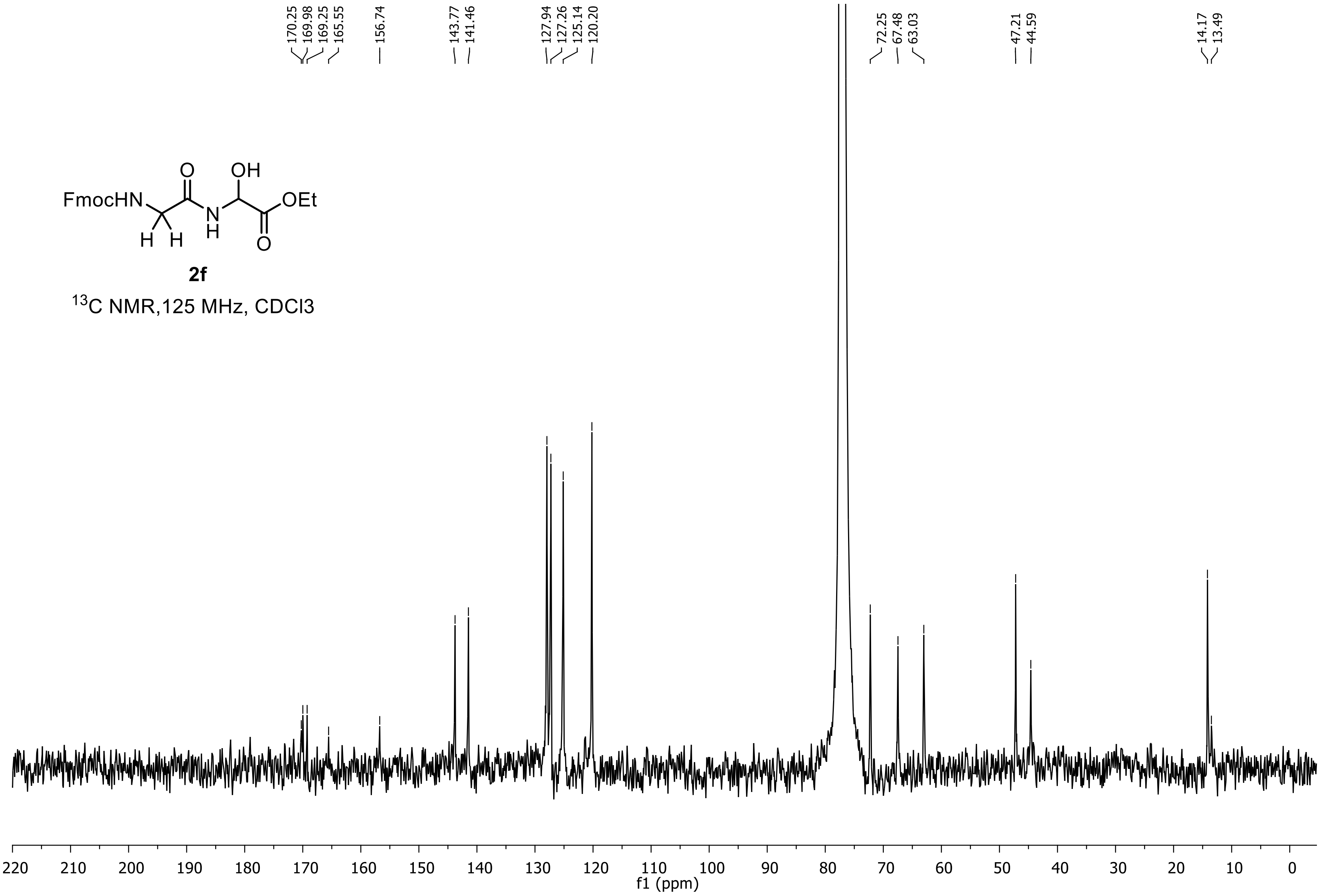


**2f**

$^{13}$C NMR,125 MHz, CDCl3

**Ethyl 2-((S)-2-((((9H-fluoren-9-yl)methoxy)carbonyl)amino)-4-(methylthio)butanamido)-2-hydroxyacetate (2g)**

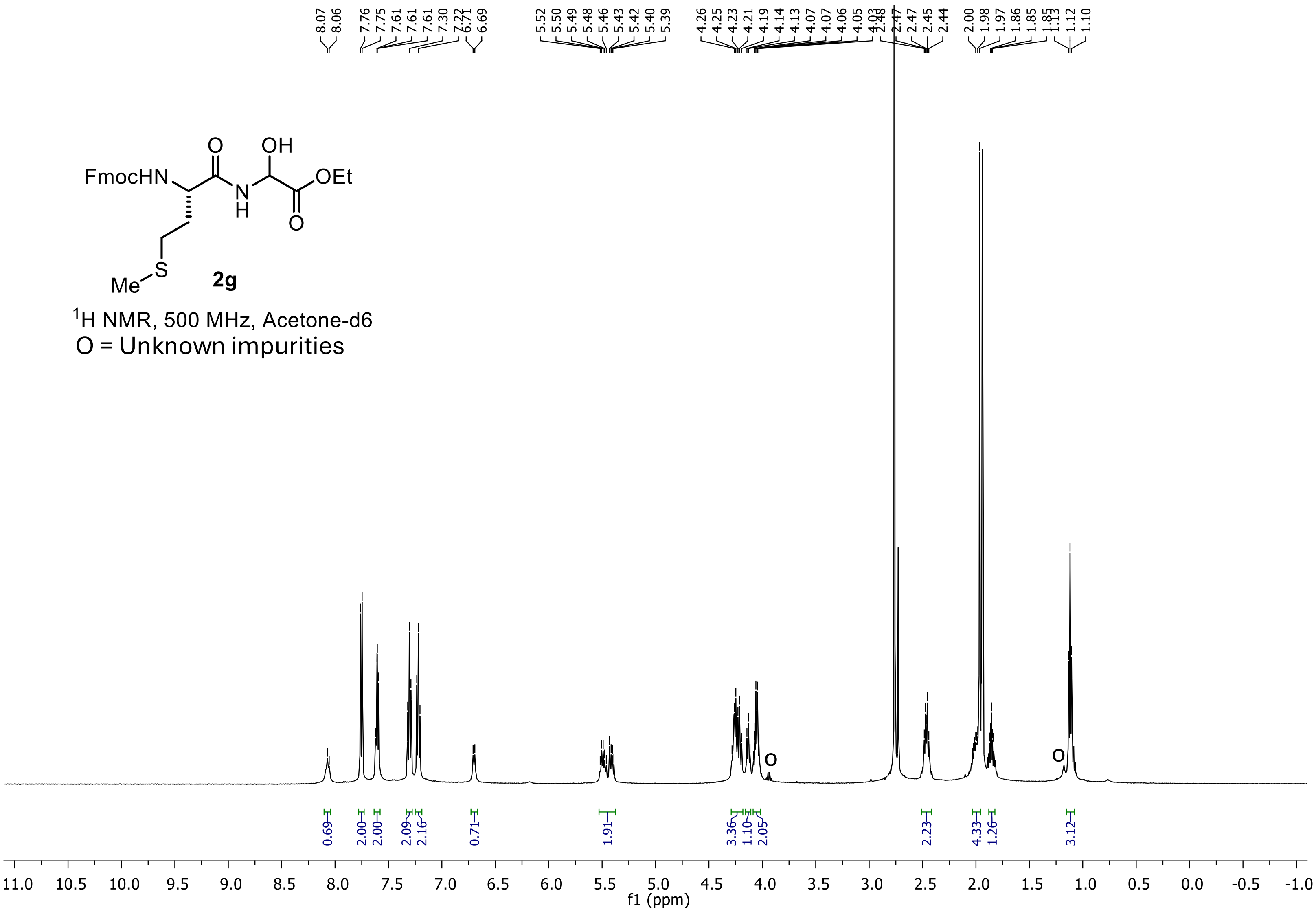

**Ethyl 2-((S)-2-((((9H-fluoren-9-yl)methoxy)carbonyl)amino)-4-(methylthio)butanamido)-2-hydroxyacetate (2g)**

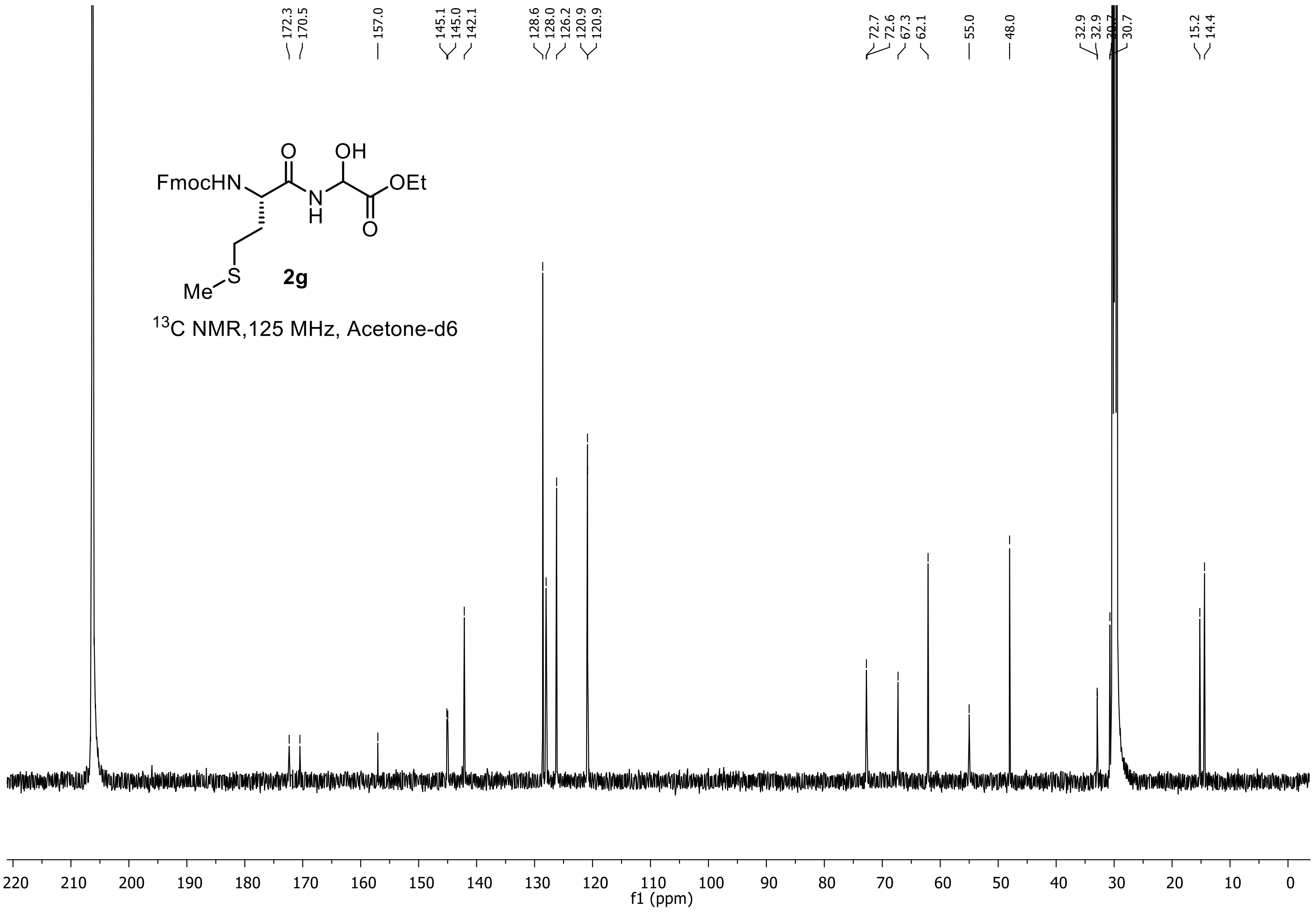

**Ethyl 2-((S)-2-((((9H-fluoren-9-yl)methoxy)carbonyl)amino)-3-phenylpropanamido)-2-hydr oxyacetate (2h)**

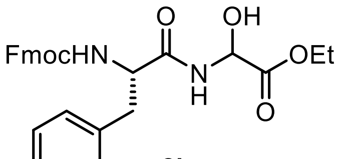


**2h**

$^{1}$H NMR, 500 MHz, Acetone-d6

O = Unknown impurities

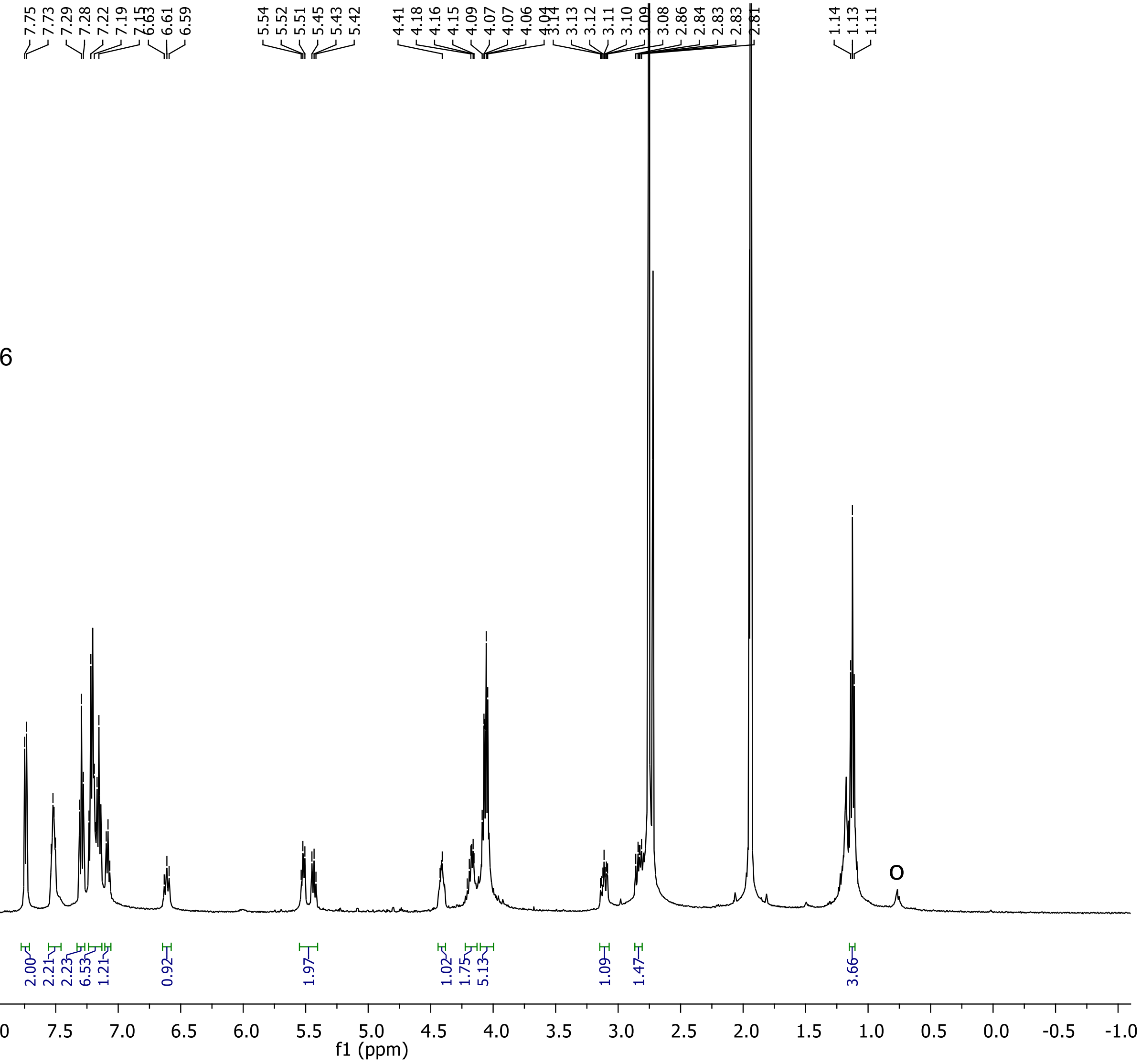

**Ethyl 2-((S)-2-((((9H-fluoren-9-yl)methoxy)carbonyl)amino)-3-phenylpropanamido)-2-hydr oxyacetate (2h)**

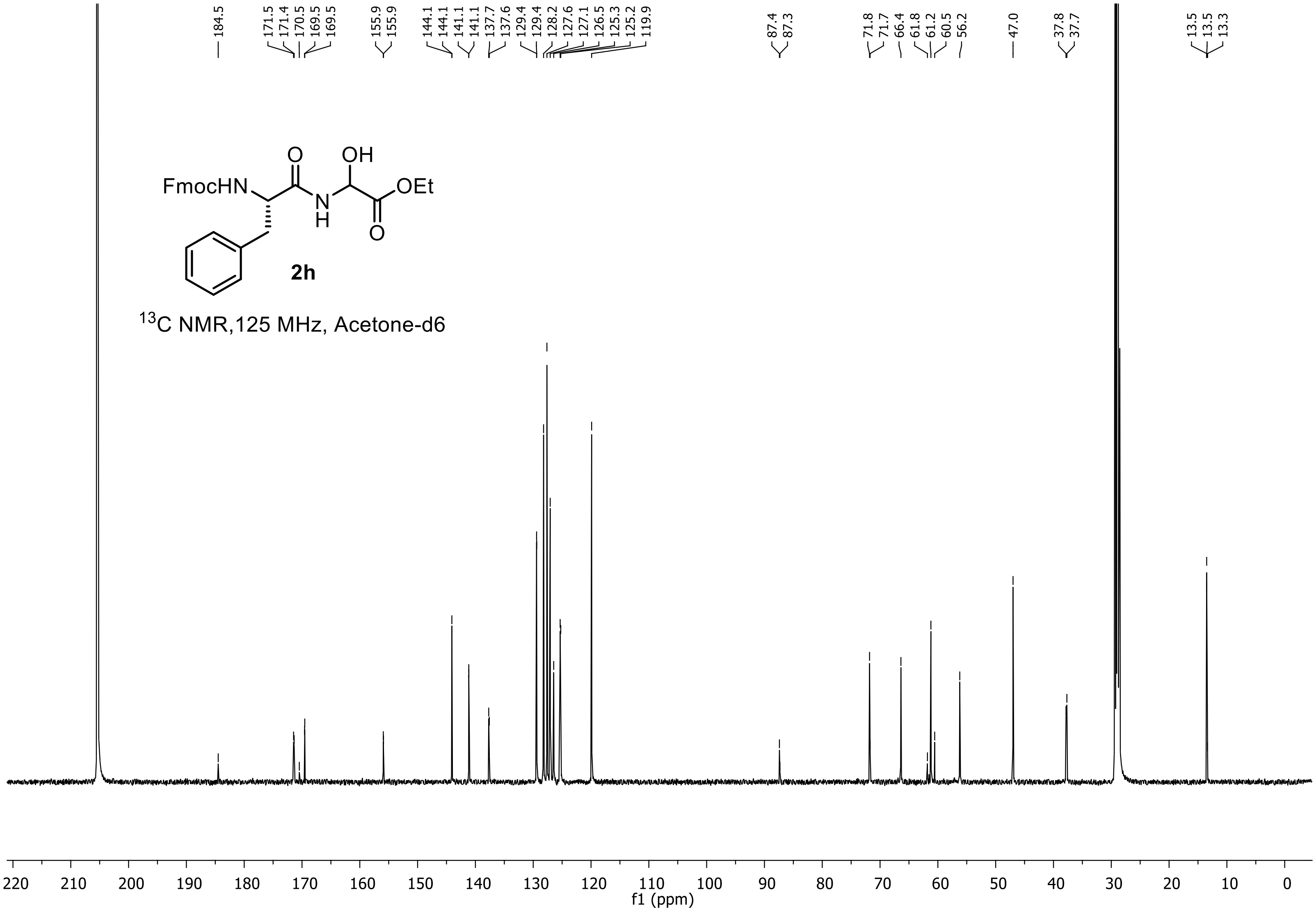

**(9H-fluoren-9-yl)methyl (2S)-2-((2-ethoxy-1-hydroxy-2-oxoethyl)carbamoyl)pyrrolidine-1-carboxylate (2i)**

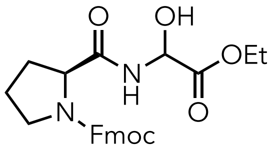


**2i**,

$^{1}$H-NMR, 500 MHz, $CDCl_3$

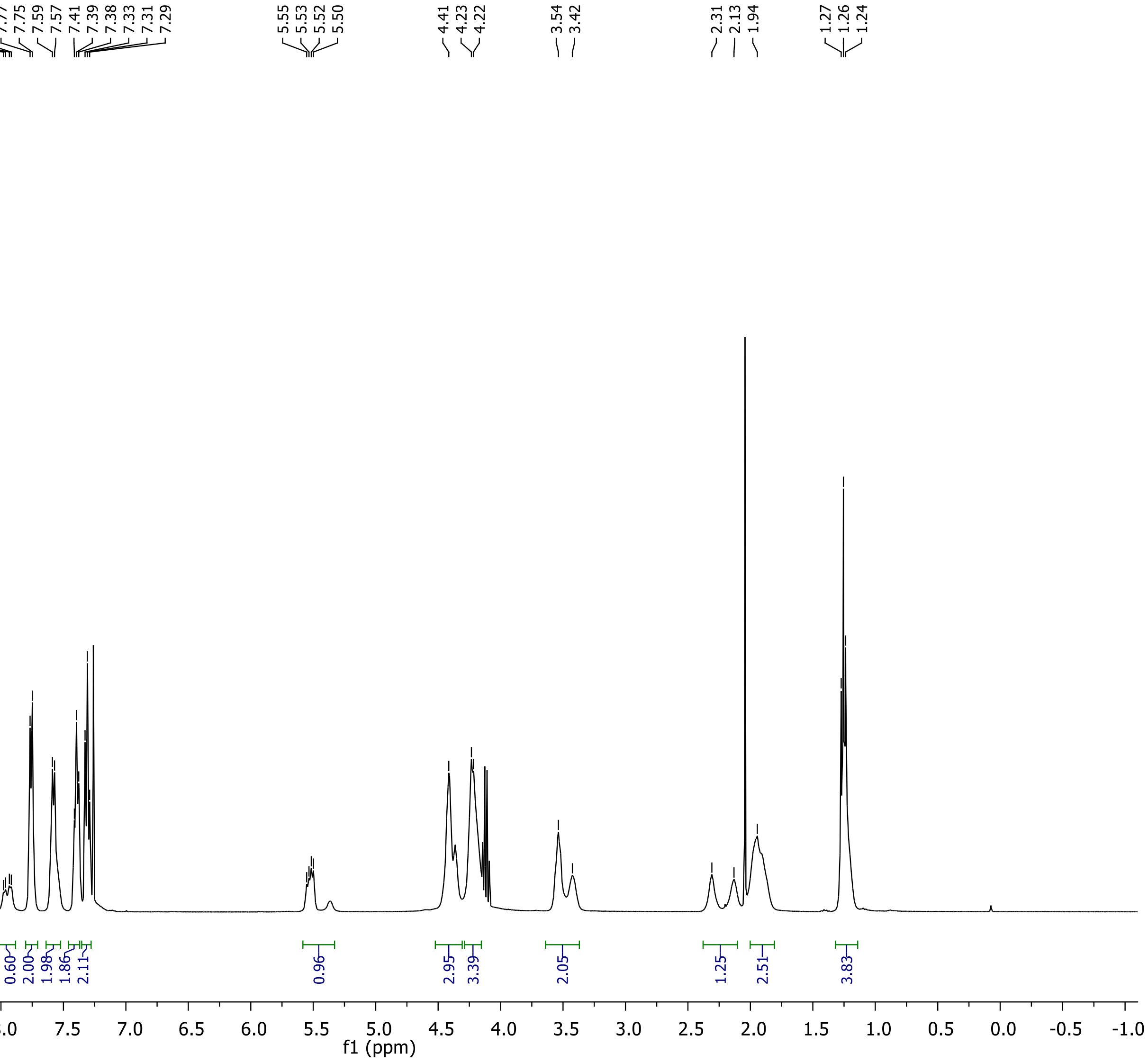

**(9H-fluoren-9-yl)methyl (2S)-2-((2-ethoxy-1-hydroxy-2-oxoethyl)carbamoyl)pyrrolidine-1-carboxylate (2i)**

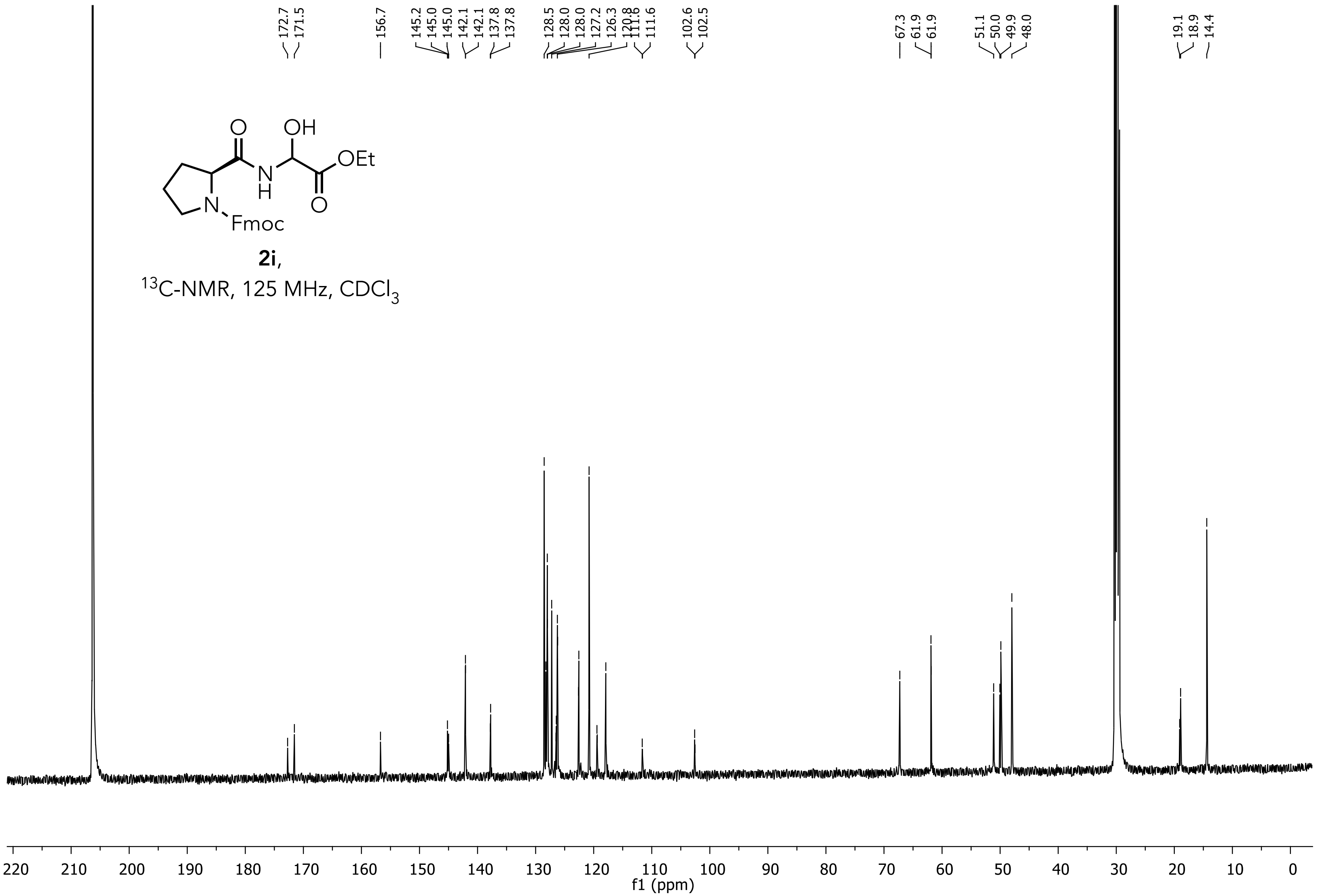

Ethyl2-((S)-2-((((9H-fluoren-9-yl)methoxy)carbonyl)amino)-6-((tertbutoxycarbonyl)amino) hexanamido)- 2-hydroxyacetate (2j)

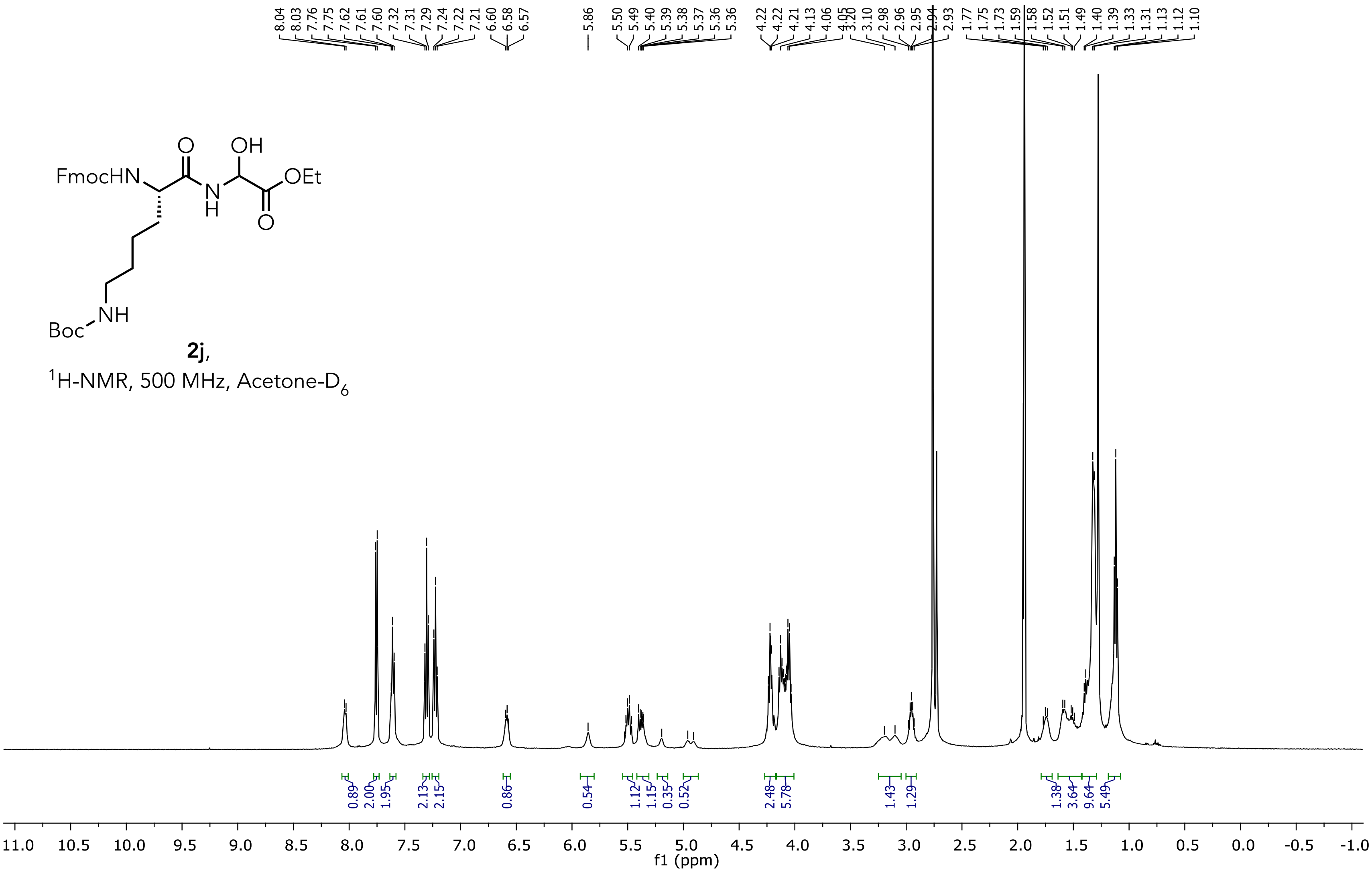

**Ethyl2-((S)-2-((((9H-fluoren-9-yl)methoxy)carbonyl)amino)-6-((tertbutoxycarbonyl)amino) hexanamido)- 2-hydroxyacetate (2j)**

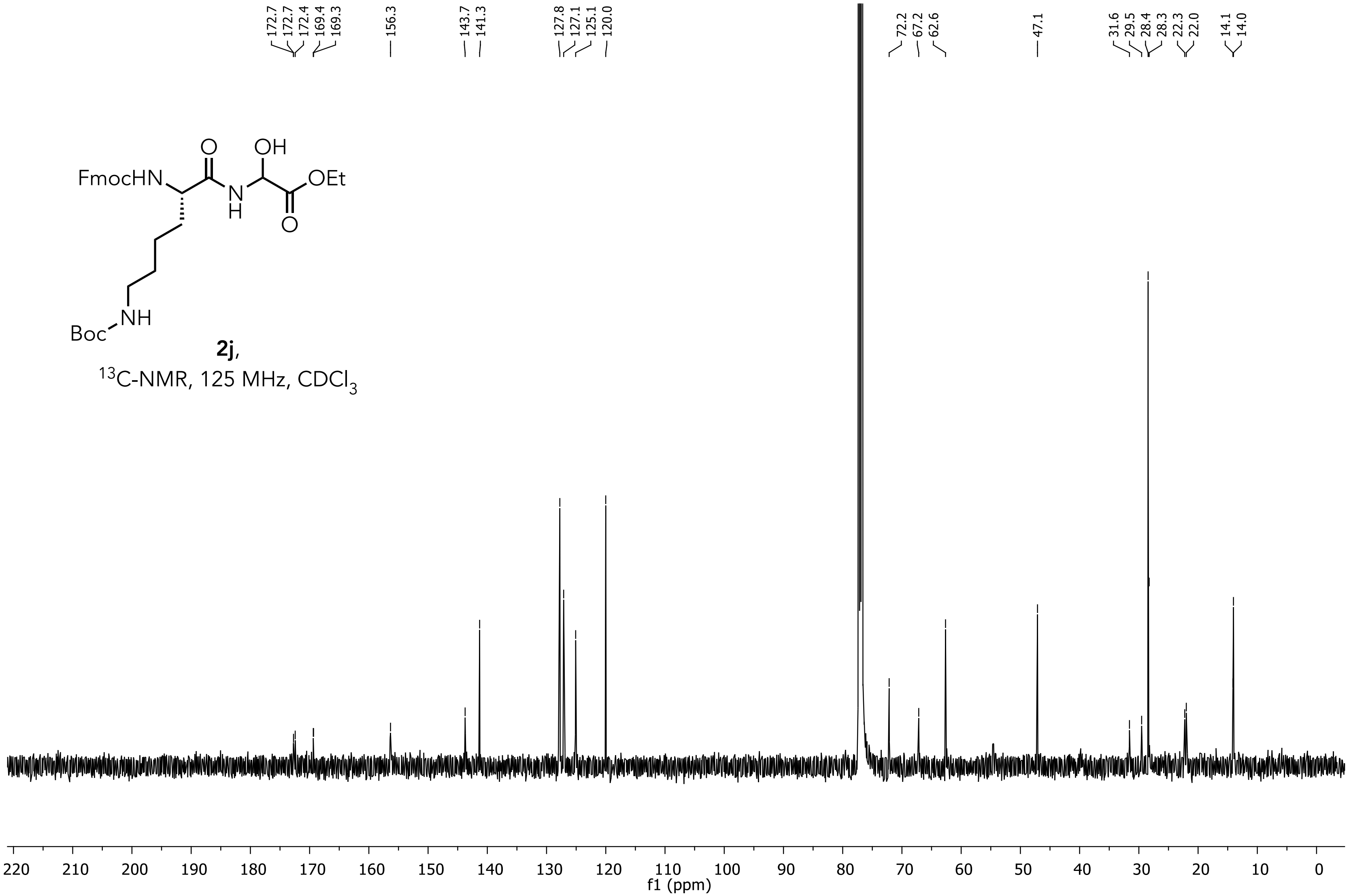

**Ethyl 2-((S)-2-((((9H-fluoren-9-yl)methoxy)carbonyl)amino)-3-(4-(tert-butoxy)phenyl)prop anamido)-2-hydroxyacetate (2k)**

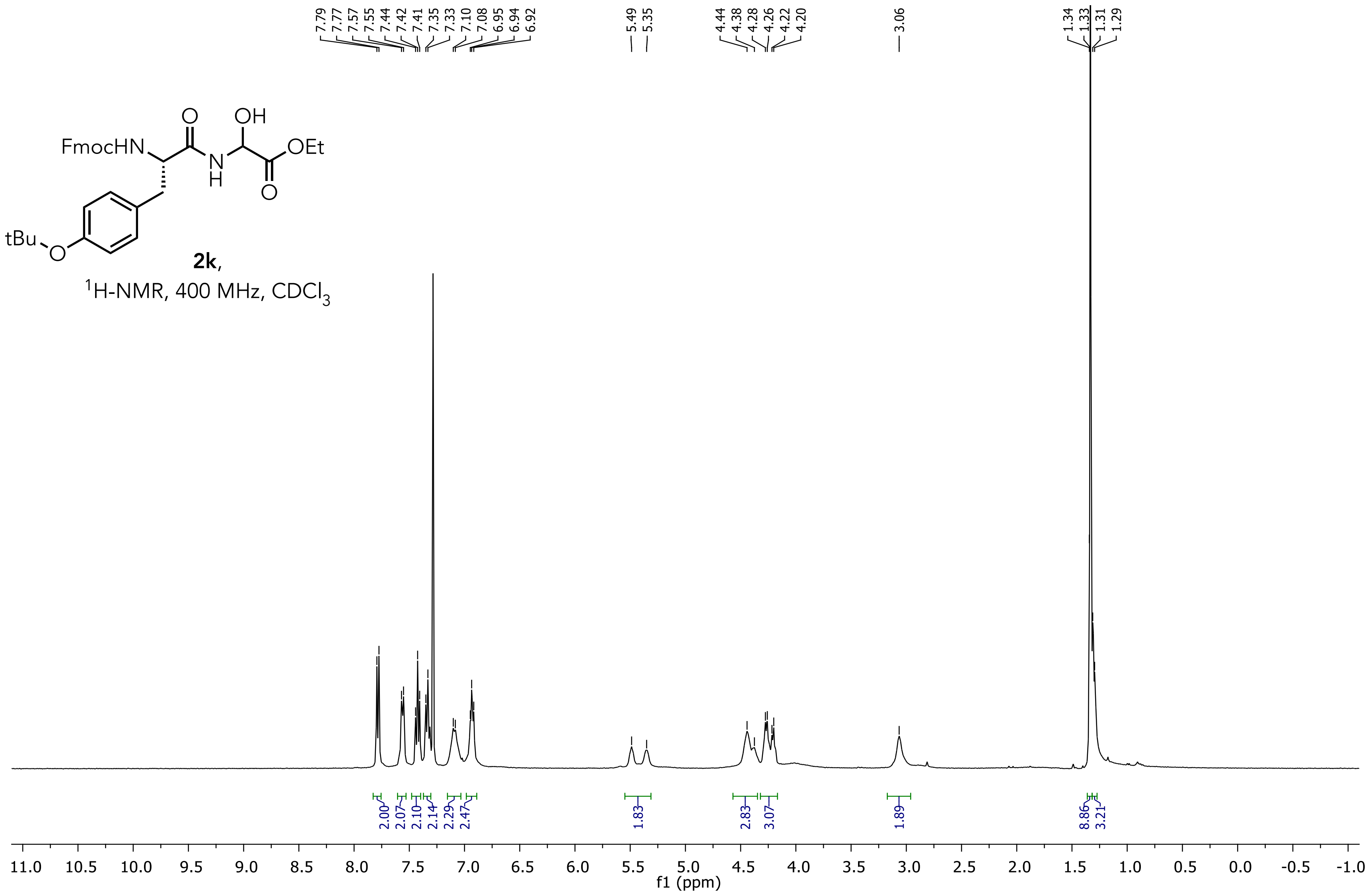

**Ethyl 2-((S)-2-((((9H-fluoren-9-yl)methoxy)carbonyl)amino)-3-(4-(tert-butoxy)phenyl)prop anamido)-2-hydroxyacetate (2k)**

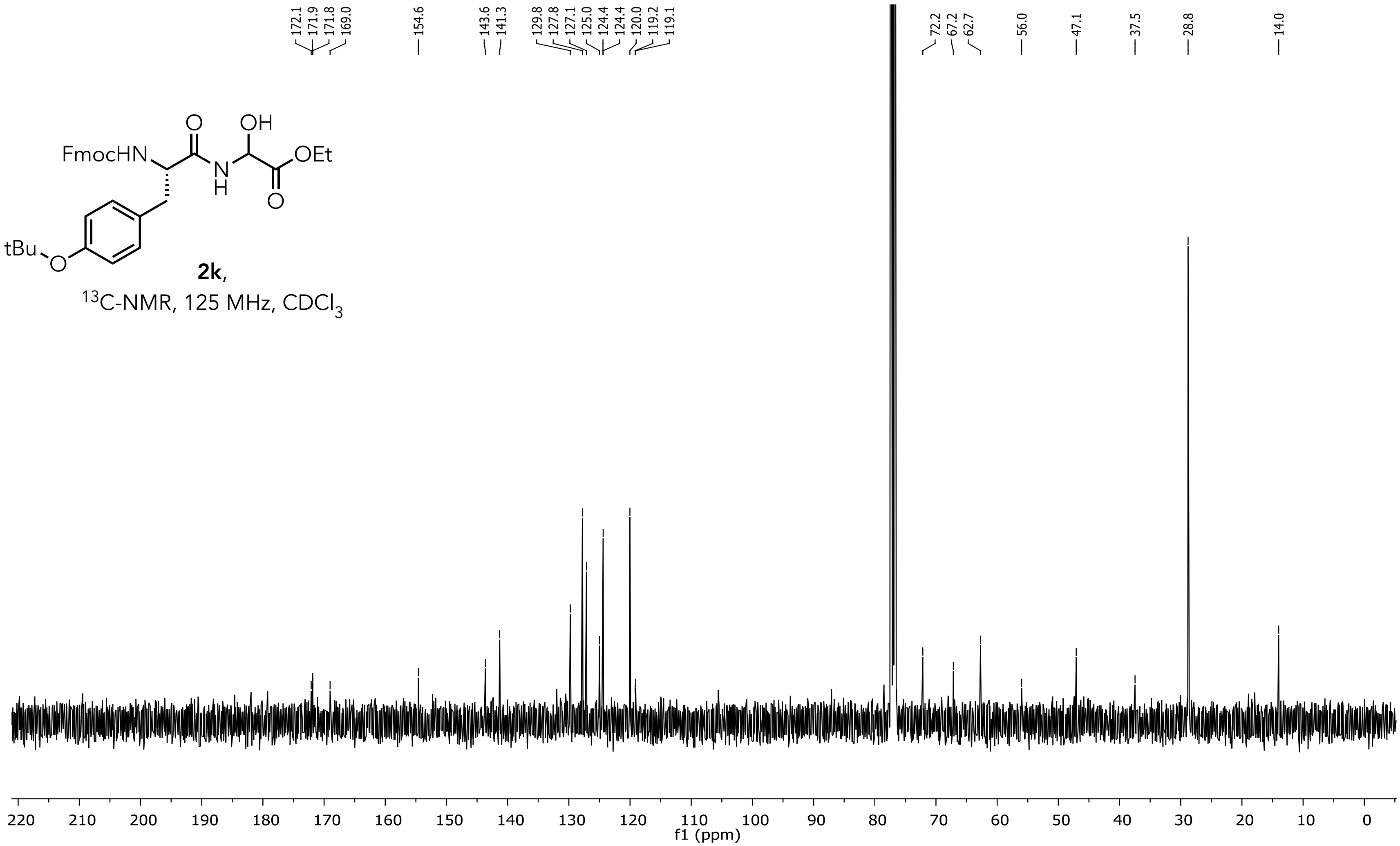

**Ethyl 2-((S)-2-((((9H-fluoren-9-yl)methoxy)carbonyl)amino)-3-(tert-butoxy)propanamido)-2-hydroxyacetate (2l)**

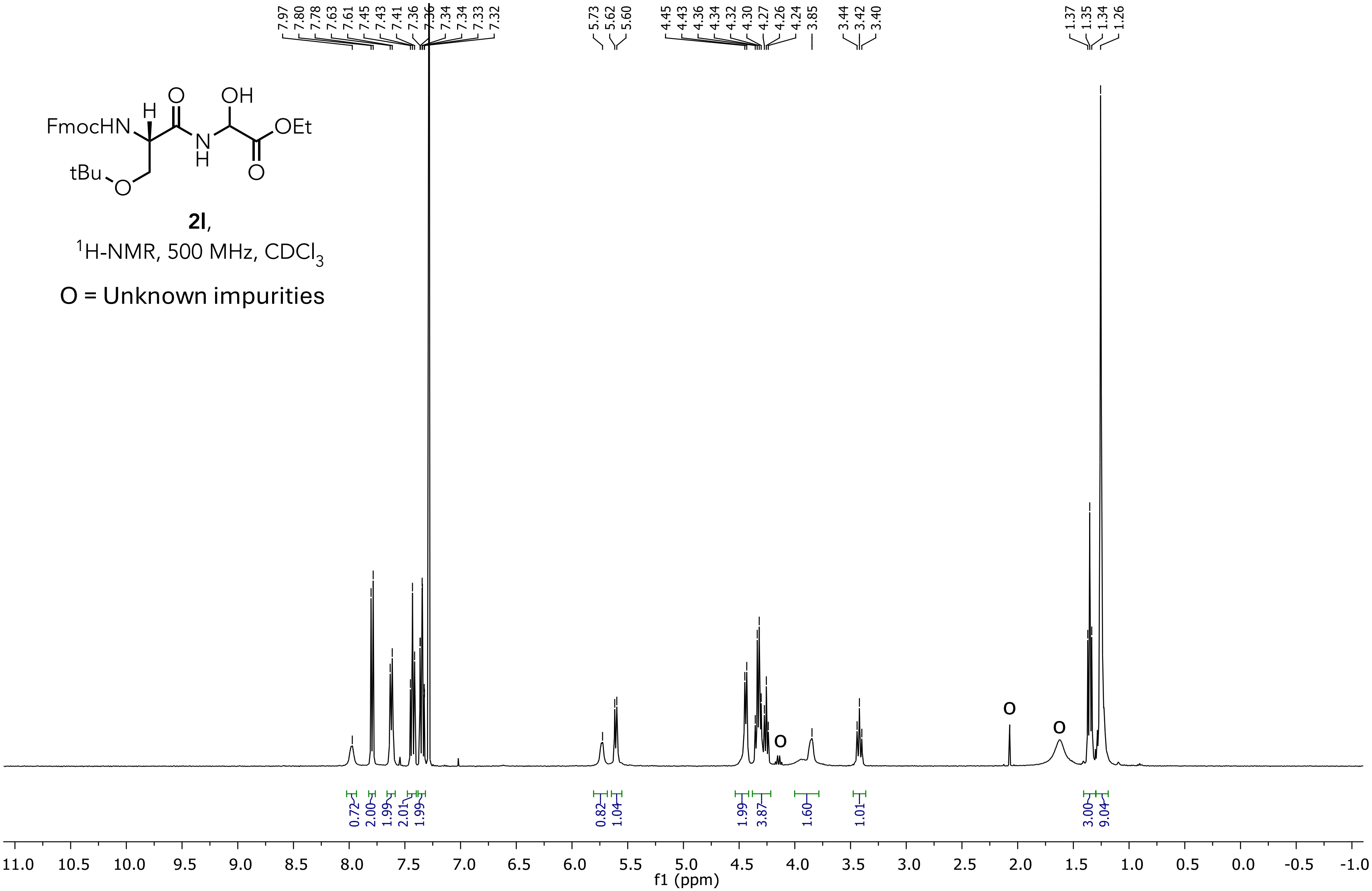

**Ethyl 2-((S)-2-((((9H-fluoren-9-yl)methoxy)carbonyl)amino)-3-(tert-butoxy)propanamido)-2-hydroxyacetate (2l)**

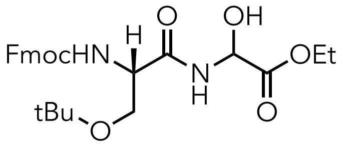


**2l**,
$^{13}$C-NMR, 125 MHz, $CDCl_3$

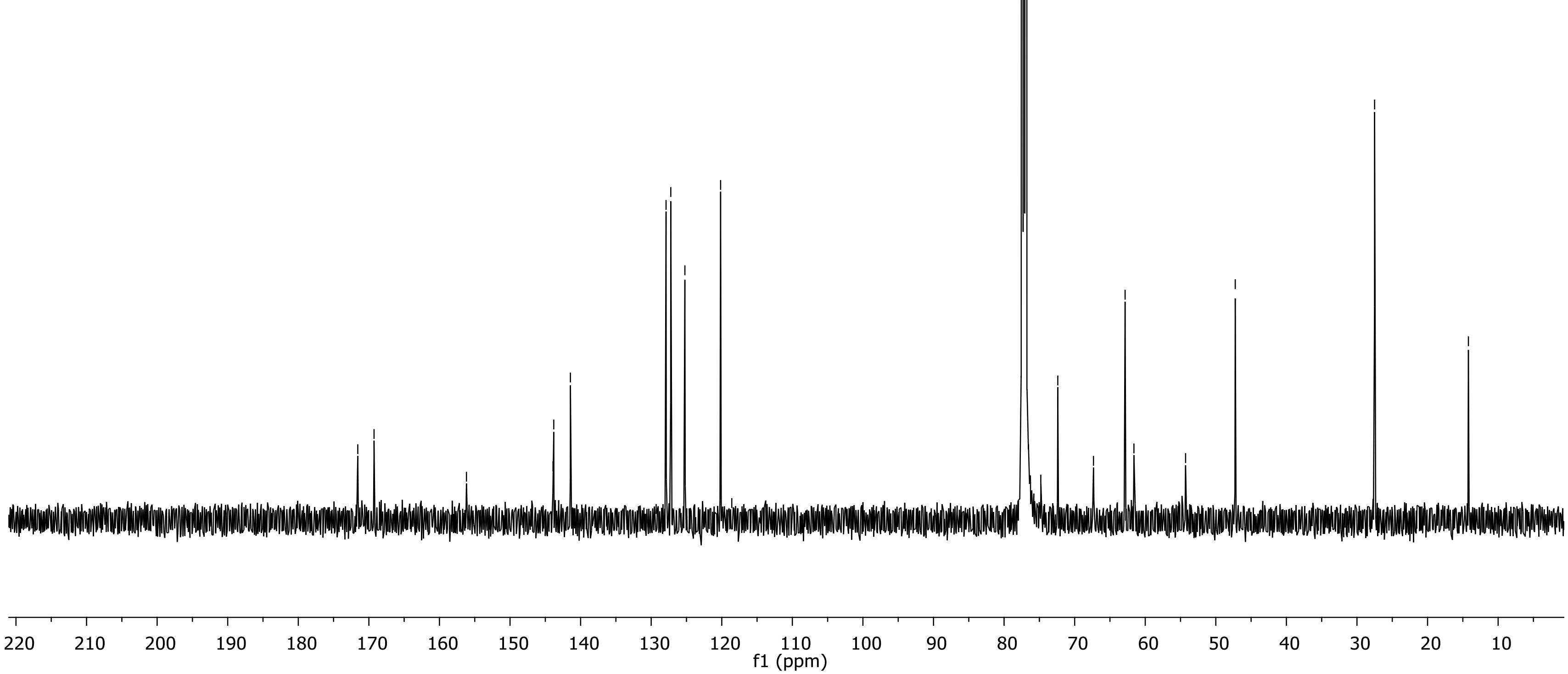

### Ethyl 2-((2S,3R)-2-((((9H-fluoren-9-yl)methoxy)carbonyl)amino)-3-(tert-butoxy)butanamido)-2-hydroxyacetate (2m)

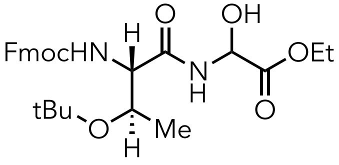


**2m**,
$^{1}$H-NMR, 500 MHz, $CDCl_3$
O = Unknown impurities

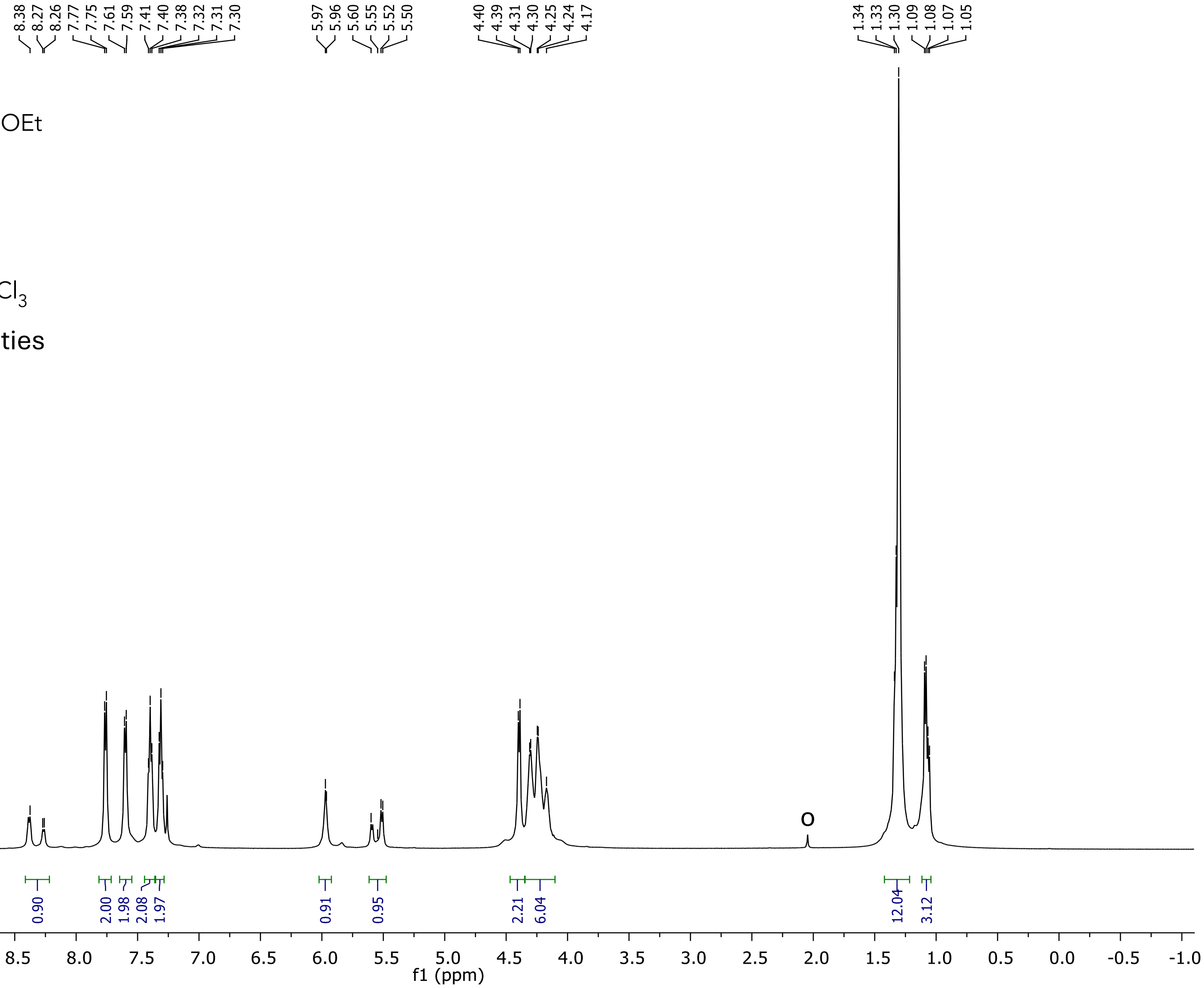

**Ethyl 2-((2S,3R)-2-((((9H-fluoren-9-yl)methoxy)carbonyl)amino)-3-(tert-butoxy)butanamido)-2-hydroxyacetate (2m)**

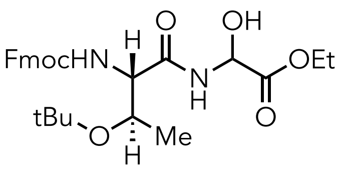


**2m**,
$^{13}$C-NMR, 125 MHz, $CDCl_3$

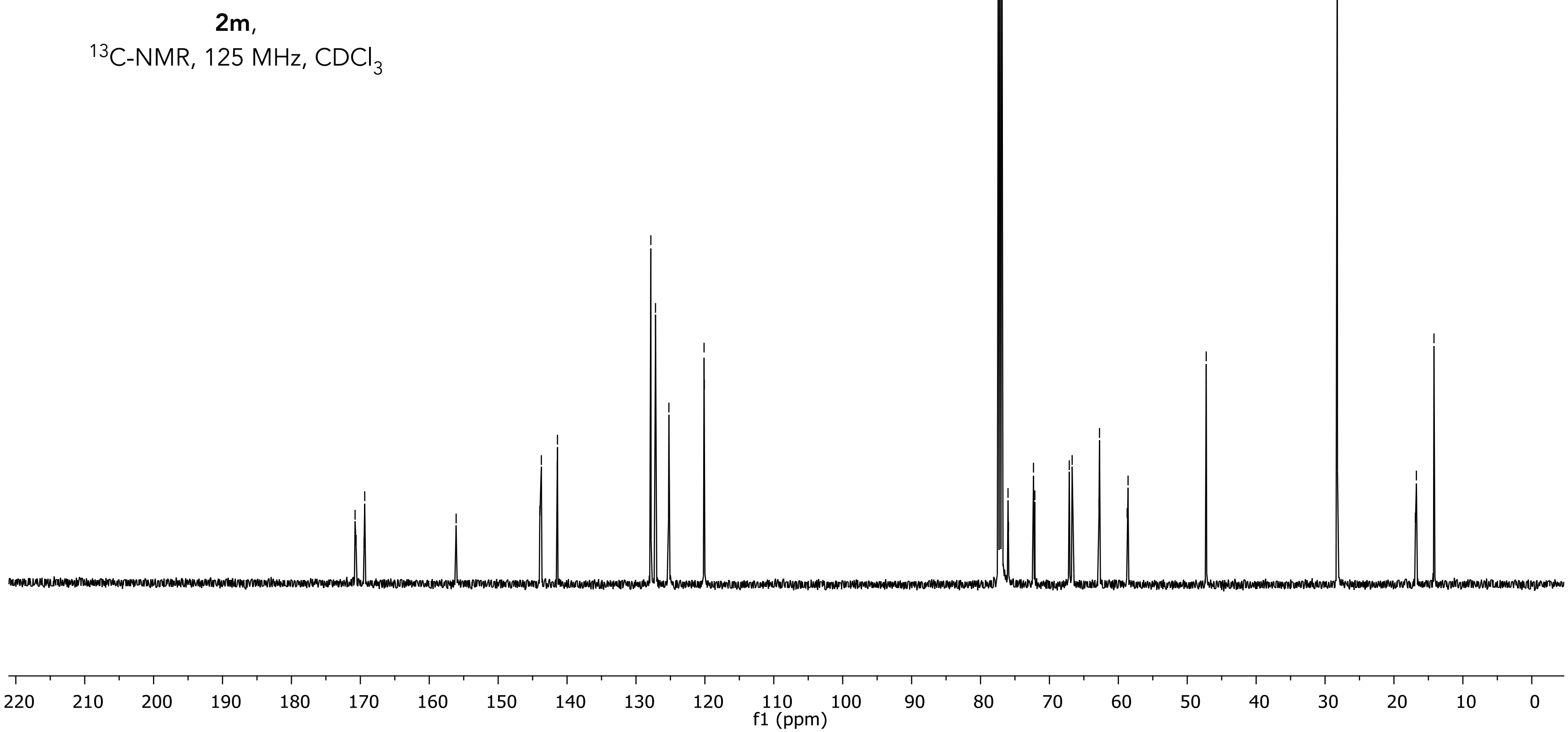

### Ethyl 2-((R)-2-((((9H-fluoren-9-yl)methoxy)carbonyl)amino)-3-(tert-butylthio)propanamido)-2-hydroxyacetate (2n)

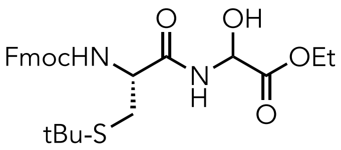


**2n**,
$^{1}$H-NMR, 500 MHz, Acetone-$D_6$

O = Unknown impurities

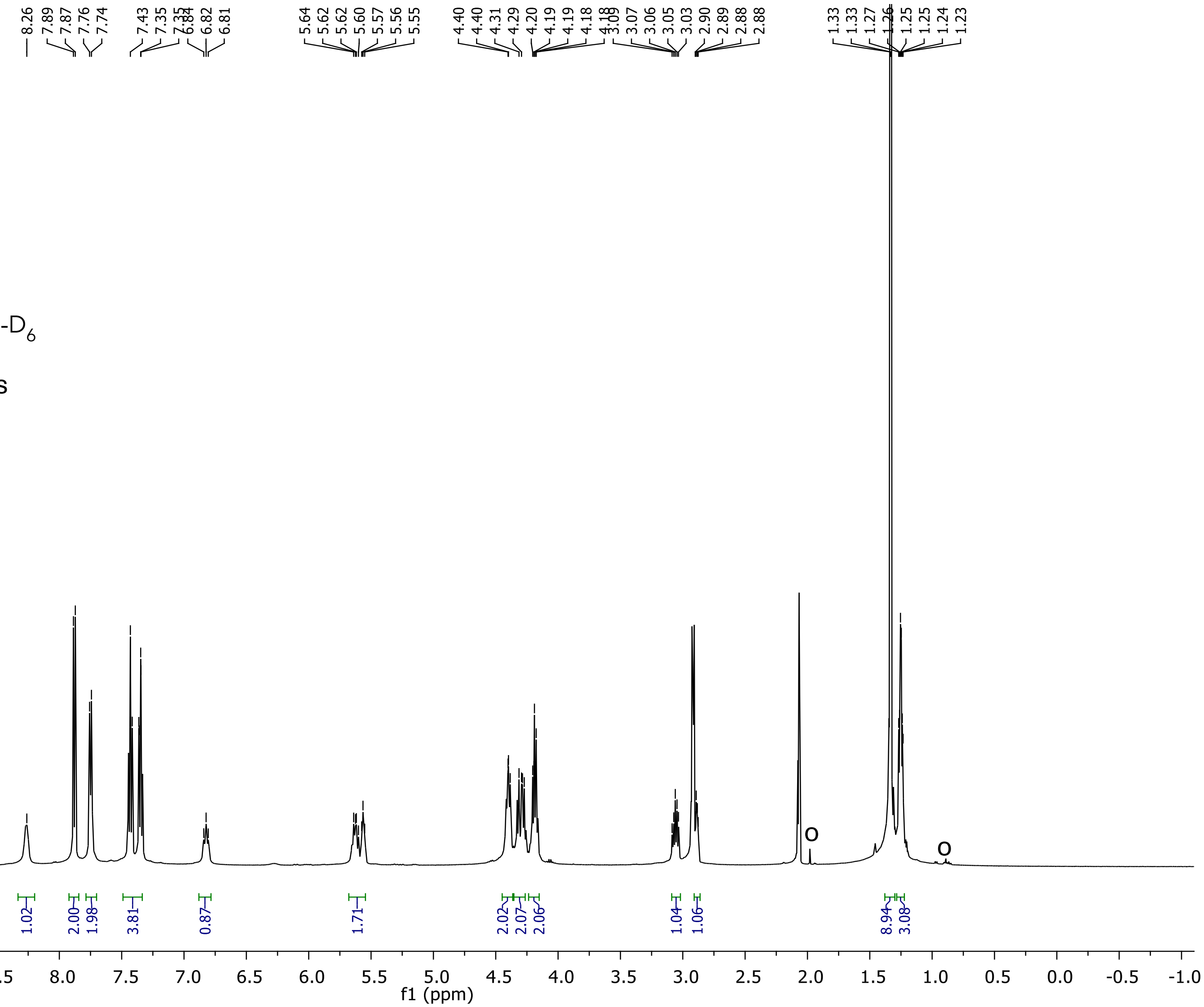

**Ethyl 2-((R)-2-((((9H-fluoren-9-yl)methoxy)carbonyl)amino)-3-(tert-butylthio)propanamido)-2-hydroxyacetate (2n)**

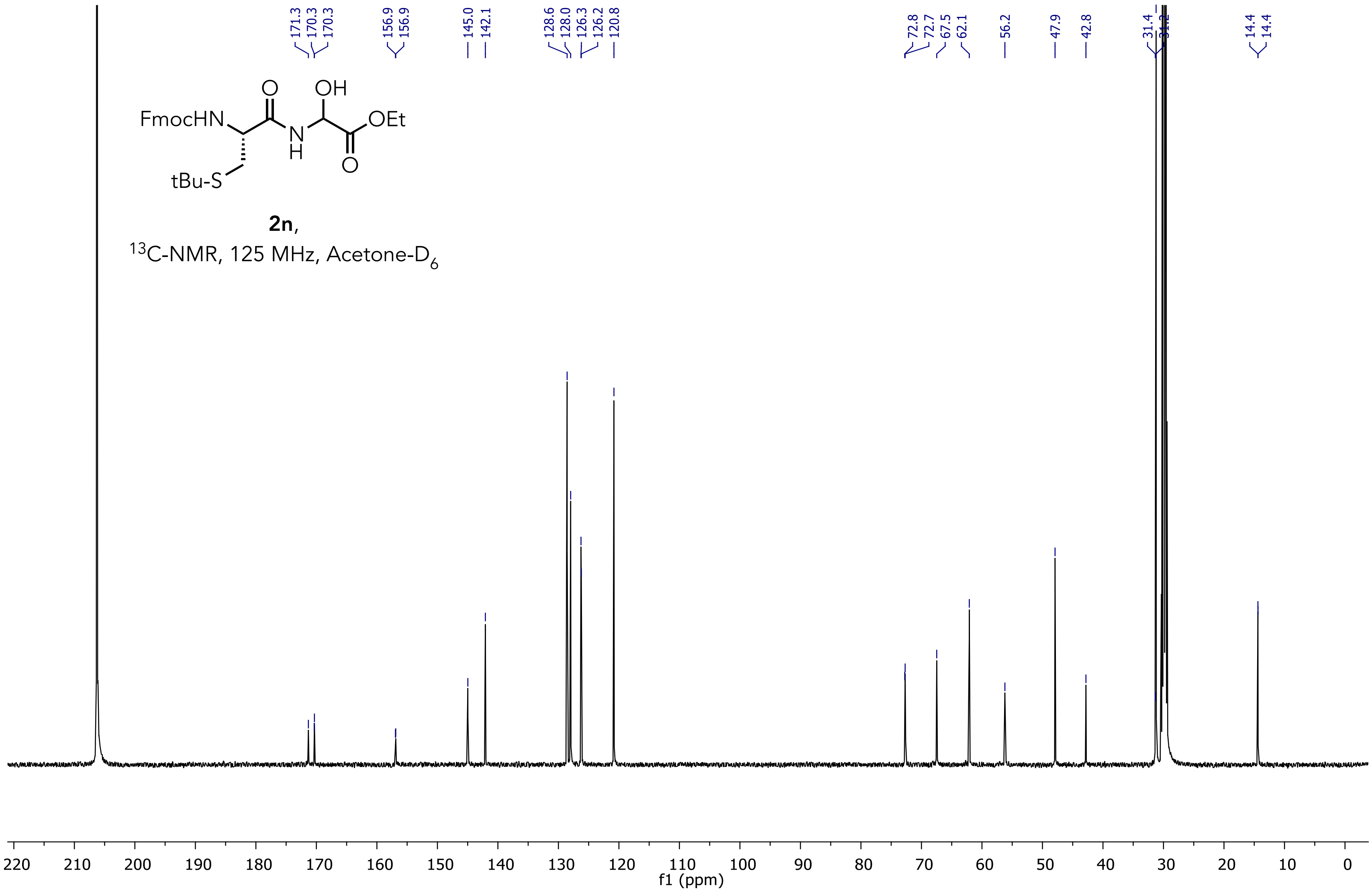

**Tert-butyl (3S)-3-((((9H-fluoren-9-yl)methoxy)carbonyl)amino)-4-((2-ethoxy-1-hydroxy-2-oxoethyl)amino)-4-oxobutanoate (2o)**

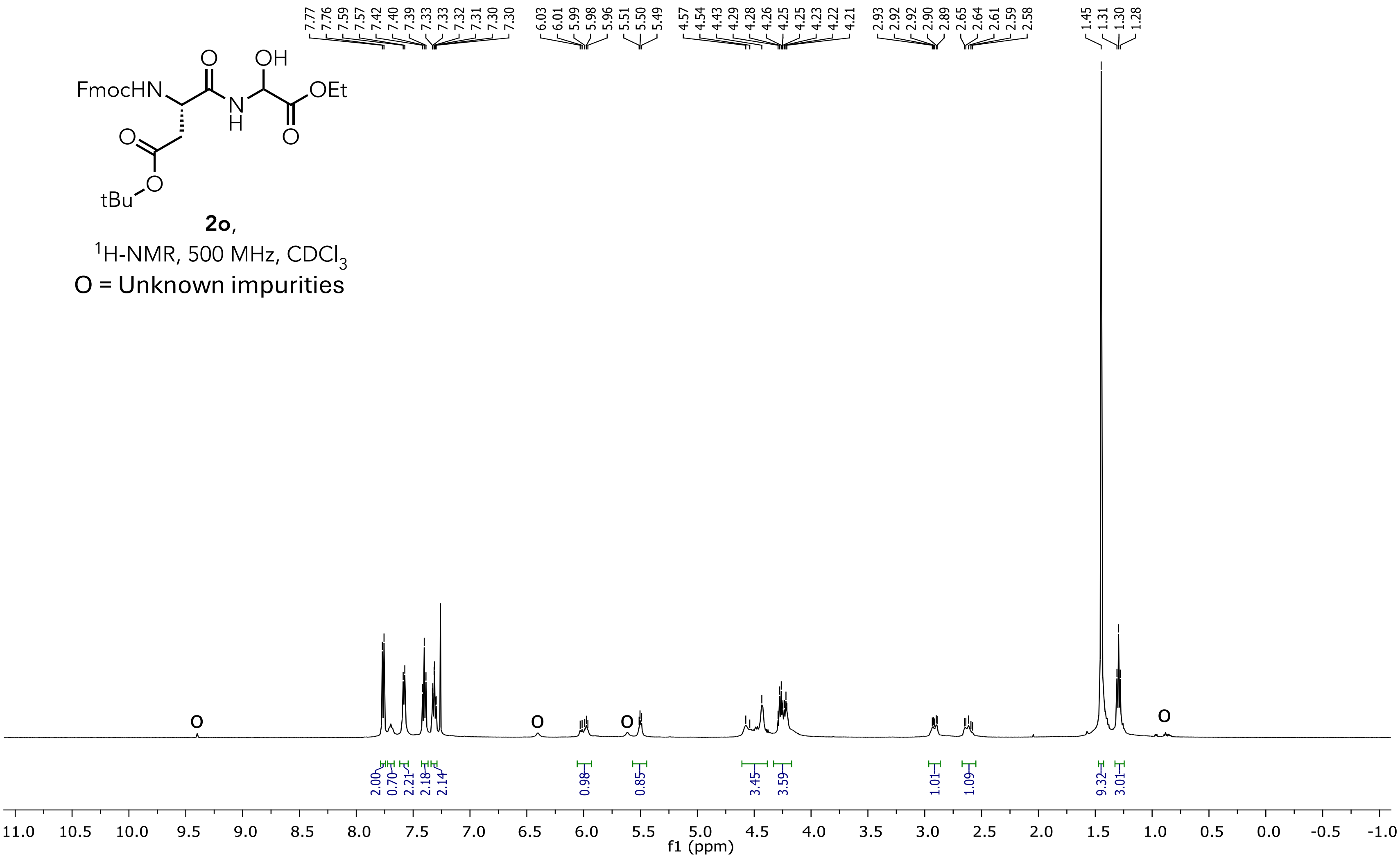

**Tert-butyl (3S)-3-((((9H-fluoren-9-yl)methoxy)carbonyl)amino)-4-((2-ethoxy-1-hydroxy-2-oxoethyl)amino)-4-oxobutanoate (2o)**

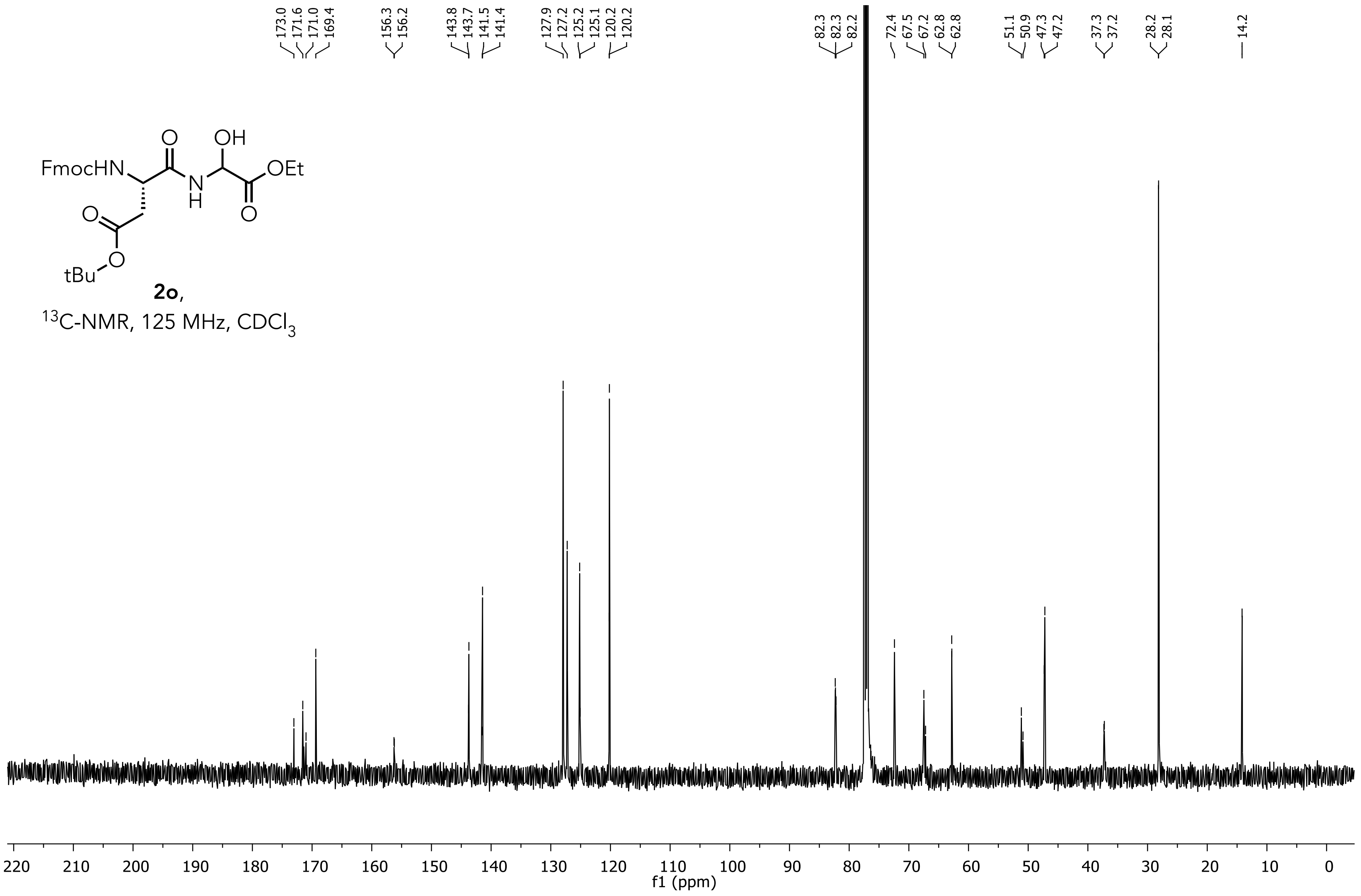

**Ethyl 2-((S)-2-((((9H-fluoren-9-yl)methoxy)carbonyl)amino)-5-oxo-5-(tritylamino)pentanami do)-2-hydroxyacetate (2p)**

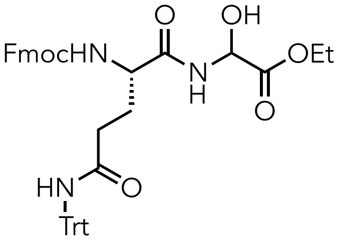

**2p**,

$^1$H-NMR, 500 MHz, $CDCl_3$

O = Unknown impurities

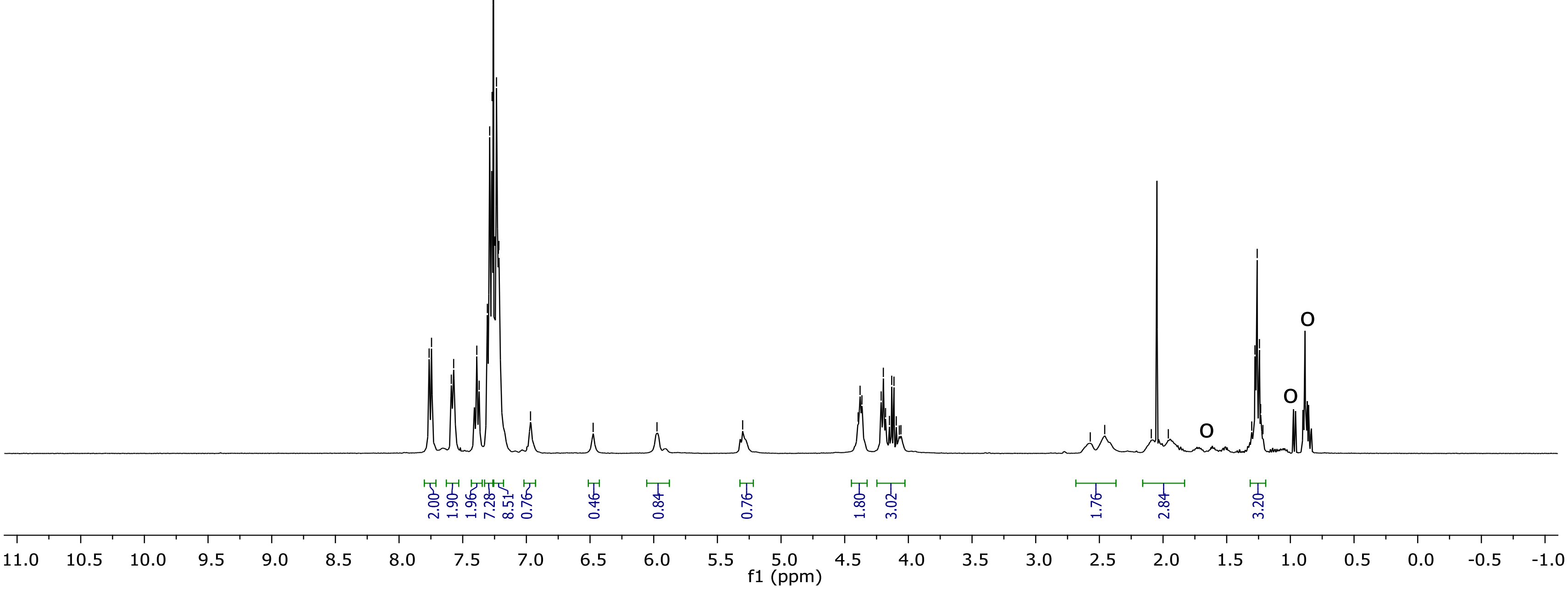

**Ethyl 2-((S)-2-((((9H-fluoren-9-yl)methoxy)carbonyl)amino)-5-oxo-5-(tritylamino)pentanami do)-2-hydroxyacetate (2p)**

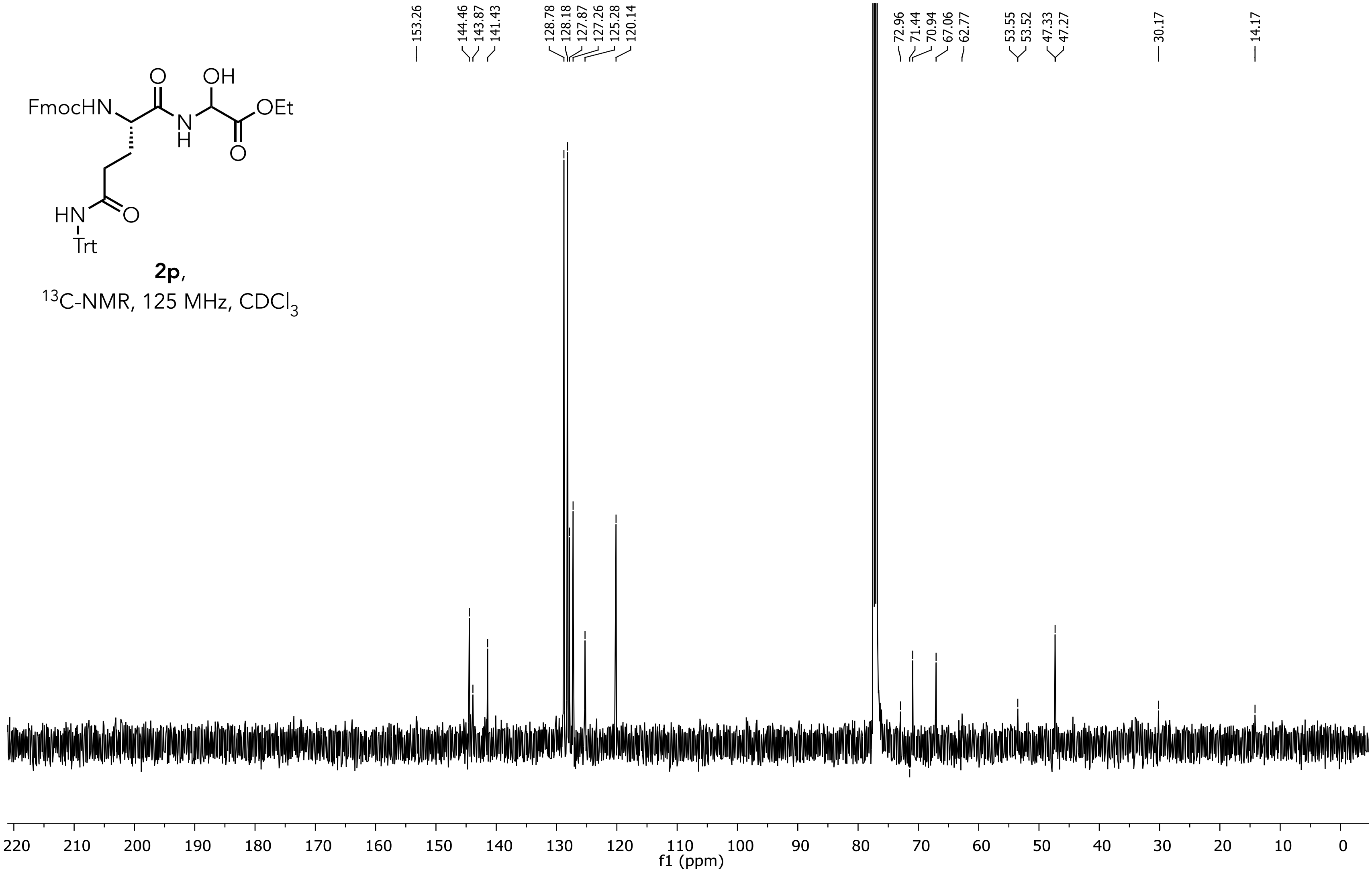


**2p**,
$^{13}$C-NMR, 125 MHz, $CDCl_3$

**Ethyl 2-((S)-2-((((9H-fluoren-9-yl)methoxy)carbonyl)amino)-5-(3-((2,2,4,6,7-pentamethyl-2,3 -dihydrobenzofuran-5-yl)sulfonyl)guanidino)pentanamido)-2-hydroxyacetate (2q)**

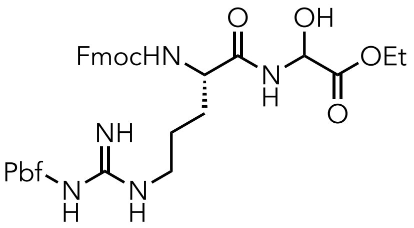


**2q**,
$^{1}$H-NMR, 500 MHz, $CDCl_3$
O = Unknown impurities

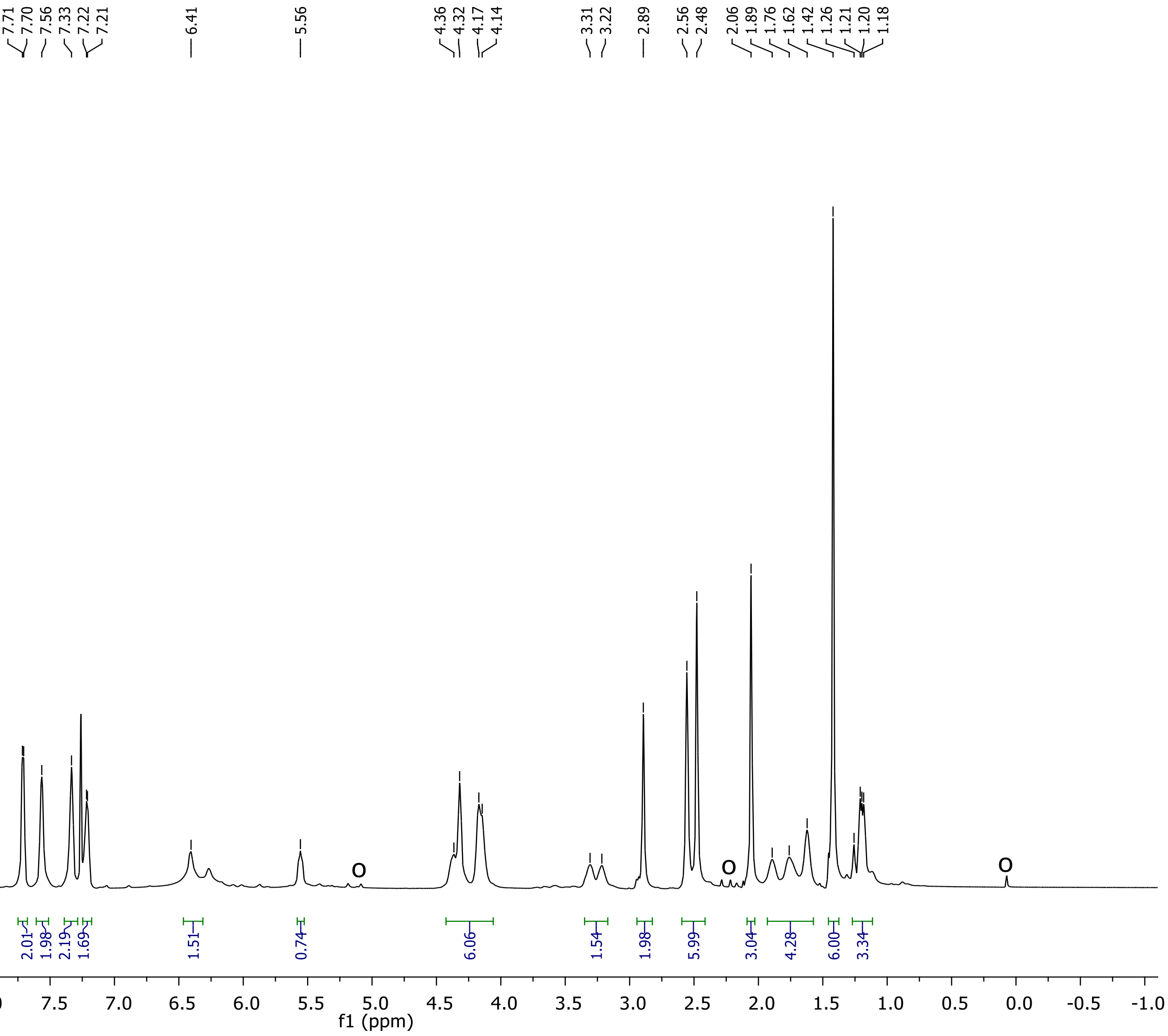

**Ethyl 2-((S)-2-((((9H-fluoren-9-yl)methoxy)carbonyl)amino)-5-(3-((2,2,4,6,7-pentamethyl-2,3 -dihydrobenzofuran-5-yl)sulfon yl)guanidino)pentanamido)-2-hydroxyacetate (2q)**

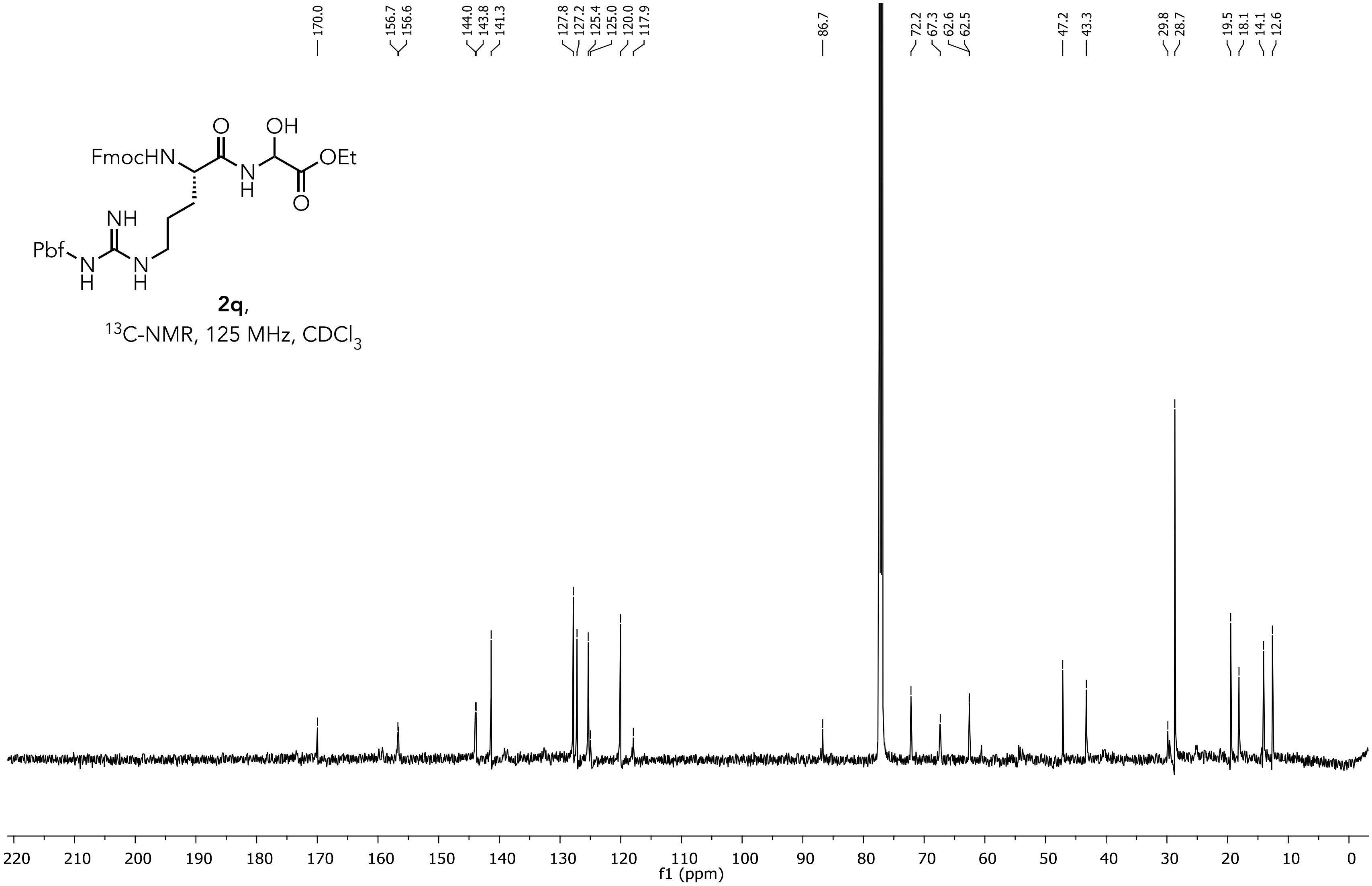


**2q**,
$^{13}$C-NMR, 125 MHz, $CDCl_3$

**Ethyl 2-((S)-2-((((9H-fluoren-9-yl)methoxy)carbonyl)amino)propanamido)-2-(1H-indol-3-yl)acetate (3a)**

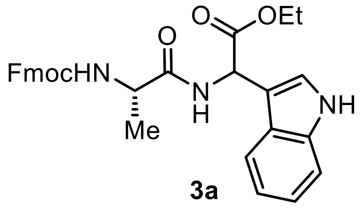


[1]H NMR, 400 MHz, $CDCl_3$

O = Unknown impurities

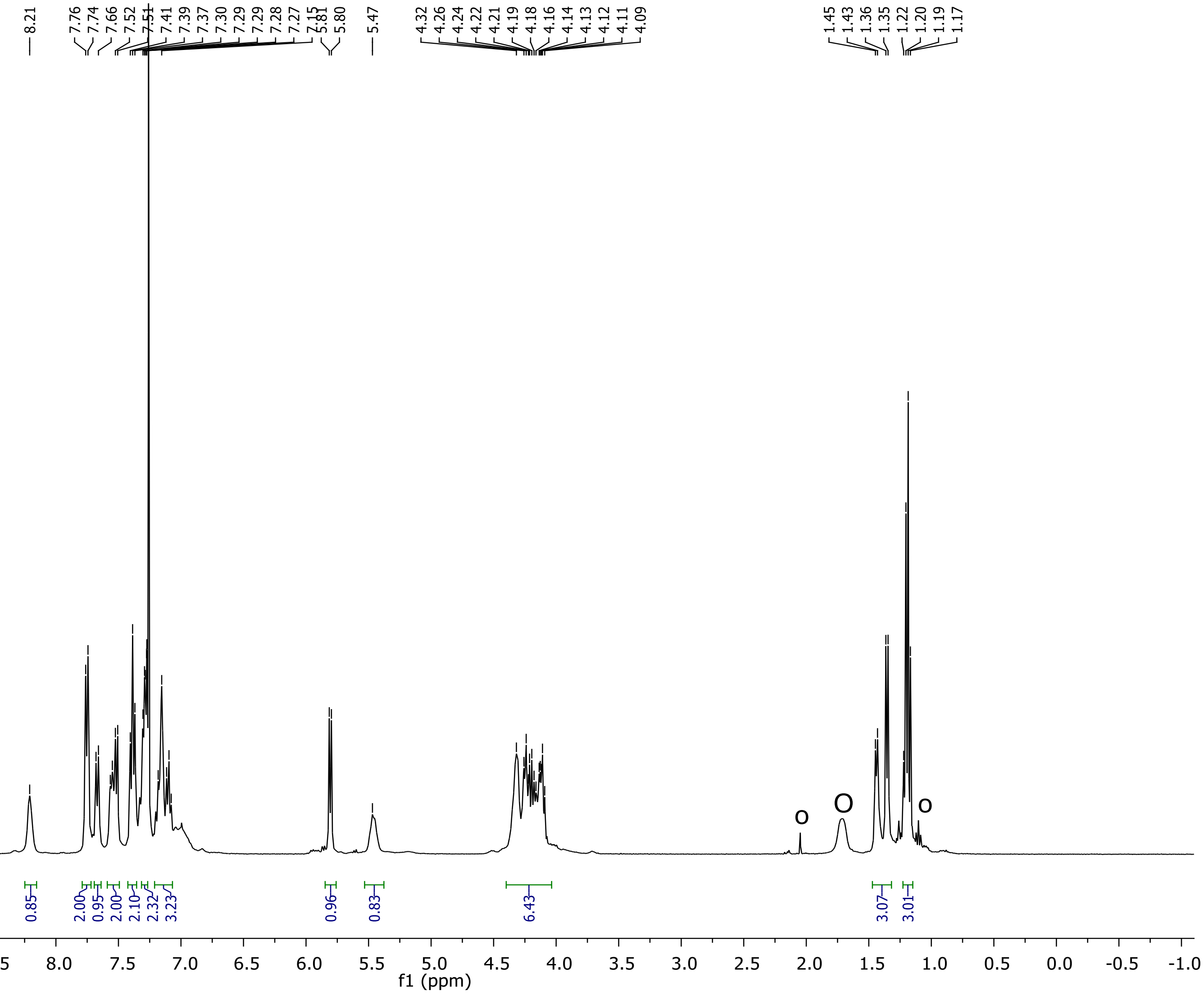

**Ethyl 2-((S)-2-((((9H-fluoren-9-yl)methoxy)carbonyl)amino)propanamido)-2-(1H-indol-3-yl)acetate (3a)**

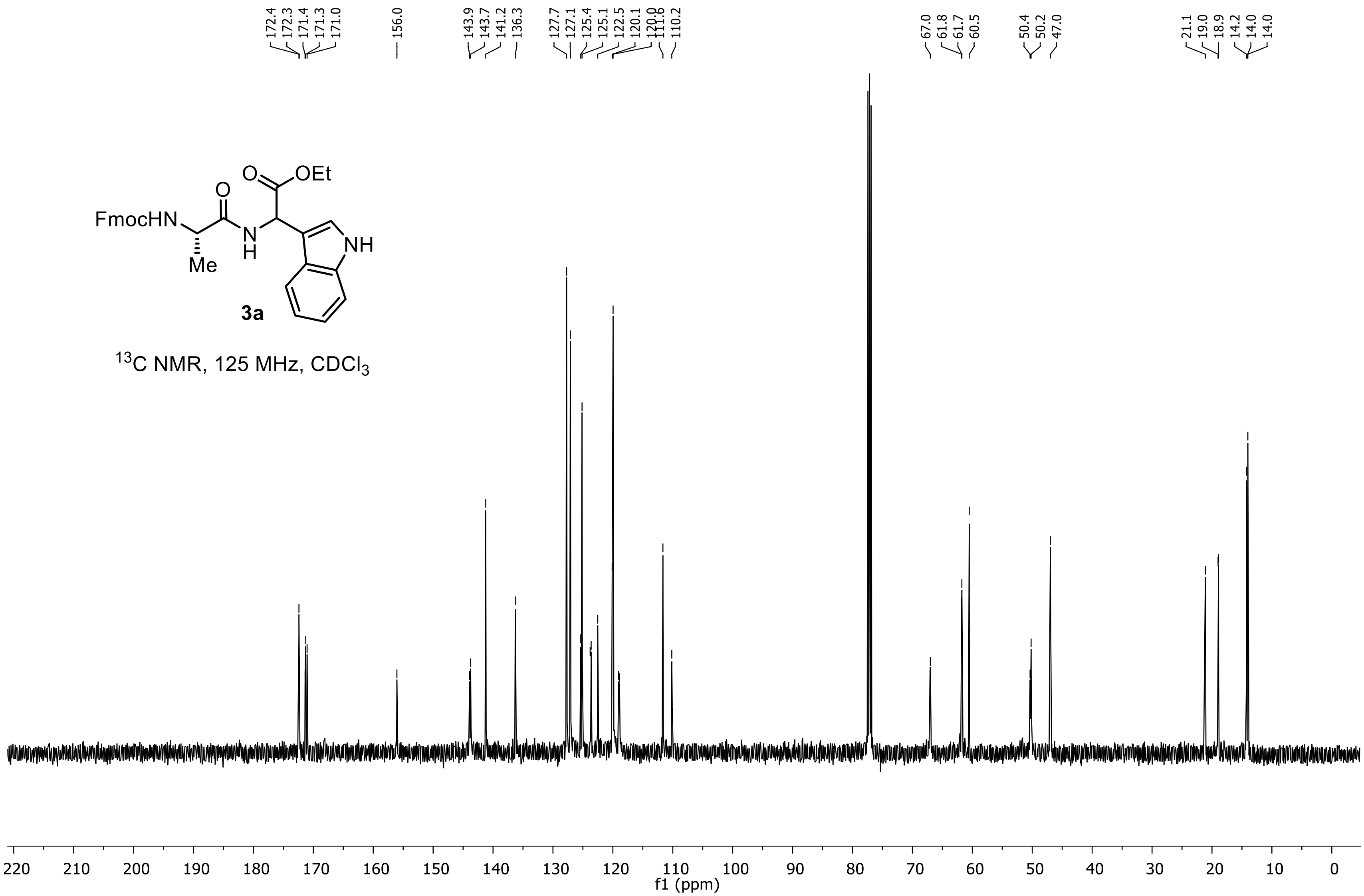

### 2-((*S*)-2-((((9*H*-fluoren-9-yl)methoxy)carbonyl)amino)propanamido)-2-(1*H*-indol-3-yl)acetic acid (3a’)

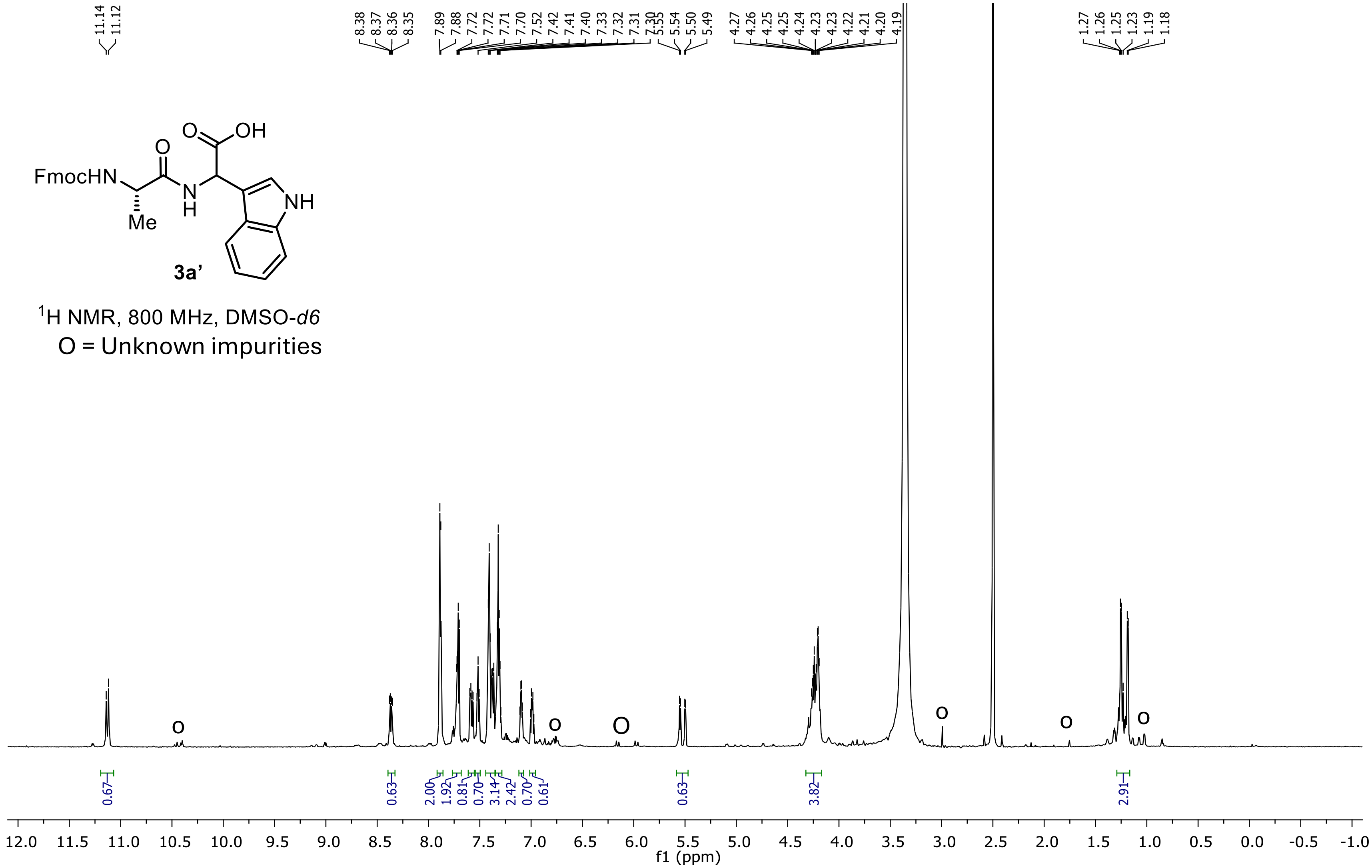

### 2-((*S*)-2-((((9*H*-fluoren-9-yl)methoxy)carbonyl)amino)propanamido)-2-(1*H*-indol-3-yl)acetic acid (3a')

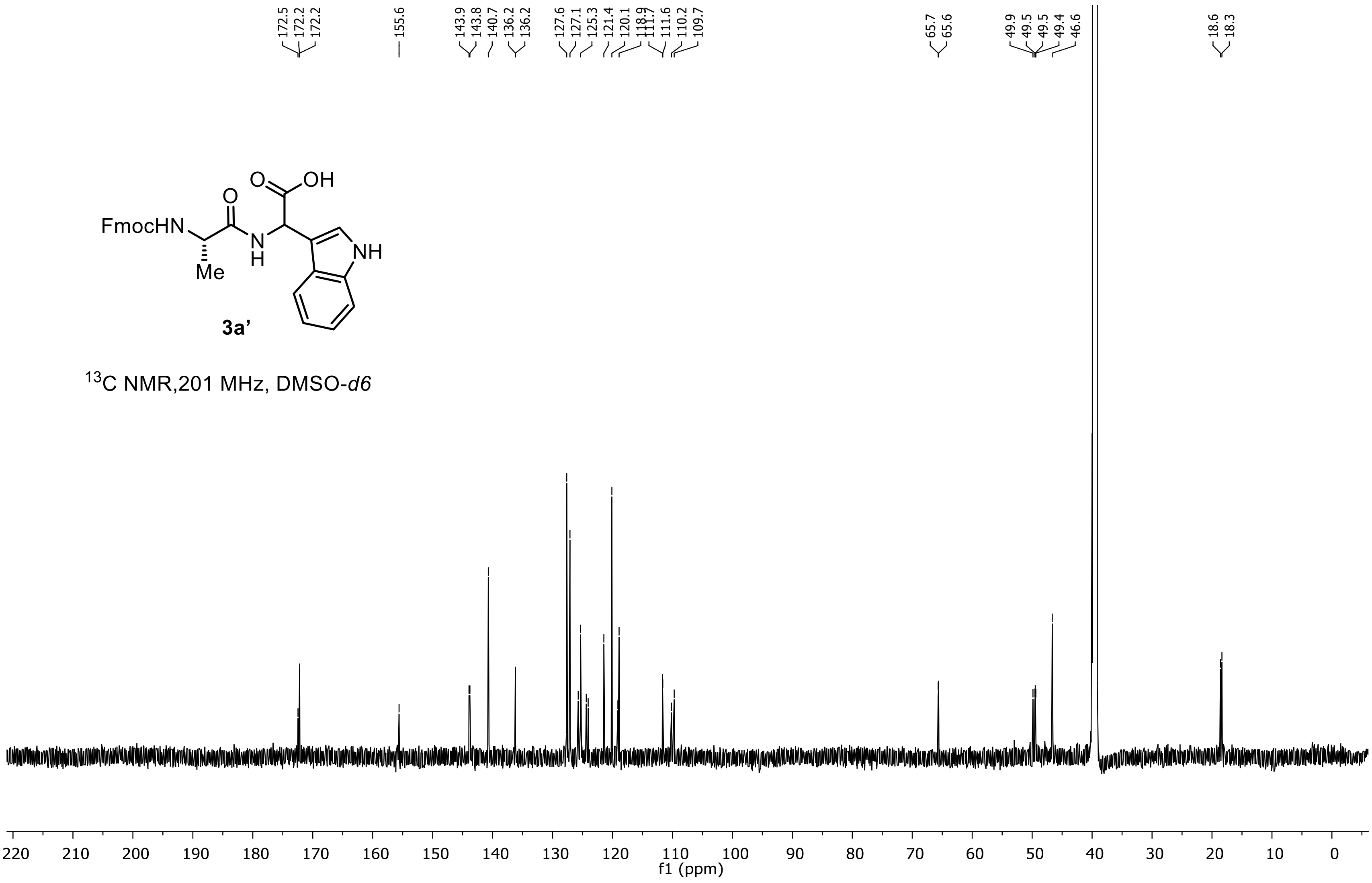


**3a'**

$^{13}$C NMR,201 MHz, DMSO-*d6*

**Ethyl 2-((S)-2-((((9H-fluoren-9-yl)methoxy)carbonyl)amino)propanamido)-2-(6-cyano-1H-indol-3-yl)acetate (3b)**

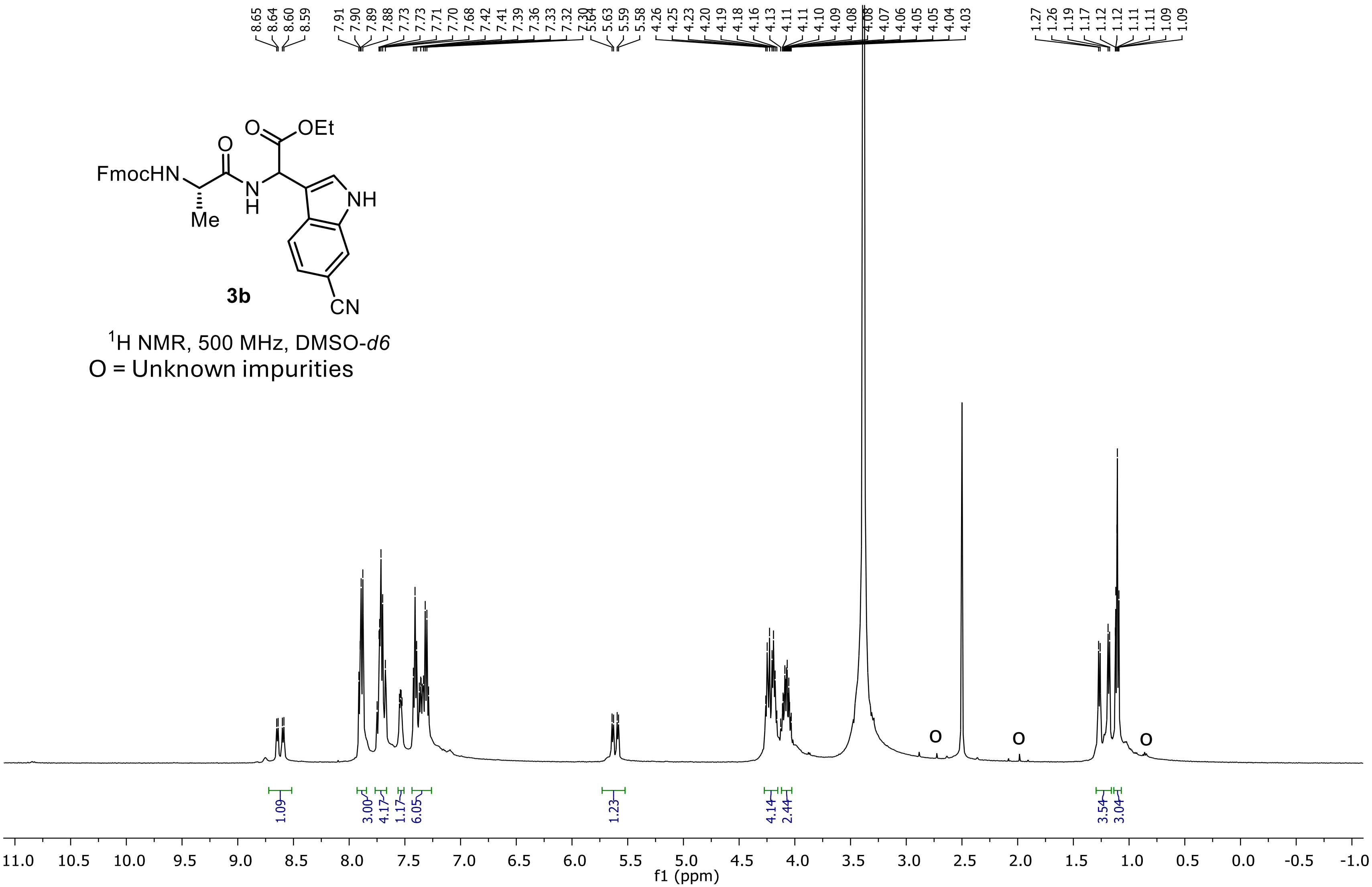

**Ethyl 2-((S)-2-((((9H-fluoren-9-yl)methoxy)carbonyl)amino)propanamido)-2-(6-cyano-1H-indol-3-yl)acetate (3b)**

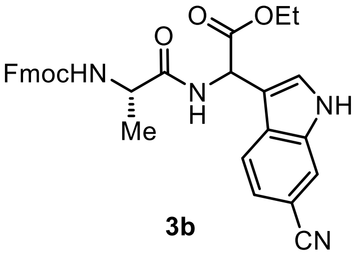


**3b**

$^{13}C$ NMR,125 MHz, DMSO-*d6*

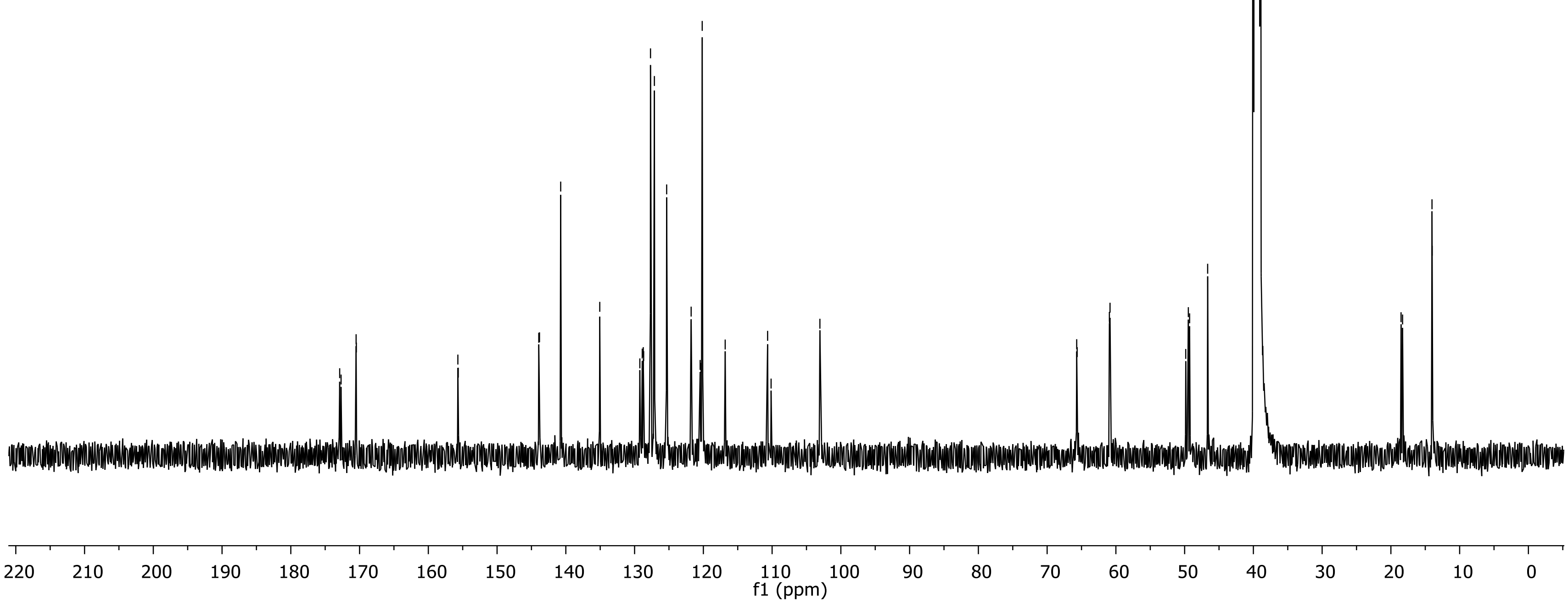

**Ethyl 2-((S)-2-((((9H-fluoren-9-yl)methoxy)carbonyl)amino)propanamido)-2-(4-cyano-1H-indol-3-yl)acetate (3c)**

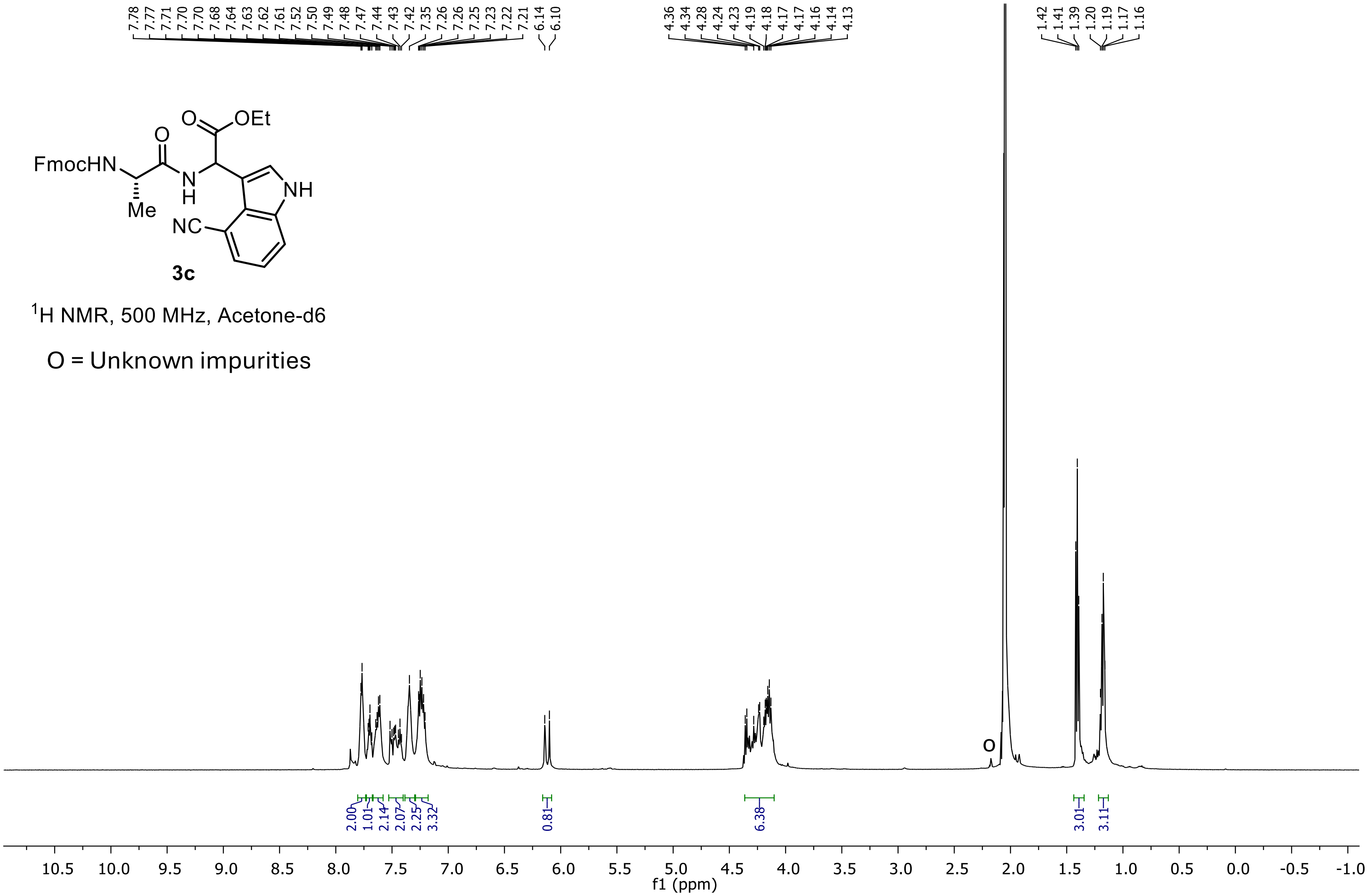

**Ethyl 2-((S)-2-((((9H-fluoren-9-yl)methoxy)carbonyl)amino)propanamido)-2-(4-cyano-1H-indol-3-yl)acetate (3c)**

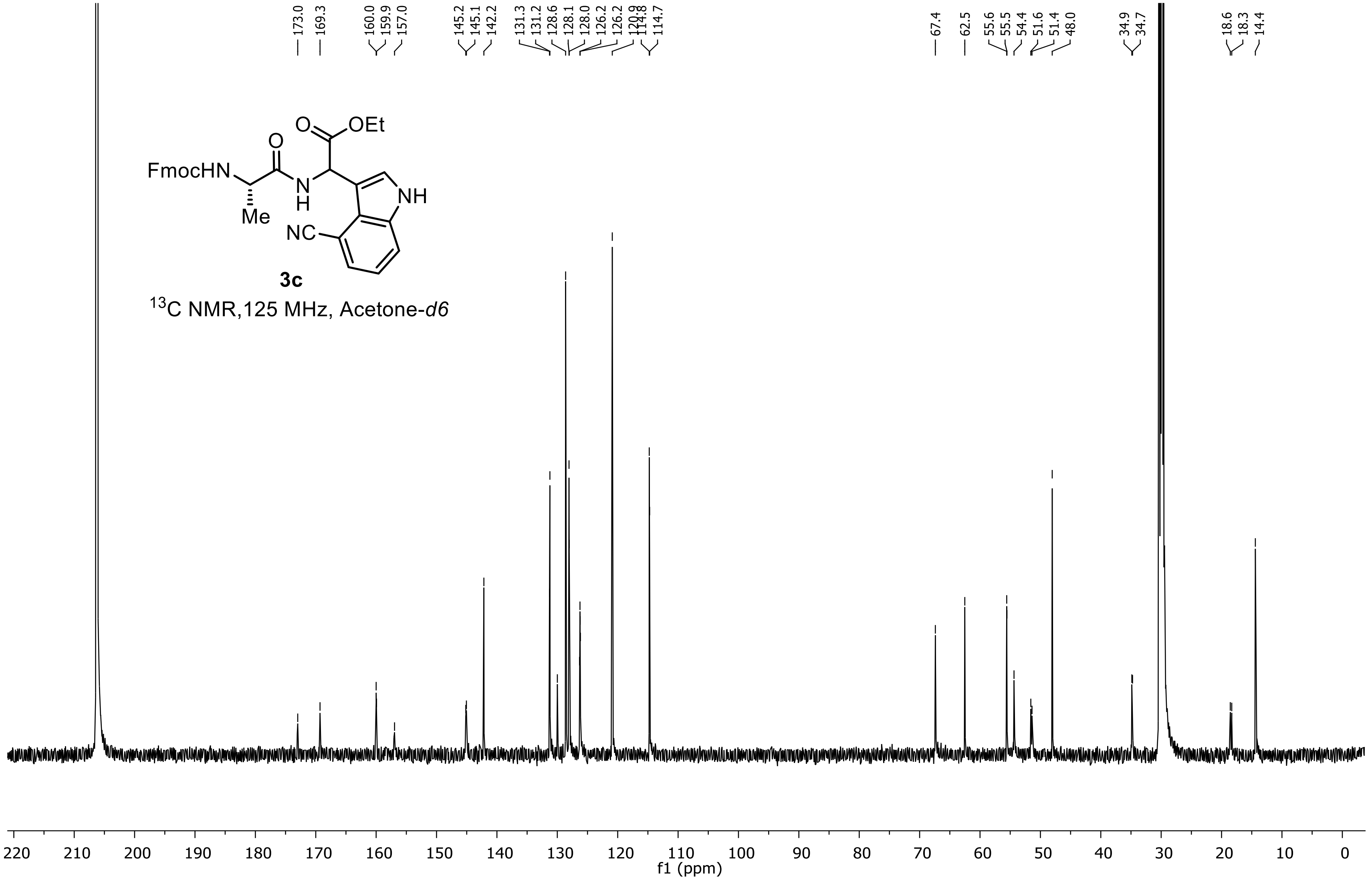

Ethyl 2-((S)-2-((((9H-fluoren-9-yl)methoxy)carbonyl)amino)propanamido)-2-(2,4,6-trimethoxyphenyl)acetate (3d)

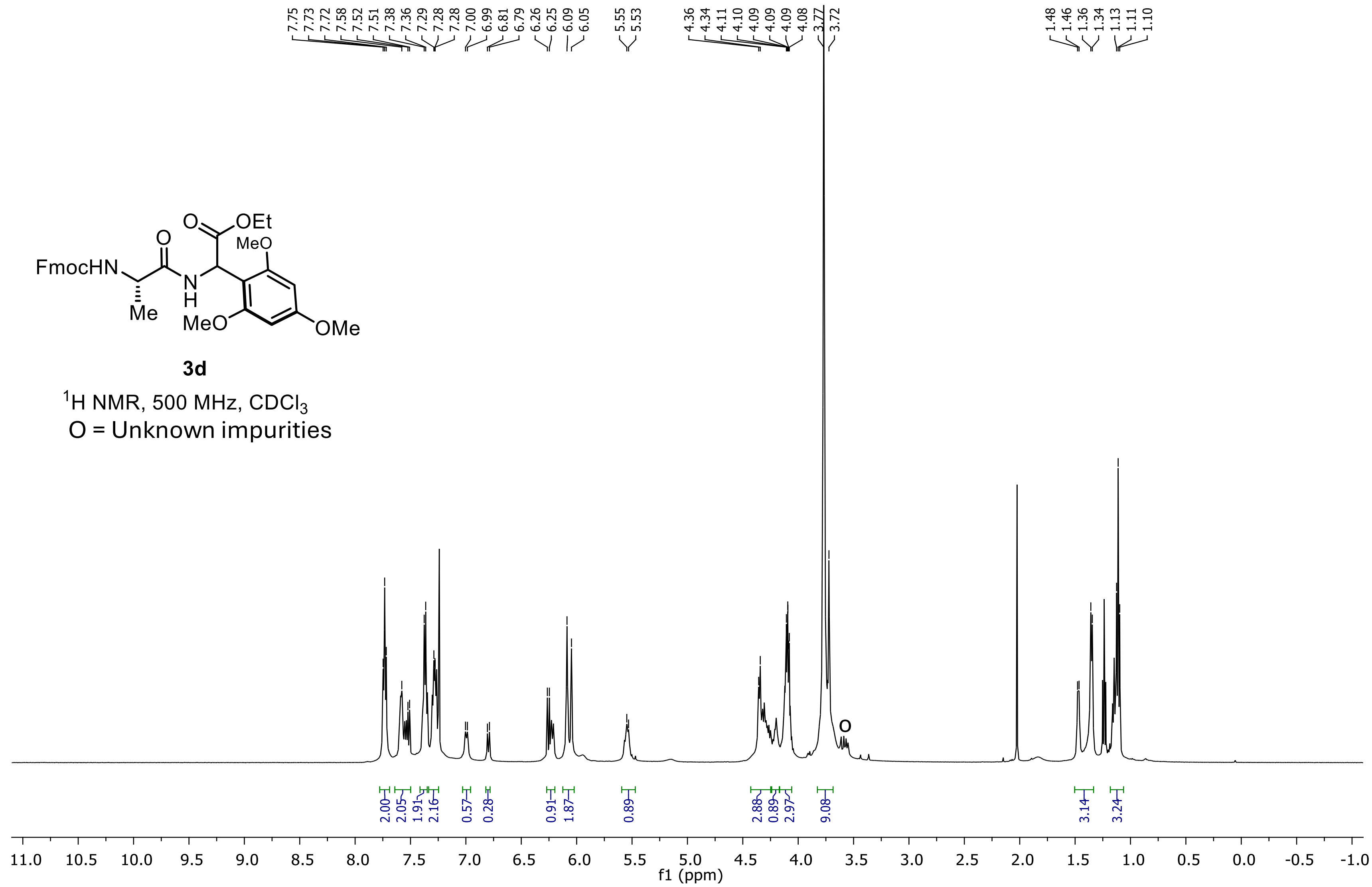

**Ethyl 2-((S)-2-((((9H-fluoren-9-yl)methoxy)carbonyl)amino)propanamido)-2-(2,4,6-trimethoxyphenyl)acetate (3d)**

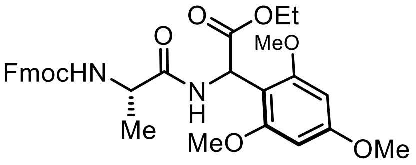


**3d**

$^{13}C$ NMR,125 MHz, $CDCl_3$

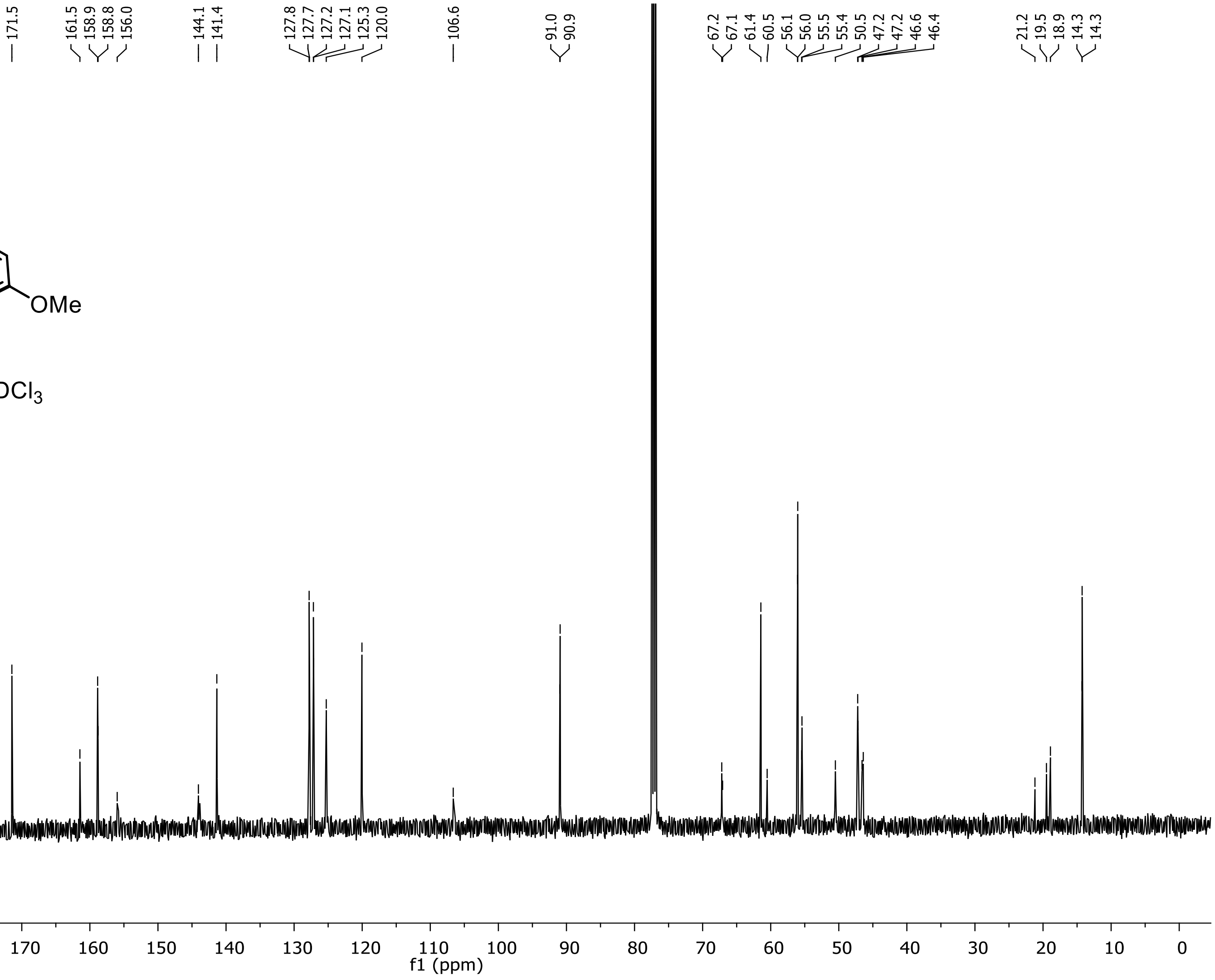

**Ethyl 2-((S)-2-((((9H-fluoren-9-yl)methoxy)carbonyl)amino)propanamido)-2-(4-phenylbutoxy)acetate (4a)**

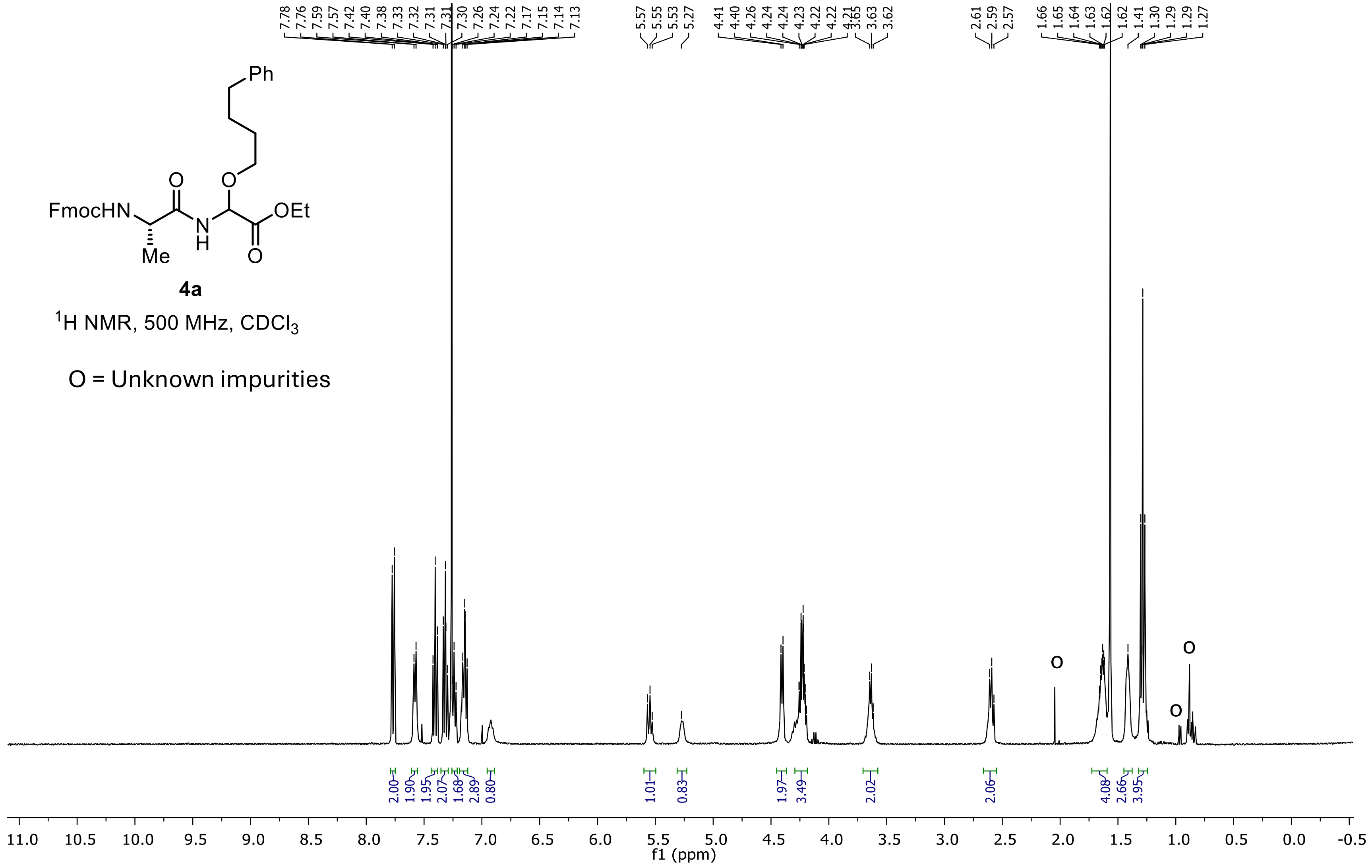

**Ethyl 2-((S)-2-((((9H-fluoren-9-yl)methoxy)carbonyl)amino)propanamido)-2-(4-phenylbutoxy)acetate (4a)**

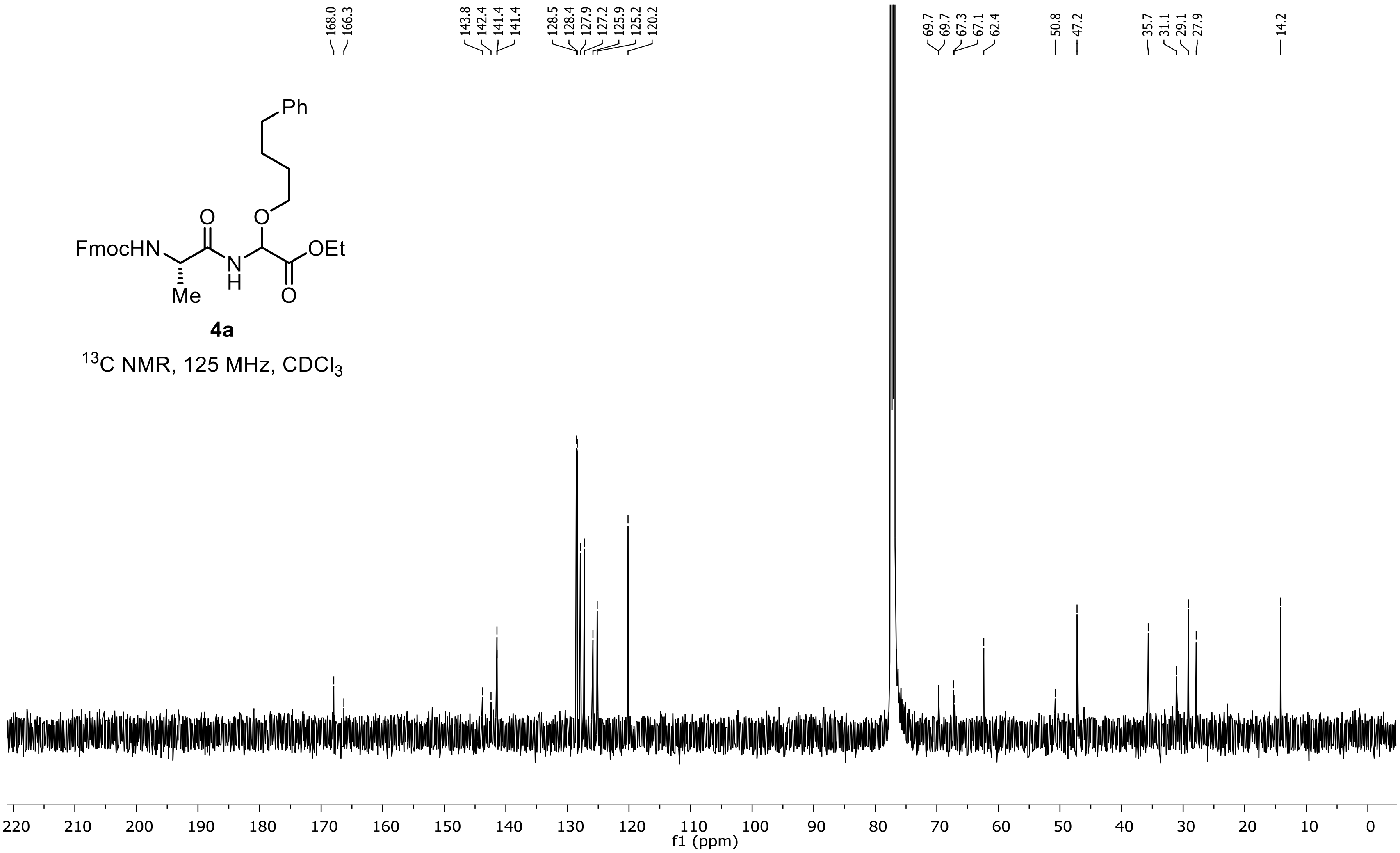

### (9H-fluoren-9-yl)methyl ((1S)-1-(5-oxo-4-(4-phenylbutoxy)-4,5-dihydrooxazol-2-yl)ethyl)carbamate (4a’)

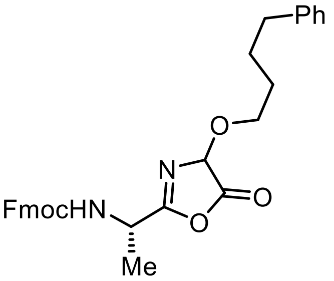


**4a'**

$^{1}$H NMR, 500 MHz, $CDCl_3$

O = Unknown impurities

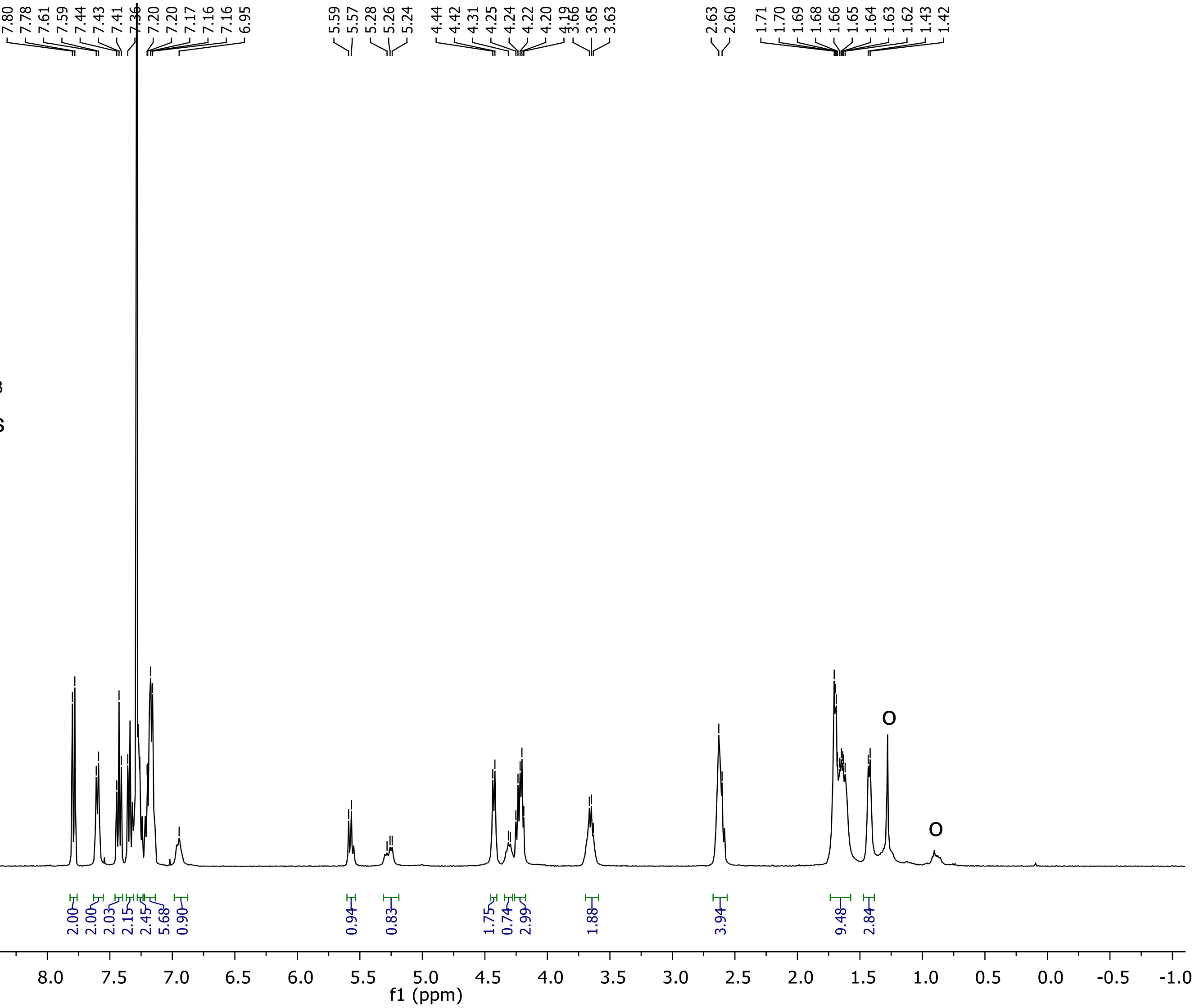

**(9H-fluoren-9-yl)methyl ((1S)-1-(5-oxo-4-(4-phenylbutoxy)-4,5-dihydrooxazol-2-yl)ethyl)carbamate (4a')**

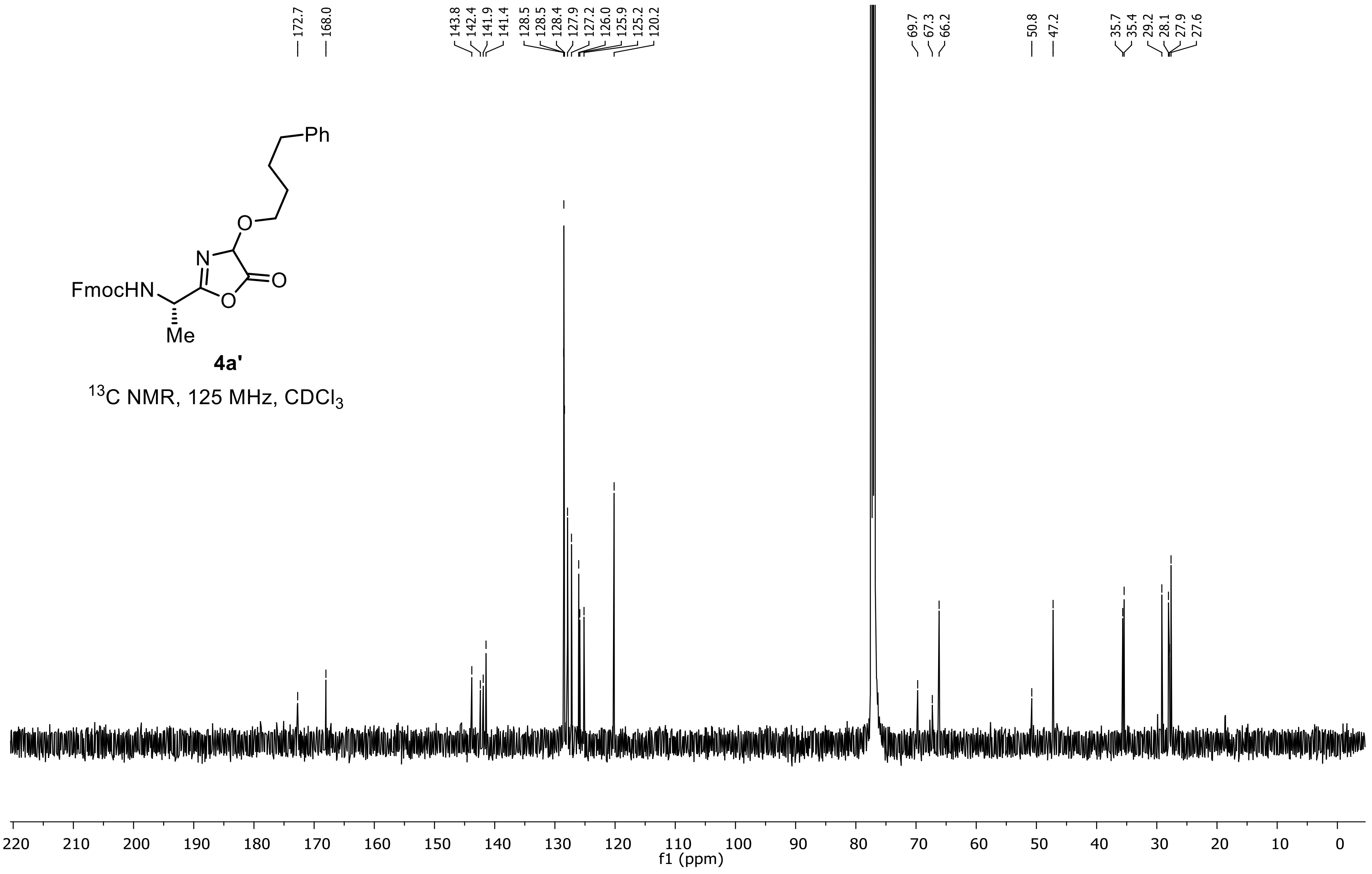

**Ethyl 2-((S)-2-((((9H-fluoren-9-yl)methoxy)carbonyl)amino)propanamido)-2-((4-(trifluoromethyl)benzyl)thio)acetate (4b)**

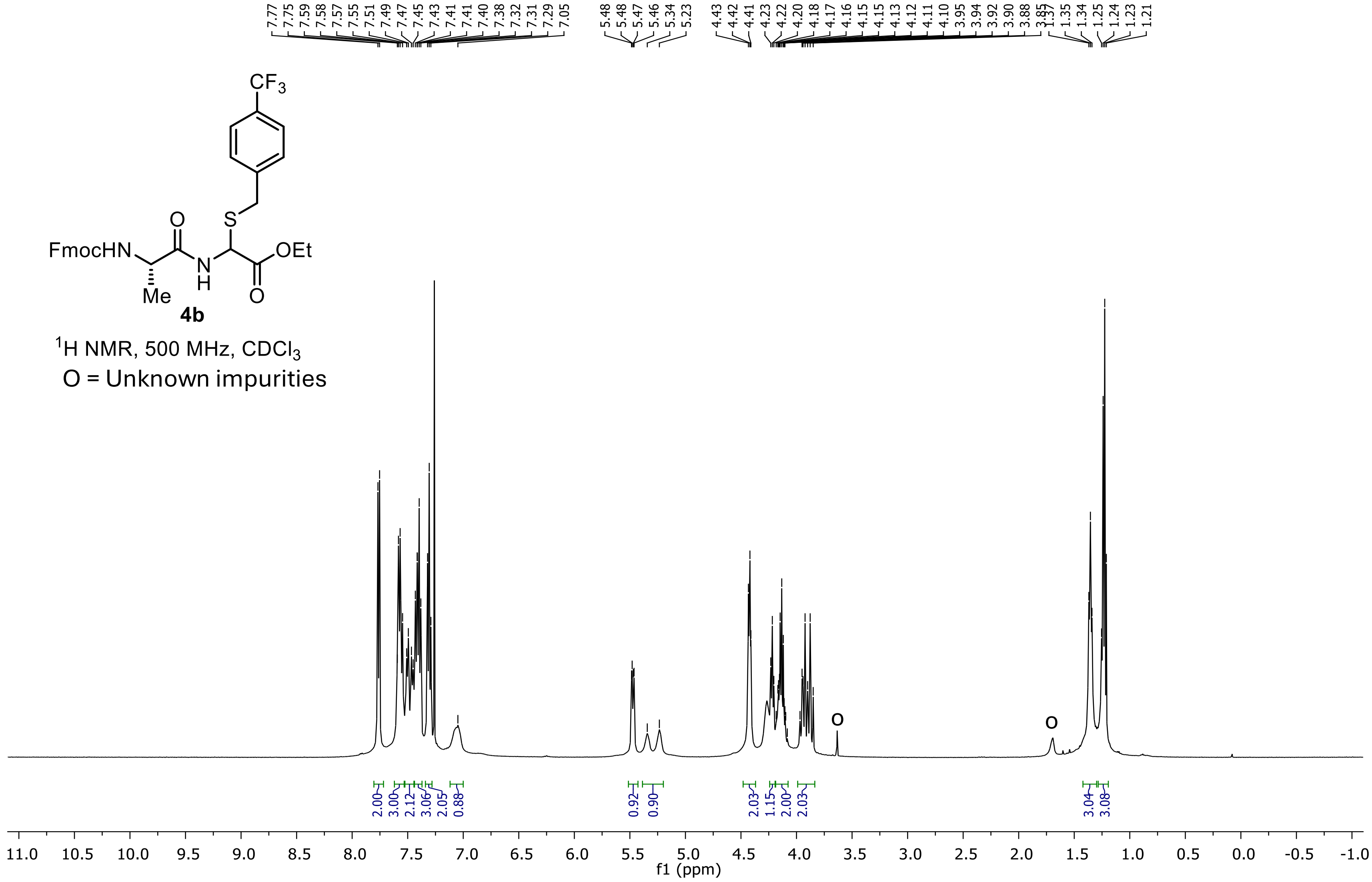

**Ethyl 2-((S)-2-((((9H-fluoren-9-yl)methoxy)carbonyl)amino)propanamido)-2-((4-(trifluoromethyl)benzyl)thio)acetate (4b)**

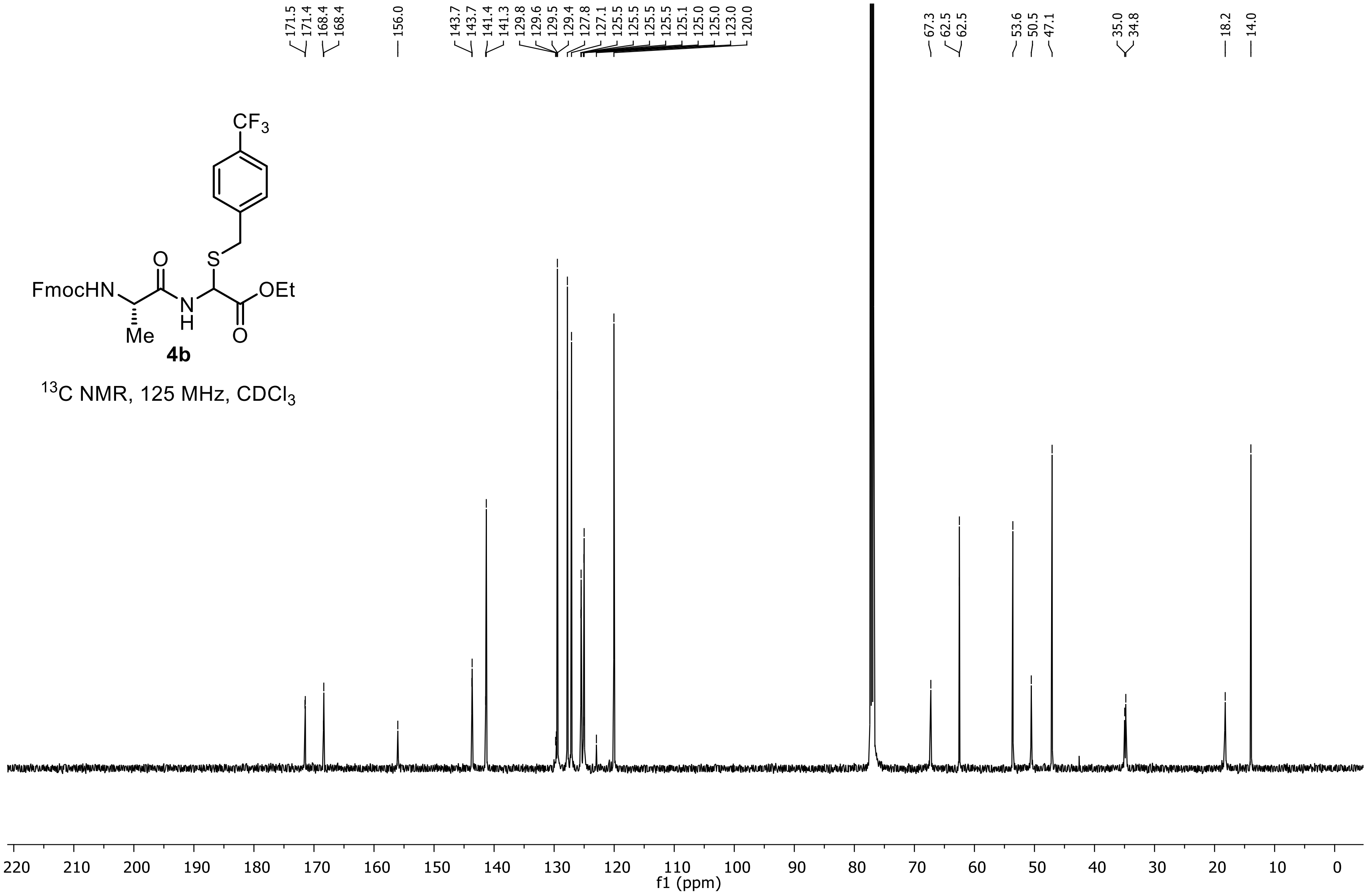

**Ethyl 2-((S)-2-((((9H-fluoren-9-yl)methoxy)carbonyl)amino)propanamido)-2-((4-(trifluoromethyl)benzyl)thio)acetate (4b)**

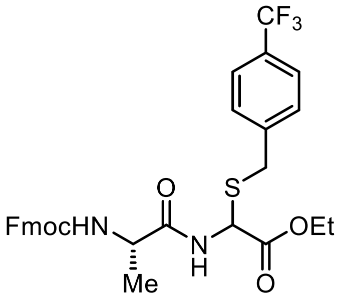


**4b**

$^{19}F$ NMR, 471 MHz, $CDCl_3$

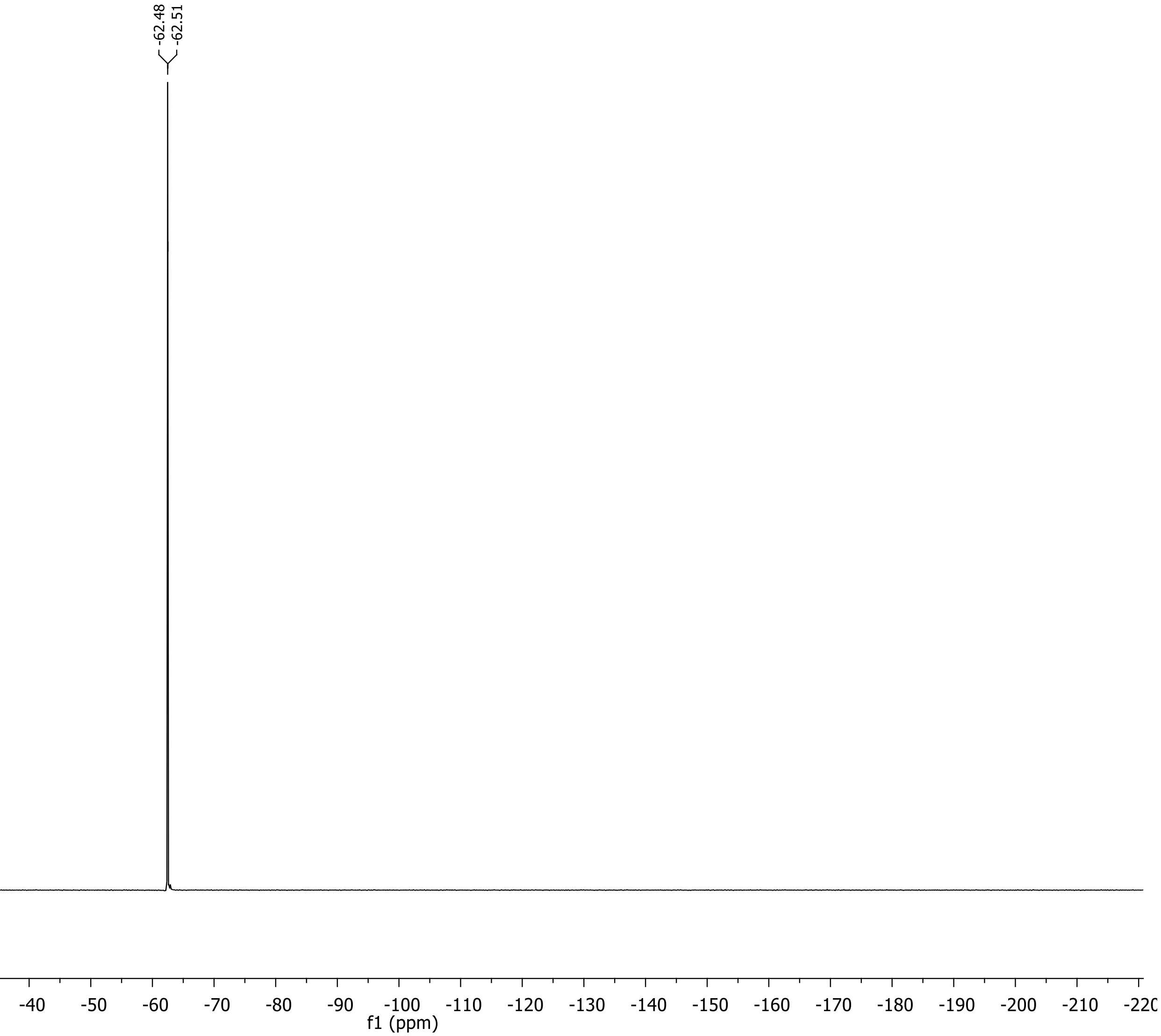

**Ethyl 2-((S)-2-((((9H-fluoren-9-yl)methoxy)carbonyl)amino)propanamido)-2-((4-methoxybenzyl)thio)acetate (4c)**

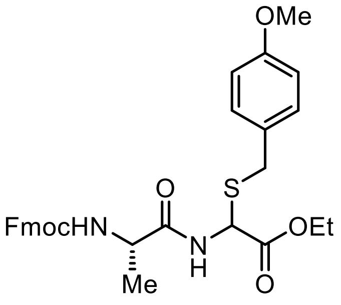


**4c**

$^{1}$H NMR, 500 MHz, Acetone-d6

O = Unknown impurities

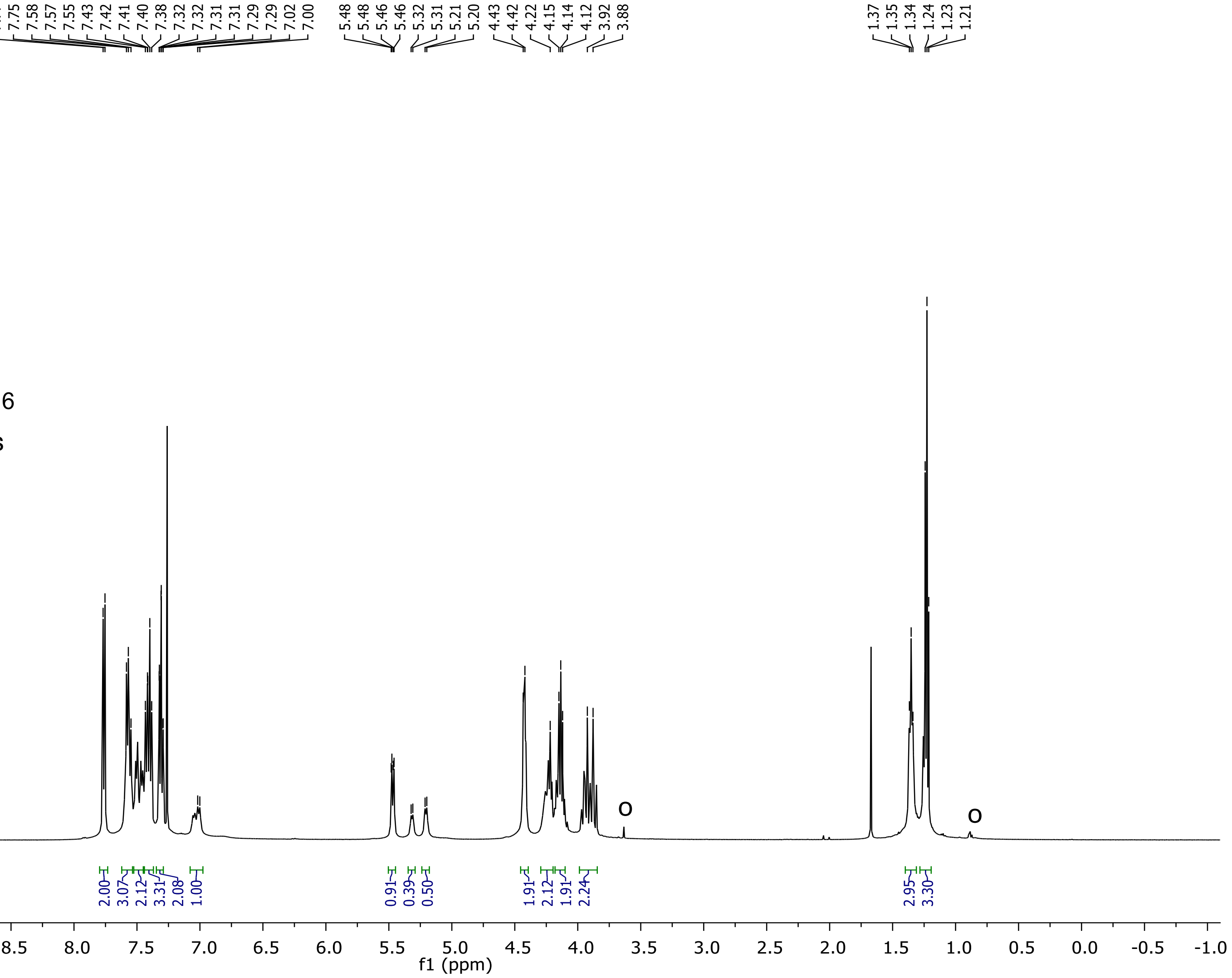

**Ethyl 2-((S)-2-((((9H-fluoren-9-yl)methoxy)carbonyl)amino)propanamido)-2-((4-methoxybenzyl)thio)acetate (4c)**

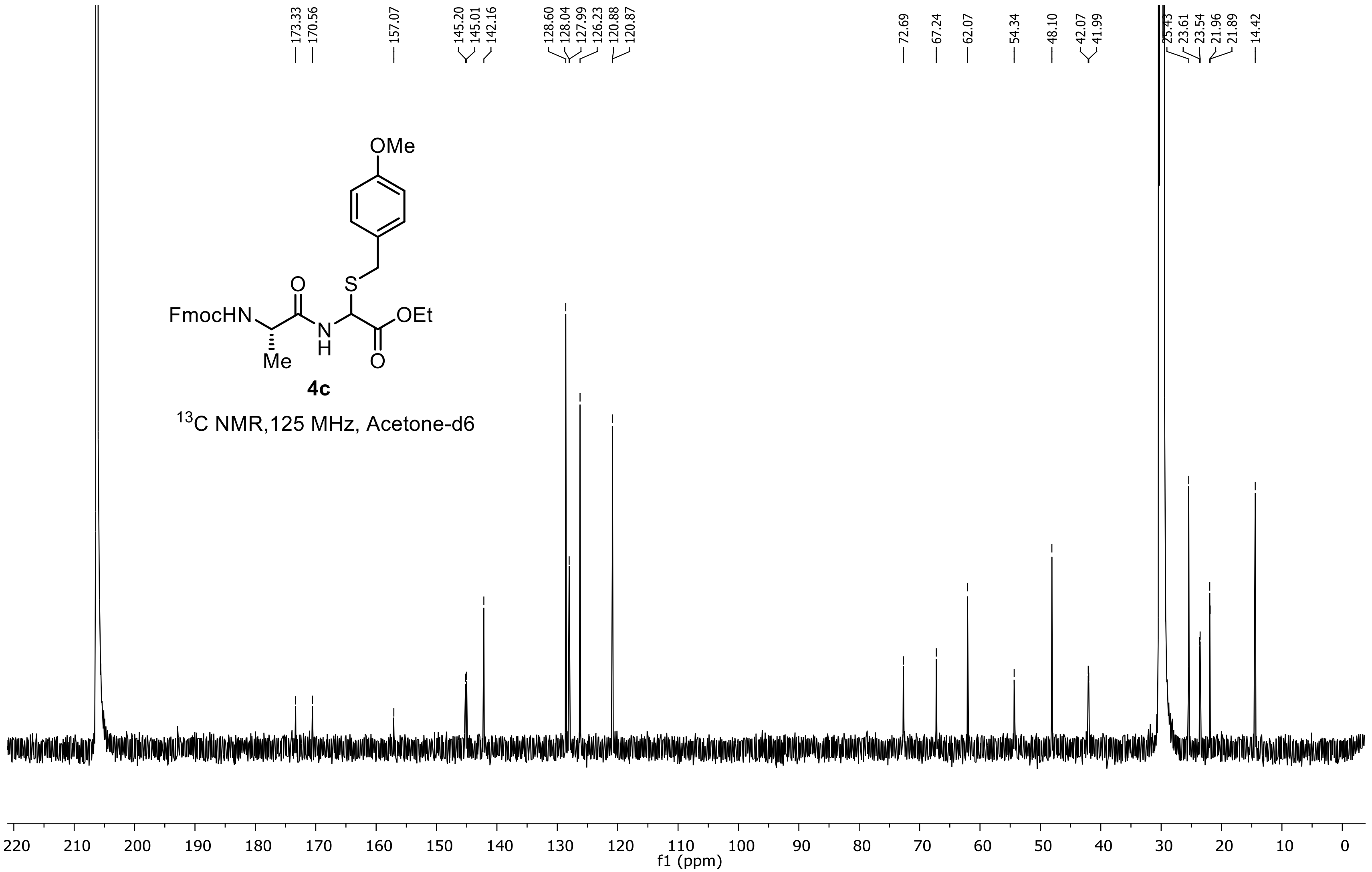
